# Acquisition of discrete immune suppressive barriers contributes to the initiation and progression of preinvasive to invasive human lung cancer

**DOI:** 10.1101/2024.12.31.630523

**Authors:** Liron Yoffe, Bhavneet Bhinder, Sung Wook Kang, Haoran Zhang, Arshdeep Singh, Hiranmayi Ravichandran, Geoffrey Markowitz, Mitchell Martin, Junbum Kim, Chen Zhang, Olivier Elemento, Wesley Tansey, Stewart Bates, Timothy E. McGraw, Alain Borczuk, Hyun-Sung Lee, Nasser K. Altorki, Vivek Mittal

## Abstract

Computerized chest tomography (CT)-guided screening in populations at risk for lung cancer has increased the detection of preinvasive subsolid nodules, which progress to solid invasive adenocarcinoma. Despite the clinical significance, there is a lack of effective therapies for intercepting the progression of preinvasive to invasive adenocarcinoma. To uncover determinants of early disease emergence and progression, we used integrated single-cell approaches, including scRNA-seq, multiplexed imaging mass cytometry and spatial transcriptomics, to construct the first high-resolution map of the composition, lineage/functional states, developmental trajectories and multicellular crosstalk networks from microdissected non-solid (preinvasive) and solid compartments (invasive) of individual part-solid nodules. We found that early disease initiation and subsequent progression are associated with the evolution of immune-suppressive cellular phenotypes characterized by decreased cytotoxic CD8 T and NK cells, increased T cell exhaustion and accumulation of immunosuppressive regulatory T cells (Tregs) and M2-like macrophages expressing TREM2. Within Tregs, we identified a unique population of 4-1BB+ Treg subset enriched for the IL2-STAT5 suppressive pathway with transcription profiles supporting discrete metabolic alterations. Spatial analysis showed increased density of suppressive immune cells around tumor cells, increased exhaustion phenotype of both CD4 and CD8 T cells expressing chemokine CXCL13, and spatial micro-complex of endothelial and lymphocyte interactions within tertiary lymphoid structures. The single-cell architecture identifies determinants of early disease emergence and progression, which may be developed not only as diagnostic/prognostic biomarkers but also as targets for disease interception. Additionally, our dataset constitutes a valuable resource for the preinvasive lung cancer research community.

---

Despite advances in treatment strategies, the five-year survival of patients with advanced non-small cell lung cancer (NSCLC) remains at a dismal 20%*^1,2^*, as the majority of patients are diagnosed at advanced/metastatic stage where available treatments are less effective. Early disease detection significantly increases the likelihood of improved survival. Indeed, widespread utilization of CT scans in clinical care, as well as for screening for lung cancer, has led to increased detection of stage I lung cancer as well as the detection of subsolid pulmonary nodules commonly associated with preinvasive or minimally invasive malignancy*^3^*. Radiographically, these subsolid nodules are characterized as pure ground-glass opacity nodules (GGO), which are either predominantly non-solid in appearance and therefore do not obscure the underlying lung parenchyma, or as part-solid nodules that are comprised of a non-solid component with an adjacent solid component*^4^*. Although the natural history of subsolid nodules is not well-defined, approximately 30–40% of subsolid nodules progress to solid nodules containing more invasive adenocarcinoma within 4 years of detection. Therefore, radiographic detection of a subsolid nodule identifies patients at risk for subsequent invasive lung adenocarcinoma*^5,6^*. A major challenge in the clinical management of patients with these subsolid nodules is the lack of reliable biomarkers that can predict progression of preinvasive to invasive disease. Furthermore, effective therapies for intercepting the progression of preinvasive to invasive adenocarcinoma are lacking as the mechanisms that contribute to the progression are poorly characterized. We hypothesized that characterizing alterations in the tumor microenvironment (TME) has the potential to uncover determinants of early disease emergence, which can be developed as therapeutic targets for interception. Recent studies have begun to evaluate the transcriptome*^7^*, genome/epigenome*^8–13^*, immune contexture*^14–16^*, and extracellular matrix*^17^* in cohorts of patients with subsolid nodules compared to those with solid nodules of invasive adenocarcinoma. However, the small number of patients in most studies, coupled with marked interpatient heterogeneity and the logistical challenges that preclude the collection of longitudinal serial biopsies from the same nodules, present significant barriers to identifying clinically actionable pathways that contribute to malignant progression. We posited that these challenges may be addressed by studying part-solid nodules, where the solid component harbors more invasive adenocarcinoma while the non-solid component harbors minimally invasive or predominantly lepidic adenocarcinoma where cancer cells line the alveolar walls without stromal invasion. Direct comparison of matched samples from the non-solid and solid components from within the same nodule in individual patients may address issues related to interpatient heterogeneity. Importantly, such an analysis has the potential to identify not only the earliest changes associated with the initiation of non-solid nodules, through the comparison of the non-solid component to adjacent normal lung tissue, but also identify changes associated with the progression of non-solid nodules to invasive solid adenocarcinoma, through the comparison of the non-solid component to its matched solid counterpart. Therefore, we used samples obtained from patients with radiographically detected and histologically characterized part-solid nodules to construct the first high-resolution map at single-cell resolution, lineage/functional states, developmental trajectories and spatial composition as well as multicellular crosstalk networks as a function of disease progression. The workflow is shown in **Fig. 1a**. Importantly, analysis of an independent cohort of patients with pure non-solid and solid nodules largely corroborated the alterations identified in the non-solid and solid components of part-solid nodules.

**Figure 1:**
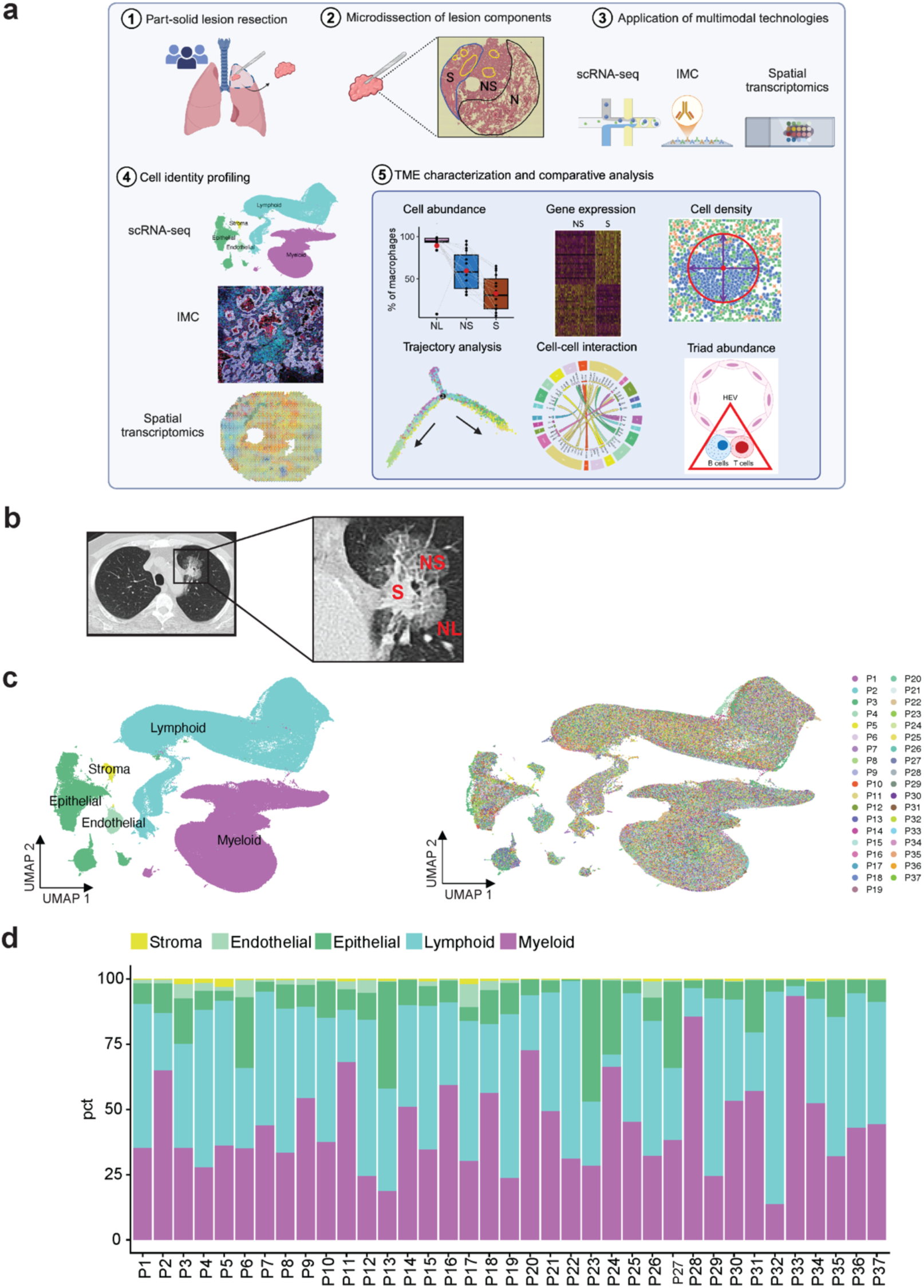
Multiomics analysis of microdissected components of part-solid nodules at single-cell resolution. a. Schematic of the workflow showing patients, technologies and data analysis used in this study. The schematic was created using BioRender (https://biorender.com). b. CT scan of a part-solid nodule showing normal lung (NL), non-solid (NS) and solid (S) components. c. UMAP plot showing 457,923 cells generated from 37 patients (89 samples) listed in Supplementary Table 1 and colored by their major cell type annotation (left panel) or by patient IDs (right panel). d. Percentage of major cell types in individual patients. pct, percentage

## Results

### Single-cell RNA-seq profiling of patients with part-solid nodules

The clinicopathological features of 89 samples collected prospectively from 37 patients in this study are shown (**Supplementary Table S1**). The cohort comprised 67.6% (n=25) females and 32.4% (n=12) males, with a median age of 73 years (range 43 to 91). The majority of patients were former smokers (n=22, 59.5%), three were current smokers (8.1%) and 12 were never smokers (32.4%). Twenty-four patients were Caucasian (64.9%), eight were Asians (21.6%), and one was African American (2.7%). One resected nodule was sampled from each patient, except for one patient, where three resected nodules were sampled (a total of 39 nodules). From freshly resected part-solid nodules (**Fig. 1b**, **Supplementary Fig. 1.1a-b**), discrete non-solid and solid components, along with matched adjacent uninvolved normal lung tissue, were microdissected by a pulmonary pathologist and subjected to single-cell RNA sequencing (scRNA-seq). A total of high-quality 457,923 cells were obtained after stringent quality control described in Methods. Integrative analysis yielded 24 high-confidence cell clusters representing 5 immune and non-immune major cell lineages (**Fig. 1c**). In total, 53,149 epithelial, 200,854 lymphoid, 193,379 myeloid, 7,906 endothelial and 2,635 stroma single-cell transcriptomes were obtained from all patients. The distribution of these cell types is depicted in individual patients (**Fig. 1d**).

### Tumor cell heterogeneity, cell of origin and transcriptome changes associated with disease progression

Reclustering of epithelial cells identified 21 distinct clusters, with the major clusters comprising club, ciliated, alveolar type 1 (AT1), alveolar type 2 (AT2) and basal cells (**Fig. 2a, Supplementary Fig. 2.1a- c, Supplementary Table S2, S3)**. Clusters EP.1, EP.6-11, EP.14-16 and EP.20 consisted of malignant tumor cells characterized by elevated copy number variation (CNV) scores (**Fig. 2b, Supplementary Fig. 2.2a**). Among them, clusters EP.1, EP.7-10, EP.15-16, and EP.20 were annotated as ‘Tumor’ and EP.14 as ‘Tumor proliferating’. As expected, high-CNV epithelial cells were enriched in non-solid and solid components compared to the normal epithelial cells in adjacent normal lungs (**Supplementary Fig. 2.2a**). Additionally, cluster EP.14, ‘Tumor proliferating’ was increased in the solid component compared to the non-solid component **(Supplementary Fig. 2.2b**). The remaining two high-CNV clusters, EP.6 and EP.11, were discretely located between normal club cells and AT2 cells (**Fig. 2b, Supplementary Fig. 2.1a**) and expressed markers of transitional club-AT2 cells (*SCGB3A1*, *SCGB3A2*, *SFTPD*, *NAPSA*; **Supplementary Fig. 2.2c**) suggesting they are early malignant cells transitioning from either club, AT2, or transitional club-AT2 cells*^18^*. These ‘Tumor transition’ cells showed enrichment of ‘glycolysis’ and ‘MYC targets V1’ pathways, which contribute to cell proliferation, growth and metabolism*^19,20^* (**Fig. 2c**). They also showed increased expression of protumorigenic genes, including *SPINK1* (Serine Peptidase Inhibitor Kazal Type 1)*^21^*, *MDK* (Midkine)*^22^*, *AQP3* (Aquaporin 3)*^23^* and *CEACAM5* (Carcinoembryonic antigen-related cell adhesion molecule 5)*^24^* (**Supplementary Fig. 2.2d**). Consistent with the UMAP, trajectory analysis revealed inferred progression of normal AT2 and club cells via the tumor-transition cells to tumor cells (**Fig. 2d**). Based on these findings, we sought to evaluate the similarity of tumor cells to AT2 and club cells. AT2 and club cell signatures (**Supplementary Table S4)** were used to assign tumor cells as ‘AT2-like’ (higher AT2 score) or ‘club-like’ (higher club score). Notably, 27/37 patients had predominantly ‘AT2-like’ or ‘club-like’ cells, and a comparison of the AT2 and club scores of the tumor cells per patient showed that 19 patients had AT2-like tumors, and eight patients had club-like tumors (**Fig. 2e, Supplementary Fig. 2.3a**). Ten patients remained unassigned due to a low number of tumor cells or low expression of the selected markers. These findings suggest that tumor heterogeneity may be attributed, among other factors to tumor cells-of-origin as shown in mouse models*^25^*.

**Figure 2:**
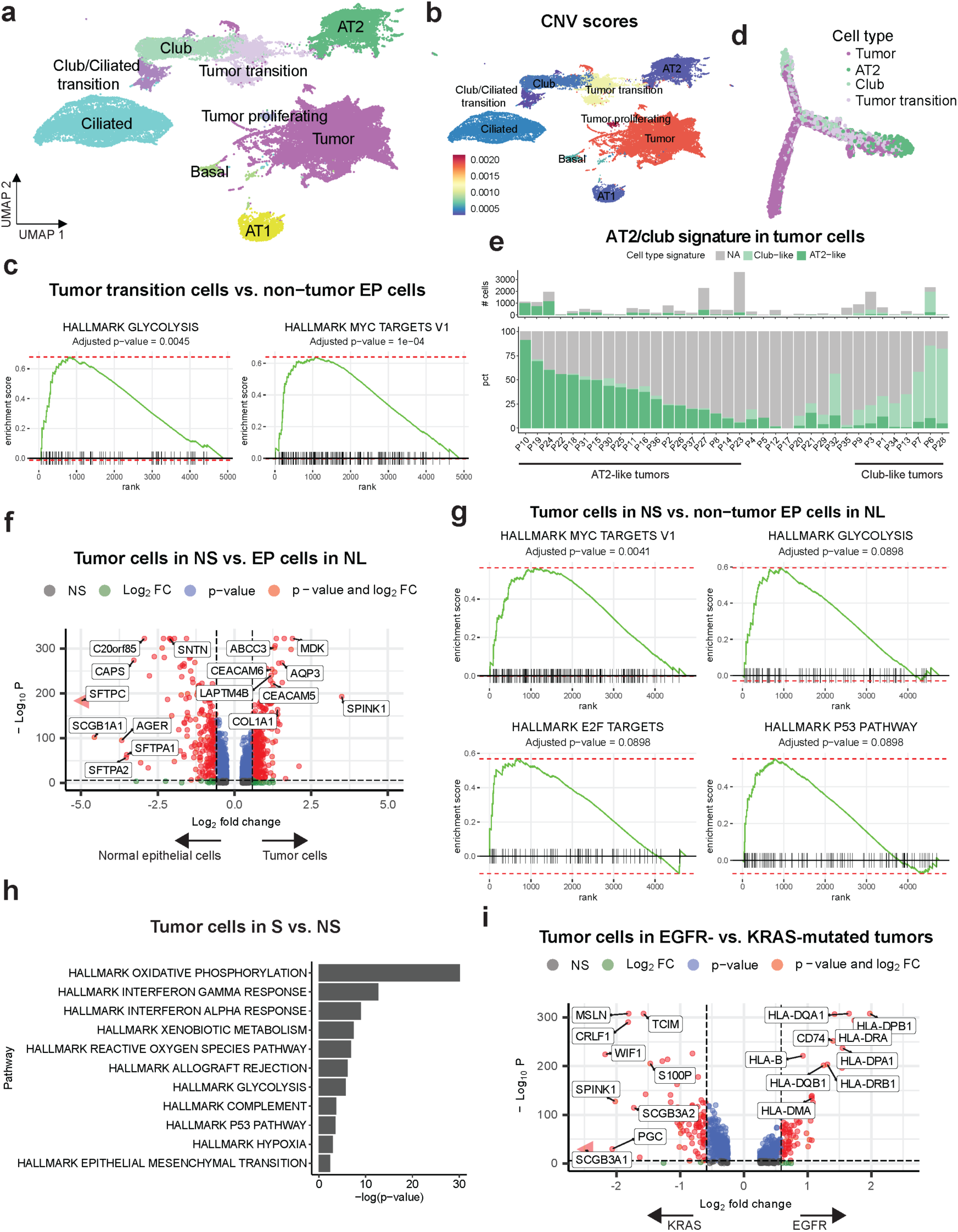
Epithelial cell heterogeneity in non-solid and solid components of the part-solid nodule. a. UMAP projection of 51,746 epithelial cells from all patients, categorized into 9 major cell types. b. UMAP plot showing mean CNV scores per cell type. c. Enrichment plots of upregulated Hallmark pathways in tumor transition cells compared to non-tumor epithelial cells. Enrichment scores and p-values were obtained using GSEA, with an FDR-adjusted p-value reported. d. Trajectory plot generated from Monocle2 analysis, illustrating the inferred progression of AT2 and club cells towards tumor cells. Cells are color-coded by the cell type. e. The percentage (lower panel) and numbers (upper panel) of tumor cells with AT-like signature, club-like signature, or without any identified signature (NA) across patients. Patients whose tumors were identified as AT2-like tumors were grouped on the left, while patients whose tumors were identified as club-like tumors were grouped on the right. pct, percentage. f. Volcano plot comparing gene expression between tumor cells in non-solid (NS) and normal epithelial cells from adjacent normal lungs (NL). The x-axis represents the log2 fold change in gene expression, with positive values indicating higher expression in tumor cells and negative values indicating higher expression in non-tumor epithelial cells. The y-axis represents the -log10 of the p-value, with higher values indicating greater statistical significance. Green dots indicate genes with significant fold changes (≥ 1.5), blue dots indicate genes with significant adjusted p-values (≤ 0.05), and red dots indicate genes significant in both fold change and p-value. g. Enrichment plots of upregulated Hallmark pathways in tumor cells in the non-solid (NS) component compared to non-tumor epithelial cells in adjacent normal lung (NL). Enrichment scores and p-values were obtained using GSEA, with an FDR-adjusted p-value reported. h. Upregulated Hallmark pathways in tumor cells from the solid component compared to tumor cells from the non-solid component. P-values were obtained using GSEA. i. Volcano plot comparing gene expression between tumor cells in EGFR vs KRAS mutated tumors. The x-axis represents the log2 fold change in gene expression, with positive values indicating higher expression in EGFR-mutated tumors and negative values indicating higher expression in KRAS-mutated tumors. The y-axis represents the -log10 of the p-value, with higher values indicating greater statistical significance. Green dots indicate genes with significant fold changes (≥ 1.5), blue dots indicate genes with significant adjusted p-values (≤ 0.05), and red dots indicate genes significant in both fold change and p-value.

Tumor cells were enriched in the solid component compared to the non-solid, with a concomitant decrease in the proportion of AT1 and AT2 cells (**Supplementary Fig 2.3b**). To identify cancer-cell intrinsic pathways that govern the emergence of early disease, we compared tumor cells in the non-solid component with epithelial cells in adjacent normal lung. In addition to protumorigenic genes observed in the tumor transiting population above, there was upregulation of (i) *COL1A1* (Collagen type I alpha 1) which promotes epithelial-to-mesenchymal transition (EMT), and confers poor prognosis*^26,27^*; (ii) *ABCC3* (ATP Binding Cassette Subfamily C Member 3) known to confer multidrug resistance in lung cancer*^28^*; (iii) *CEACAM6^24^*; and (iv) *LAPTM4B* (Lysosomal Protein Transmembrane 4 Beta), an oncogene known to stimulate tumor growth*^29^* (**Fig. 2f**). Additionally, there was marked enrichment of the ‘MYC targets V1’, ‘glycolysis’ and ‘E2F targets’ pathways, which contribute to cell proliferation, growth, and metabolic alterations*^19,20,30^*, as well as enrichment of the ‘P53 pathway’, suggesting increased response to oncogenic stress and DNA damage*^31^* (**Fig. 2g**). Compared to the non-solid component, tumor cells in the solid component showed upregulation of metastasis-promoting genes (*SERPINA1^32^*, *CXCL14^33^* and *FTL^34^*; **Supplementary Fig. 2.4a, Supplementary Table S5**), and pathways including ‘oxidative phosphorylation’, ‘glycolysis’, ‘interferon alpha response’, ‘interferon gamma response’, and the ‘reactive oxygen species (ROS) pathway’ (**Fig. 2h)**, suggesting that metabolic alterations and oxidative stress may be associated with the progression of non-solid nodules to solid adenocarcinoma. Importantly, tumor cells from patients with EGFR mutation showed increased expression of several MHC class I and II genes compared to tumor cells from patients with KRAS mutant tumors, along with an increase in CD74, which regulates the presentation of MHC class II proteins (**Fig. 2i)**. These findings are notable in light of previous studies showing that EGFR mutations are negatively correlated with the expression of MHC molecules*^35^*. EGFR mutant tumor cells also showed upregulation of ‘interferon gamma response’ and ‘allograft rejection’, whereas KRAS-mutated tumor cells showed upregulation of the ‘TNFa signaling via NFkB pathway’, consistent with previous studies*^36^* (**Supplementary Fig. 2.4b**). To consider the possibility that the upregulation of HLA genes in EGFR tumors may be specific to preinvasive disease, we interrogated advanced-stage LUAD patient samples (TCGA) and observed a similar increase in HLA and CD74 in EGFR mutant patients compared to KRAS patients (**Supplementary Fig. 2.4c**). Together, these findings underscore the significant complexity and heterogeneity in tumor cell-of-origin, impact of driver oncogenes and alteration in gene expression, not only as the earliest changes in non-solid nodule formation but also with the progression of non-solid (preinvasive) to solid (invasive) adenocarcinoma.

### Lymphocytes show decreased cytotoxic and increased suppressive phenotypes

To dissect the identity and functional phenotypes of the lymphocytic population, we performed unsupervised clustering, which identified major cell types including CD8 and CD4 T, NK, B and plasma cell clusters (**Fig. 3a, Supplementary Fig. 3.1a, Supplementary Table S2)**. Analysis of relative abundances showed an increase in CD4 T cells, and a decrease in NK cells in the non-solid component compared to normal lungs and in the solid compared to non-solid (**Fig. 3b**). Reclustering of CD8 T cells identified naïve, cytotoxic, memory, inflammatory, and exhaustion phenotypes (**Fig. 3c, Supplementary Fig. 3.1b, 3.2a and 3.3a, Supplementary Table S6**). Clusters CD8.3 and CD8.5 with increased expression of cytotoxic markers (*GNLY, PRF1, FGFBP2, FCGR3A/CD16,* and *NKG7*; **Supplementary Fig. 3.1b and 3.2a**), and a high cytotoxicity score (**Fig. 3d, Supplementary Fig. 3.3a**, **Supplementary Table S4**), showed progressively decreased abundance in the non-solid and solid components (**Fig. 3e**). Cluster CD8.7 resembling NKT-like cells expressing T cell receptor delta (TCRδ), and with increased cytotoxicity score was also decreased in abundance in the non-solid and solid components (**Supplementary Fig. 3.3b).** Cluster CD8.11 (*CCL3*^high^) with mem/effector phenotypes and expressing cytotoxic markers did not show changes in abundance. Conversely, CD8.8 (CXCL13^high^) expressing exhaustion markers (*CTLA4, ENTPD1/CD39, HAVCR2/TIM3, LAG3, TOX* and *PDCD1*; **Supplementary Fig. 3.1b, 3. 2a**) and exhibiting the highest exhaustion score (**Fig. 3f, Supplementary Fig. 3.3a)** was increased in the non-solid and further in the solid component (**Fig. 3g**). These results suggest that reduced cytotoxic and increased exhaustion phenotypes of CD8 T cells may be critical determinants of not only non-solid nodule formation, but also their progression to solid adenocarcinoma. To determine developmental trajectories of the CD8 T cell clusters, we applied Monocle 2*^37^* and set the root point to cluster CD8.2 that expressed naive cell markers including *CCR7, TCF7, LEF1,* and *SELL*, and also GZMK, indicating that this cluster may contain both naive cells and transition cells*^38^* (**Supplementary Fig. 3.1b, 3.2a and 3.3a**). The trajectory showed two main primary branches, one leading to cytotoxic phenotype (state 1; **Fig. 3h, Supplementary Fig. 3.4a-b**) and the other to exhaustion phenotype (state 5; **Fig. 3h**, **Supplementary Fig. 3.4a-c**). Notably, CD8.1 expressing pre-exhausted and exhausted markers (GZMK, RGS1), was enriched on the naïve/transition branch (states 2 and 4), and together with CD8.4 (*ZNF683*, *KLRC1*/*NKG2A*) was enriched along the exhausted branch (state 5; **Fig. 3h, Supplementary Fig. 3.4d**), suggesting that CD8.1 and CD8.4 may represent pre-exhausted CD8 T cells*^39^*. In the non-solid and solid components, these pre-exhausted clusters were increased in abundance and showed elevated exhaustion scores (**Supplementary Fig. 3.4e-f**). The trajectory structure did not significantly alter between normal, non-solid, and solid components (**Supplementary Fig. 3.4c)**. It is noteworthy that both the CD8.8 (exhausted) and CD8.4 (pre-exhausted) clusters expressed tissue-resident memory (TRM) markers including *ITGAE*/*CD103, ITGA1/CD49a, CD69* and *CXCR6* (**Supplementary Fig. 3.1b**), and furthermore, the exhausted branch exhibited higher TRM scores (state 5; **Supplementary Fig. 3.4b**). These findings are consistent with previous studies and suggest that TRM cells undergo exhaustion during tumor progression*^40,41^*. Together, these data suggest that disease progression is associated with an increased abundance of pre-exhausted CD8 T cells, which progressively acquire enhanced dysfunctional phenotypes.

**Figure 3:**
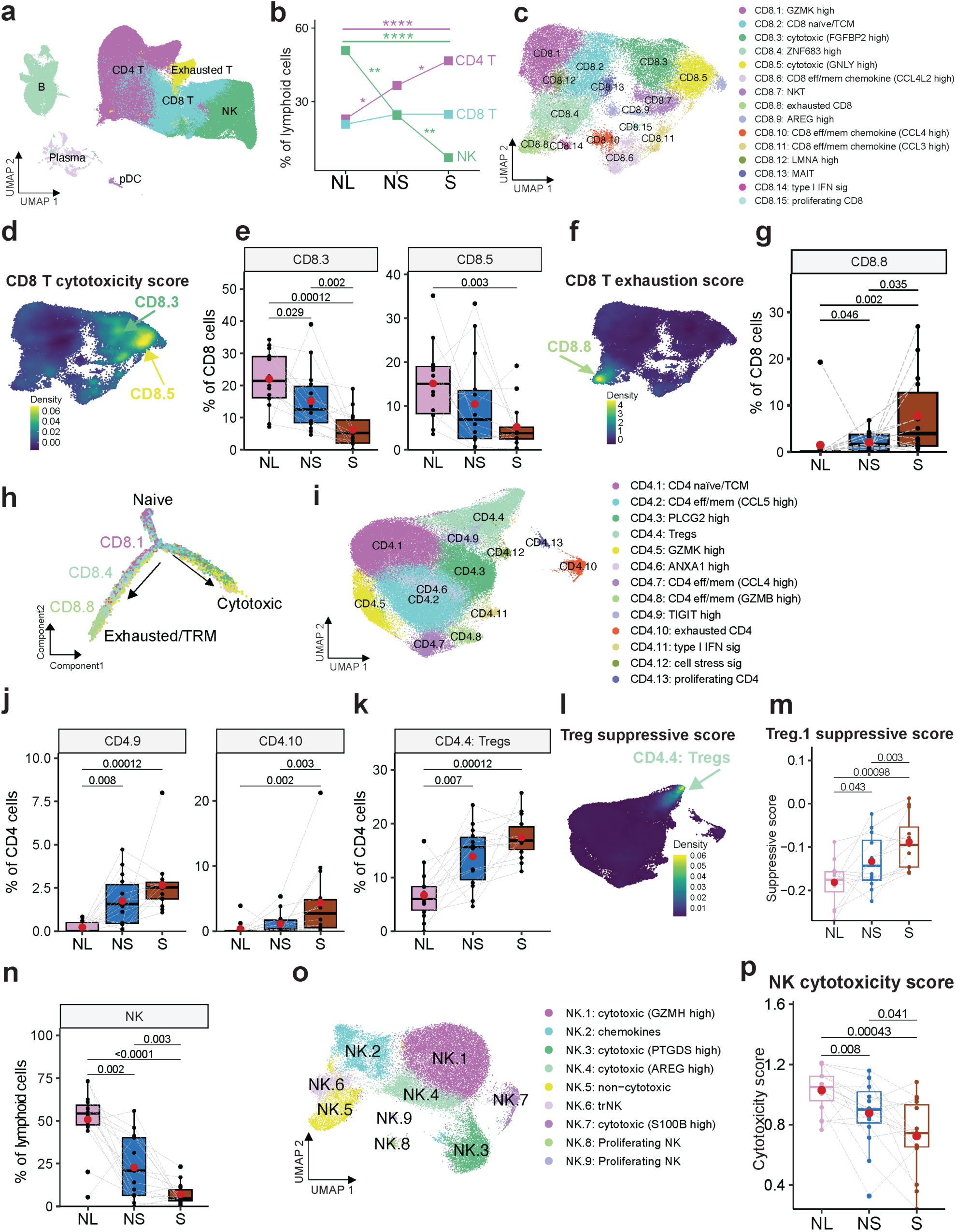
Lymphoid cell subtypes, abundance and functional states in non-solid and solid components of part-solid nodules. (a) UMAP projection of 197,188 lymphoid cells from all patients categorized into 7 major cell types. (b) Major lymphoid cell type abundance in adjacent normal lungs (NL), non-solid (NS) and solid (S) components. Percentages are based on the entire lymphoid cell population. P-values were obtained using the Wilcoxon matched-pairs signed rank test on paired samples from the same patients. *: p < 0.05, **: p < 0.01, ***: p < 0.001, ****: p < 0.0001. Only significant p-values (< 0.05) are shown. (c) UMAP visualization of reclustered CD8 T cells, showing distinct CD8 T cell subsets. (d) Density plot showing cytotoxicity scores of CD8 T cells. (e) Proportion of cytotoxic CD8 T cell populations (CD8.3 and CD8.5) in adjacent normal tissue (NL), non-solid (NS) and solid (S) lesion components. Percentages are based on the entire CD8 T cell population. P-values were obtained using the Wilcoxon matched-pairs signed rank test on paired samples from the same patients, with matched samples connected by a line. Mean values are indicated by red dots. Only significant p-values (< 0.05) are shown. (f) Density plots showing exhaustion score of CD8 T cells. (g) Proportion of exhausted CD8 T cell population (CD8.8) in adjacent normal tissue (NL), non-solid (NS) and solid (S) lesion components. Percentages are based on the entire CD8 T cell population. P-values were obtained using the Wilcoxon matched-pairs signed rank test on paired samples from the same patients, with matched samples connected by a line. Mean values are indicated by red dots. Only significant p-values (< 0.05) are shown. (h) Trajectory plot depicting the inferred progression from naïve CD8 T cells to cytotoxic or exhausted/TRM states using Monocle2 analysis. Cells are color-coded as in panel (c). (i) UMAP visualization of reclustered CD4 T cells showing distinct CD4 T cell subsets. (j) Proportion of TIGIT^high^ (CD4.9) and exhausted (CD4.10) CD4 T cell populations in adjacent normal tissue (NL), non-solid (NS) and solid (S) lesion components. Percentages are based on the entire CD4 T cell population. P-values were obtained using the Wilcoxon matched-pairs signed rank test on paired samples from the same patients, with matched samples connected by a line. Mean values are indicated by red dots. Only significant p-values (< 0.05) are shown. (k) Proportion of Treg (CD4.4) cell population in adjacent normal tissue (NL), non-solid (NS) and solid (S) lesion components. Percentages are based on the entire CD4 T cell population. P-values were obtained using the Wilcoxon matched-pairs signed rank test on paired samples from the same patients, with matched samples connected by a line. Mean values are indicated by red dots. Only significant p-values (< 0.05) are shown. (l) Density plot displaying the suppressive scores of Treg cells. (m) Suppressive scores of Treg.1 cells in adjacent normal tissue (NL), non-solid (NS) and solid (S) components. Scores are averaged per sample. P-values were obtained using the Wilcoxon matched-pairs signed rank test on paired samples from the same patients, with matched samples connected by a line. Mean values are indicated by red dots. Only significant p-values (< 0.05) are shown. (n) Proportions of NK cells in adjacent normal tissue (NL), non-solid (NS) and solid (S) lesion components. Percentages are based on the entire lymphoid cell population. P-values were obtained using the Wilcoxon matched-pairs signed rank test on paired samples from the same patients, with matched samples connected by a line. Mean values are indicated by red dots. Only significant p-values (< 0.05) are shown. (o) UMAP visualization of reclustered NK cells showing distinct NK cell subsets. (p) Chemokine-expression scores of cytotoxic NK cells (i.e., excluding NK.5) in adjacent normal tissue (NL), non-solid (NS) and solid (S) components. Scores are averaged per sample. P-values were obtained using the Wilcoxon matched-pairs signed rank test on paired samples from the same patients, with matched samples connected by a line. Mean values are indicated by red dots. Only significant p-values (< 0.05) are shown.

Given the increased abundance of CD4 T cells in both non-solid and solid components (**Fig. 3b**), we determined their subtypes and activation/suppressive phenotypes. Reclustering of CD4 T cells yielded major subtypes including naïve/TCM (CD4.1), exhausted (CD4.10), memory/effector (CD4.2, CD4.7, CD4.8) and Tregs (CD4.4) (**Fig. 3i, Supplementary Fig. 3.1c, 3.5 and 3.6a-b, Supplementary Table S7).** CD4.10 showed increased expression of exhaustion markers (*TIGIT*, *CTLA4, TOX, LAG3, PDCD1*), elevated exhaustion score (**Supplementary Fig 3.1c, 3.5 and 3.6a)**, and also expressed *CXCL13* characteristic of human T_FH_ cell*^42^* **(Supplementary Fig 3.1c and 3.5)**. CD4.9 expressed both inhibitory markers (*TIGIT, TOX*, *PDCD1*) and naïve/TCM markers (*LEF1, CCR7, TCF7, SELL*; **Supplementary Fig. 3.1c, 3.5, and 3.6a**), representing a transitioning population between naïve and exhausted CD4 T cells*^43^*. Notably, clusters CD4.9 and CD 4.10 showed increased abundance and elevated exhaustion scores in the non-solid and the solid components **(Fig. 3j, Supplementary Fig. 3.6b**). The rest of the CD4 T cells (excluding CD4.4, CD4.9 and CD4.10) also showed increased exhaustion scores in the solid component compared to non-solid (**Supplementary Fig. 3.6c).**

Clusters CD4.6 (ANXA1, LMNA, CRIP1, VIM) described as effector/memory CD4 T cells in the gut*^44^*, and CD4.7 (*CCL4*^high^) with increased expression of effector/memory markers and chemokines (*GZMA, GZMH, IFNG, PRF1, CCL3, CCL4* and *CCL5* showed decreased abundance in solid compared to non-solid component (**Supplementary Fig. 3.6d, Supplementary Fig. 3.1c and 3.5a**). CD4.6 expressed *MYADM* and *ANKRD28* known to support cell migration and tissue homing*^45^* and LMNA which augments Th1 polarization to attenuate Treg differentiation*^46^* (**Supplementary Fig. 3.5)**. Together, these data suggest that an immunosuppressive microenvironment, characterized by increased CD8 and CD4 T cell exhaustion and decreased cytotoxic CD8 T cells, may contribute not only to non-solid nodule formation but also to the progression of non-solid to solid adenocarcinoma.

### 4-1 BB+ Tregs with marked suppressive activity are increased in the early nodules

Cluster CD4.4 expressing characteristic markers of regulatory T cells (Tregs: *FOXP3, IKZF2*/*HELIOS, IL2RA*/*CD25, CTLA4*; **Supplementary Fig. 3.1c, 3.5)** showed increased abundance in the non-solid component compared to normal lungs, and further increased in solid compared to non-solid (**Fig. 3k-l**). Reclustering of Tregs (CD4.4) identified three discrete subclusters (**Supplementary Fig 3.7a**). Treg.1 showed elevated expression of naïve markers (*LEF1, SELL, CCR7, and TCF7*), Treg.2 showed elevated expression of suppressive markers (*FOXP3, CTLA4, TIGIT, PDCD1*/*PD1, HAVCR2*/*TIM3, ICOS*, *LAG3*), and Treg.3 displayed an intermediate phenotype (**Supplementary Fig. 3.7b**). While we did not observe a significant increase either in the relative abundance or suppressive scores of Treg.2 and Treg.3 as a function of disease progression, Treg.1 suppressive scores increased in non-solid compared to normal lungs and further increased in solid compared to non-solid **(Fig. 3m).** Further characterization of the Treg subclusters showed that Treg.2 had the highest suppressive score (**Supplementary Fig. 3.7c)**, and showed upregulation of the ‘IL-2-STAT5 signaling’ pathway which is known to induce *FOXP*3 and mediate Treg differentiation and elicit suppressive phenotypes*^47,48^* (**Supplementary Fig. 3.7d**).

Compared to Treg.1, Treg.2 showed significantly up-regulated genes (*TNFRSF4*/*OX40*, *TNFRSF18*/*GITR*, *TNFRSF9/4-1BB, TNFRSF1B*/*TNFR2* and *LAIR2*) associated with suppressive functions of tumor-infiltrating Tregs*^49^* (**Supplementary Fig. 3.7e, Supplementary Table S8**). Notably, gene signature from *TNFRSF9/4-1BB*^high^ Tregs was predictive of worse overall survival in TCGA LUAD*^50^*. Additionally, the Treg.2 cluster showed altered expression of key metabolic regulators (*PKM*, *ENO1*, *GAPDH*, *ENP1,* and *LDHA*)*^51^*, (**Supplementary Fig. 3.7e**), and enhancement of pathways including ‘oxidative phosphorylation’, ‘glycolysis/gluconeogenesis’, ‘metabolism of polyamines’, ‘metabolism of lipids’, ‘fatty acid metabolism’, and ‘pyruvate metabolism’, associated with Treg suppressive phenotype*^52–55^* (**Supplementary Fig. 3.7f**). Particularly, ‘oxidative phosphorylation’ and ‘glycolysis/gluconeogenesis’ were increased in the solid components (**Supplementary Fig. 3.7g**). Trajectory analysis showed Tregs progression from naïve/resting-like towards a suppressive phenotype, with enrichment of naïve/resting Treg.1 on the upper-right branch (state 1), enrichment of Treg.3 on the lower-right branch (state 2), and divergence of highly suppressive Treg.2 from the main branch (state 7) **(Supplementary Fig. 3.7h-j).** In states 1 and 2, Treg.1 and Treg.3 showed limited *FOXP3* expression and conspicuous *IL2RA*/*CD25* and *IKZF2*/*HELIOS* expression (**Supplementary Fig. 3.7k**), suggesting that these cells are in a transitional phase, shifting from CD4 naïve T cells to Tregs in state 1 and CD4+IKZF2+Foxp3− T cells in state 2*^56^*. The suppressive score was increased along the trajectory peaking at state 7, indicating a diverse population of Treg cells within the TME, each possessing varying degrees of suppressive capabilities (**Supplementary Fig. 3.7l**). Together, these data indicate that suppressive Tregs contribute to the initiation and progression of non-solid nodules to adenocarcinoma.

### Decreased NK cell abundance and cytotoxicity are associated with chemokine expression

NK cells constitute an important arm of the innate immune system that contributes to protective antitumor immunity*^57^*. NK cell abundance was significantly decreased in the non-solid component relative to the normal lungs and in the solid compared to non-solid (**Fig. 3n**), consistent with a previous report*^16^*. NK cells have been characterized at single-cell resolution across many cancer types*^58^*, however, little is known about the heterogeneity and functional states of NK cells in human preinvasive nodules. Reclustering the NK cells identified 9 clusters with distinct gene expression programs indicative of significant functional specialization, including cytotoxicity and chemokine expression programs (**Fig. 3o, Supplementary Fig. 3.8b, Supplementary Table S9**). In particular, all subclusters excluding NK.5 and NK.6 showed increased expression of cytotoxic genes including *GZMA*, *GZMB*, *GZMH*, *GZMM*, *PRF1*, *GNLY* and *NKG7* (**Supplementary Fig. 3.8, Supplementary Fig. 3.9a)** and with high cytotoxic score (**Supplementary Fig 3.9b**), indicative of cytotoxic CD56^dim^ NK phenotype. Clusters with high cytotoxicity scores (NK.1, NK.3 and NK.4) showed reduced abundance, whereas clusters with low cytotoxicity scores (CD56^bright^ NK cells: NK.5 and NK.6), showed increased abundance in the non-solid and solid components (**Supplementary Fig 3.9c**). Importantly, the overall NK cytotoxicity score was markedly decreased in the non-solid component relative to the normal lung, and in the solid compared to the non-solid (**Fig. 3p)**. NK.6 showed increased expression of *GZMK,* which was shown to be upregulated in NK cells in melanoma*^59^*, as well as *ITGA1*/*CD49a*, *CD69*, and *CXCR6*, which serve as markers for tissue-resident NK (trNK) cells*^60–62^*. Notably, *CD69*^high^ trNK cells have been shown to accumulate in lung cancer*^60–62^*. NK.5 and NK.6 also showed increased expression of the chemokines *XCL1* and *XCL2* (**Supplementary Fig 3.8b**), which are known to recruit XCR1^high^ cross-presenting cDC1 cells into tumors*^59^*. Cluster NK.2 showed increased expression of chemokines *CCL4, CCL4L2, CCL3,* and *CCL3L1* (**Supplementary Fig. 3.8, Supplementary Fig. 3.9**), and was associated with high chemokine-expressing score (**Supplementary Fig. 3.9b**). Possibly, NK chemokines recruit key immune cell populations required for generating protective tumor immunity*^63^*.

Among the remaining lymphoid cell types, we observed enrichment of B cells in the solid component compared to adjacent normal lungs (**Supplementary Fig 3.10a**). Within the subclusters of B and plasma cells (memory B, naïve B, plasma IgG/IgA/IgM/IgD and plasmablast; **Supplementary Fig 3.10b-c**), germinal center B cells were enriched in the solid component compared to the normal lungs (**Supplementary Fig 3.10d**). The overall increase in B cells, particularly the germinal center B cells, may be attributed to the presence of tertiary lymphoid structures (TLS).

### Myeloid cells exhibit marked heterogeneity with enhanced immunosuppressive phenotypes

Unsupervised clustering of the myeloid population identified monocytes, macrophages, dendritic cells (DC), polymorphonuclear myeloid-derived suppressor cells (PMN-MDSCs) and neutrophils (**Fig. 4a, Supplementary Fig. 4.1a)**. Monocytes, neutrophils and macrophages were decreased, whereas DCs and PMN-MDSCs were increased in non-solid and solid components compared to normal lungs (**Fig. 4b**). To dissect the heterogeneity of the macrophage population*^64^*, we reclustered and identified 11 subtypes (**Fig. 4c, Supplementary Fig. 4.1b, Supplementary Table S10)**. Use of gene signatures of monocyte-derived macrophages (MoMac) and tissue-resident alveolar macrophages (TRM-AM) from early-stage NSCLC*^65^* identified clusters Mac.2, Mac.5, and Mac.6 as MoMacs and the remaining clusters as TRM-AMs (**Supplementary Fig. 4.1a-b**). Next, we assessed the pro- vs. anti-inflammatory activity of macrophages*^66^*. Of the MoMac clusters, Mac.5 showed increased expression of proinflammatory genes (*CXCL9, CXCL10* and *CXCL11*), associated with increased pro-inflammatory (M1) score (**Supplementary Fig. 4.1, Supplementary Fig. 4.2a,c**), and increased expression of proinflammatory transcription factors including *STAT1, IRF1*, *IRF2, IRF7* and *IRF9^67,68^* (**Supplementary Fig. 4.2d**). Mac.5 abundance was significantly increased in the solid component compared to normal lung (**Supplementary Fig. 4.2e)**. In contrast to Mac.5, Mac.2 and Mac.6 were associated with anti-inflammatory (M2) score (**Supplementary Fig. 4.2a, Supplementary Fig. 4.2c**). We observed expression of *SELENOP* in Mac.2 and *SPP1* in Mac.6 (**Supplementary Fig. 4.1b and 4.2a**), which have been shown to contribute to macrophage polarization from pro-inflammatory (M1-like) to anti-inflammatory (M2-like)*^69,70,71^* and also confer poor prognosis in LUAD*^72,73^* (**Supplementary Fig. 4.2a)**. The Mac.2 and Mac.6 clusters also showed increased expression of *TREM2* (triggering receptor expressed on myeloid cells 2; **Supplementary Fig. 4.2f)**. In lung cancer, TREM2^high^ macrophages constrain NK and T cell function*^74^*, and targeting these macrophages is being considered as a therapeutic modality. Importantly, the M2-like Mac.2 and Mac.6 clusters were significantly increased in the non-solid component relative to the normal lungs and in the solid component compared to the non-solid (**Fig. 4d**). In contrast to the MoMac clusters, TRM-AMs which were abundant in normal lungs, decreased significantly in the non-solid and solid components (**Fig. 4e).** Notably, the M2 score of both TRM-AM and all the macrophages increased in the non-solid component relative to the normal lungs, and in the solid compared to non-solid (**Fig. 4f**). Moreover, TRM-AMs in the non-solid and solid component showed decreased expression of TRM-AM markers (*FABP4*, *PDLIM1* and *IGFBP2)* and increased expression of MoMac markers (*BASP1*, *CEBPD*, *TMEM176B*, *TREM2*, *APOE*, *A2M*, *SPP1*, *MARCKS*), suggesting that the TRM-AMs may have been reprogramed into tumor-associated macrophage in the non-solid and solid nodules (**Supplementary Fig. 4.2g**). These results suggest that marked enhancement of anti-inflammatory/ immunosuppressive phenotypes of M2-like macrophages contributes to the initiation of non-solid nodules and their progression to solid adenocarcinoma.

**Figure 4.**
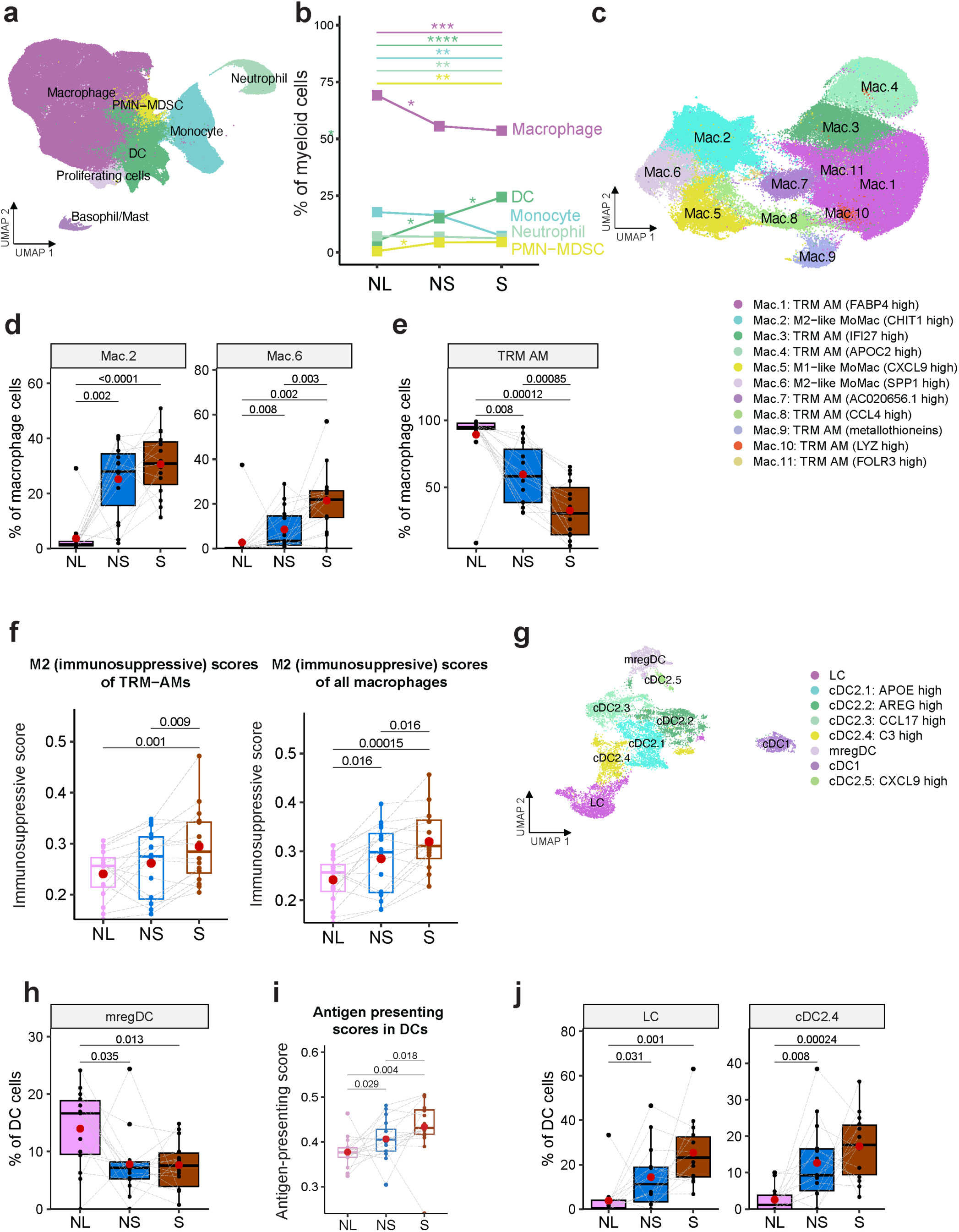
Myeloid cell subtypes, abundance and functional states in non-solid and solid components of part-solid nodules. a. UMAP projection of 193,379 myeloid cells from all patients, categorized into 7 major cell types. b. Comparison of the major myeloid cell types in adjacent normal tissue (NL), non-solid (NS) and solid (S) lesion components. Percentages are based on the entire myeloid cell population. P-values were obtained using the Wilcoxon matched-pairs signed rank test on paired samples from the same patients. *: p < 0.05, **: p < 0.01, ***: p < 0.001, ****: p < 0.0001. Only significant p-values (< 0.05) are shown. c. UMAP visualization of reclustered macrophages, illustrating 11 distinct clusters. d. Proportions of M2-like MoMac cell populations (Mac.2 and Mac.6) in adjacent normal tissue (NL) and solid (S) lesion component. Percentages are based on the entire macrophage cell population. P-values were obtained using the Wilcoxon matched-pairs signed rank test on paired samples from the same patients, with matched samples connected by a line. Mean values are indicated by red dots. Only significant p-values (< 0.05) are shown. e. Proportions of TRM-AM cell populations (Mac.1,3-4,7-11). in adjacent normal tissue (NL) and solid (S) lesion component. Percentages are based on the entire macrophage cell population. P-values were obtained using the Wilcoxon matched-pairs signed rank test on paired samples from the same patients, with matched samples connected by a line. Mean values are indicated by red dots. Only significant p-values (< 0.05) are shown. f. M2 (immunosuppressive) scores of TRM-AM (left) and all macrophages (right) in adjacent normal tissue (NL), non-solid (NS) and solid (S) components. Scores are averaged per sample. P-values were obtained using the Wilcoxon matched-pairs signed rank test on paired samples from the same patients, with matched samples connected by a line. Mean values are indicated by red dots. Only significant p-values (< 0.05) are shown. g. UMAP visualization of reclustered DCs. h. Proportions of mregDC in adjacent normal tissue (NL) and solid (S) lesion component. Percentages are based on the entire DCs population. P-values were obtained using the Wilcoxon matched-pairs signed rank test on paired samples from the same patients, with matched samples connected by a line. Mean values are indicated by red dots. Only significant p-values (< 0.05) are shown. i. Antigen-presenting scores of DCs in adjacent normal tissue (NL), non-solid (NS) and solid (S) components. Scores are averaged per sample. P-values were obtained using the Wilcoxon matched-pairs signed rank test on paired samples from the same patients, with matched samples connected by a line. Mean values are indicated by red dots. Only significant p-values (< 0.05) are shown. j. Proportions of top antigen-presenting DC populations (LC and cDC2.4) in adjacent normal tissue (NL) and solid (S) lesion component. Percentages are based on the entire DC population. P-values were obtained using the Wilcoxon matched-pairs signed rank test on paired samples from the same patients, with matched samples connected by a line. Mean values are indicated by red dots. Only significant p-values (< 0.05) are shown.

Dendritic cells play key roles in initiating and maintaining anti-tumor T cell immunity*^75^*. There was an increased abundance of DCs in the non-solid component relative to the normal lungs, and in the solid compared to the non-solid component (**Fig. 4b**). To dissect DC heterogeneity, we reclustered DCs and identified discrete subsets including type 1 conventional DC (cDC1) expressing *XCR1* and *CLEC9A*, type 2 conventional DC (cDC2) expressing *CLEC10A* and *AREG*, mature-regulatory DCs (mregDC) and Langerhans cells (LCs) (**Fig. 4g, Supplementary Fig. 4.3a-b, Supplementary Table S11**). cDC2s exhibited marked heterogeneity with several subpopulations: cDC2.1 (*APOE^high^*), cDC2.2 (*AREG^high^*), cDC2.3 (*CCL17^high^*) cDC2.4 (*C3^high^*) and cDC2.5 (CXCL9*^high^*) (**Supplementary Fig. 4.4a)**. Of these, cDC2.1 exhibited the highest inflammatory score (*CD14, S100A8, S100A9, and VCAN*) (**Supplementary Fig. 4.4a and 4.5a**). cDC2.4 showed marked expression of complement pathway genes including classical *C1Q* (*C1QA*, *C1QB*, and *C1QC*), and alternative *C3* **(Supplementary Fig. 4.3a, Supplementary Table S11)**, characteristic of monocyte-derived DCs*^76^*. Notably, the complement pathway was upregulated in cDC2 in non-solid compared to normal lungs and in solid compared to non-solid with upregulation of the complement genes including *C1QA*, *C1QB*, and *C1QC* (**Supplementary Fig. 4.4b-c**). C1Q enhances the chemotaxis of mature DC to CXCL19 via upregulation of CCR7 expression*^77^*. However, the role of the DC-specific complement pathway in preinvasive lung cancer is not well understood.

mregDCs expressing immunoregulatory genes *CD274 (PD-L1*), *PDCD1LG2* (*PD-L2*), *IDO1*, *IL4I*, *CD200*, and *EBI3* (*IL27B*) were associated with the highest immunosuppressive score*^78^* (**Supplementary Fig. 4.3b and 4.5a-b**). Moreover, mregDCs expressed CCR7 which facilitates their migration to lymph nodes for antigen presentation*^79^*, and chemokines CCL19 and CCL22 which facilitate the recruitment of CCR4+ Tregs*^80,81^* (**Supplementary Fig. 4.3a and 5b**). mregDC abundance was decreased in the non-solid and solid components compared to the normal lungs (**Fig. 4h**). Given that DCs are efficient antigen-presenting cells, we assessed DCs for their antigen-presentation capabilities. Clusters LC, cDC2.1, cDC2.4 and cDC2.5 conferred the highest antigen-presenting score (**Supplementary Fig. 4.5c**). Notably, the overall antigen-presenting score was higher in the non-solid component relative to the normal lungs, and in the solid compared to the non-solid (**Fig. 4i**), consistent with increased abundance of LC and cDC2.4 in the non-solid relative to the normal lungs, and in the solid compared to the non-solid (**Fig. 4j**). Additionally, gene-set enrichment analysis (GSEA) showed that cDC2s in the non-solid component were associated with an enrichment of the GO gene-set ‘antigen processing and presentation’ (**Supplementary Fig. 4.5d**). These findings suggest that DCs increase antigen presentation, perhaps due to a greater prevalence of tumor neoantigens as a function of disease progression*^82^*.

### Endothelial cell heterogeneity contributes to angiogenesis modulation and immune suppression

To dissect endothelial cell heterogeneity, we used signatures from the lung endothelial cell (EC) atlas*^83^* and categorized EC clusters into general capillary, aerocyte capillary, systemic venous, pulmonary venous, artery and lymphatic ECs (**Fig. 5a, Supplementary Fig. 5.1a-b, Supplementary Table S12**). The relative abundance of aerocyte capillary ECs and lymphatic ECs was decreased in the solid and non-solid components, respectively, whereas systemic venous ECs abundance, a major fraction of angiogenic ECs in lung tumors*^84^*, was increased (**Fig. 5b)**. ECs in the non-solid and solid component showed increased expression of proangiogenic genes (*PLVAP*, *HSPG2, FLT1, SPRY1*, *ANGPT2*, *COL15A1*, and *COL18A1)*, and downregulation of *HPGD* (15-Hydroxyprostaglandin Dehydrogenase), which is expressed by aerocyte capillary ECs and degrades prostaglandins in vessels to overcome immunosuppression*^85^* (**Fig. 5c, Supplementary Fig. 5.1a,c, Supplementary Table S13**). Increased angiogenesis corresponded with decreased expression of immunomodulatory genes (*IL7R*, *ICAM1*, *BIRC3*, *IRF1*; **Fig. 5c**) and downregulation of immune pathways in the solid component (**Fig. 5d**). Consistent with the immune phenotype, we observed decreased expression of *IRF1* and *JUN* regulons (**Fig. 5e)**, previously implicated with diminished immune activity in lung tumor ECs*^86^*. Conversely, there was upregulation of *MAFB* and *FOXC1* regulons, which regulate endothelial sprouting and angiogenesis*^87,88^*, and of MYC regulon, reported to be upregulated in ECs in NSCLC*^86^* (**Fig. 5e**). Taken together, ECs show increased angiogenic phenotypes with a concomitant reduction in immune activity suggesting the potential of EC-directed therapeutic strategies to intercept disease progression.

**Figure 5.**
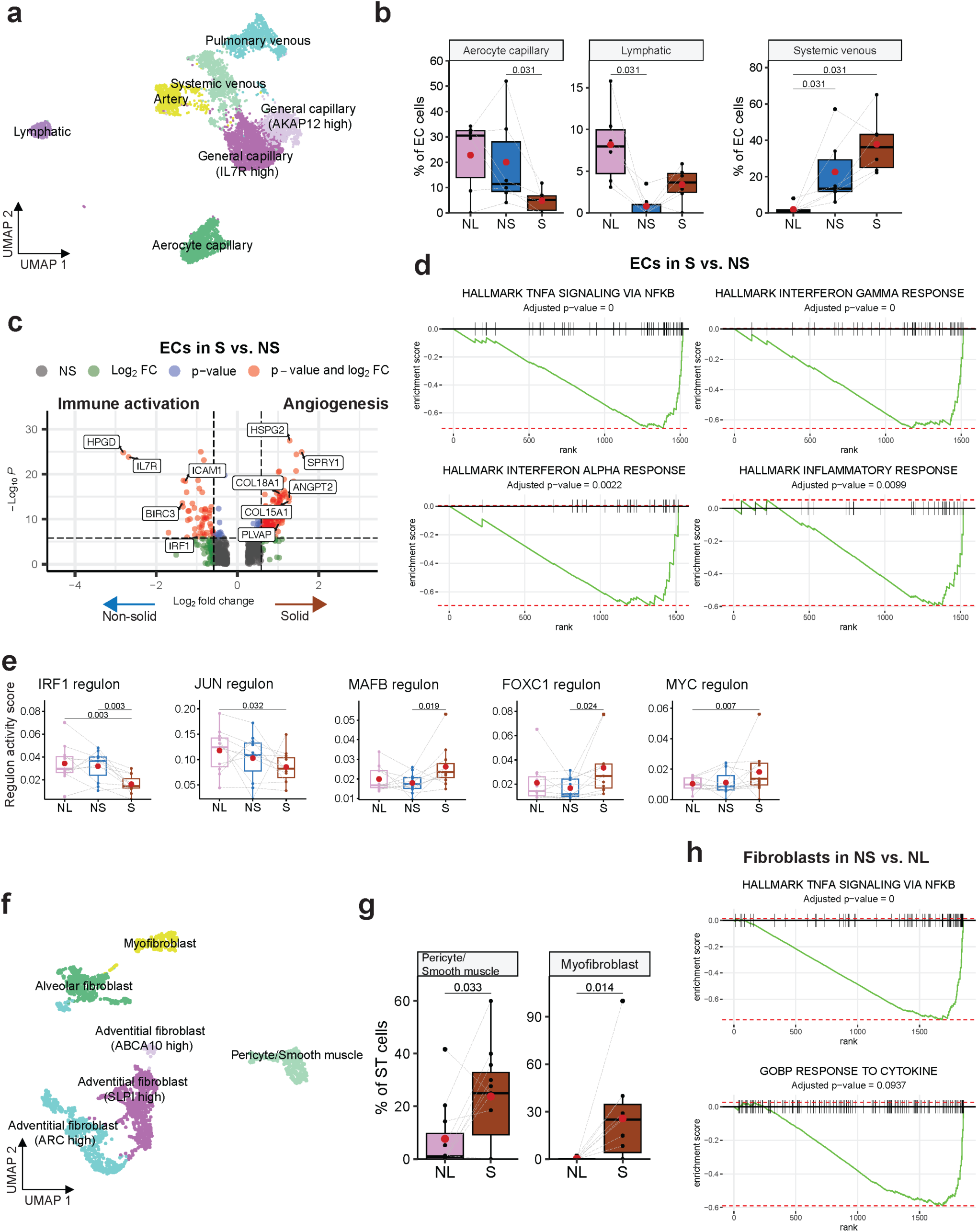
Stromal cell subtypes, abundance and functional states in non-solid and solid components of part-solid nodules. (a) UMAP projection of 7,565 endothelial cells (EC) from all patients, categorized into six major cell types. (b) Proportions of aerocyte capillary, lymphatic, and systemic venous EC populations in adjacent normal tissue (NL) and solid (S) lesion component. Percentages are based on the entire EC population. P-values were obtained using the Wilcoxon matched-pairs signed rank test on paired samples from the same patients, with matched samples connected by a line. Mean values are indicated by red dots. Only significant p-values (< 0.05) are shown. (c) Volcano plot comparing gene expression in ECs in solid (S) vs. non-solid (NS). The x-axis represents the log2 fold change in gene expression, with positive values indicating higher expression in solid samples and negative values indicating higher expression in non-solid samples. The y-axis represents the -log10 of the p-value, with higher values indicating greater statistical significance. Green dots indicate genes with significant fold changes (≥ 1.5), blue dots indicate genes with significant adjusted p-values (≤ 0.05), and red dots indicate genes significant in both fold change and p-value. (d) Enrichment plot of Hallmark pathways down-regulated in ECs in the solid (S) compared to the non-solid (NS) component. Enrichment scores and p-values were obtained using GSEA, with an FDR-adjusted p-value shown. (e) Regulon activity of the IRF1, JUN, MAFB, FOXC1 and MYC regulons in ECs from normal (NL), non-solid (NS) and solid (S) components. Scores are averaged per sample. P-values were obtained using the Wilcoxon matched-pairs signed rank test on paired samples from the same patients, with matched samples connected by a line. Mean values are indicated by red dots. Only significant p-values (< 0.05) are shown. (f) UMAP projection of 2,435 fibroblasts from all patients, categorized into four major cell types. (g) Proportions of pericyte/smooth-muscle cell and myofibroblast populations in adjacent normal tissue (NL) and solid (S) lesion component. Percentages are based on the entire fibroblast population. P-values were obtained using the Wilcoxon matched-pairs signed rank test on paired samples from the same patients, with matched samples connected by a line. Mean values are indicated by red dots. Only significant p-values (< 0.05) are shown. (h) Enrichment plots of down-regulated pathways in fibroblasts in the non-solid (NS) compared to the normal tissue (NL). Enrichment scores and p-values were obtained using GSEA, with an FDR-adjusted p-value shown.

### Fibroblasts and extracellular matrix (ECM) remodeling contribute to tumor progression and immunosuppression

Reclustering of stroma cells identified alveolar fibroblasts, adventitial fibroblasts, myofibroblasts, and pericytes/smooth muscle cells (**Fig. 5f, Supplementary Fig. 5.2a-b; Supplementary Table S2, S14**), with an increase in myofibroblast and pericyte/smooth muscle cells in the solid component (**Fig. 5g**). Pericytes/smooth muscle cells, important regulators of angiogenesis and vascular integrity expressed *ACTA2*/α-SMA, *COL4A1*, and *MYH11* (**Supplementary Fig. 5.2c, Supplementary Table S14**), markers previously implicated in T-cell exclusion by cancer-associated fibroblasts (CAFs) in early-stage lung adenocarcinoma*^89^*. Myofibroblasts, predominantly found in the nodule, expressed genes related to ECM/collagenase (*MMP11, COL10A1, COL1A1, COL3A1, COL11A1, COL8A1* and *COL5A2*), migration (*POSTN^90^, INHBA^91^*), regulation of T-cell localization (*FAP^89^*) and fibroblast-to-myofibroblast transition markers (IGFBP3*^92^*,*TGFB1^93^*; **Supplementary Fig. 5.2c**).

In the non-solid component compared to the normal tissue, fibroblasts showed increased expression of genes associated with EMT and ECM remodeling (*BGN*, *COL4A1*, *COL4A2*, *ACTN1* and *MYLK*; **Supplementary Fig. 5.2d**), potentially contributing to fibrosis*^94^*, and conversely, downregulation of *ADH1B* associated with T-cell permissive CAFs*^89^* (**Supplementary Fig. 5.2d**). Additionally, there was downregulation of immune pathways ‘TNFa signaling via NFkB’ and ‘response to cytokines’ (**Fig. 5h**), which may drive an immunosuppressive TME in the early phase*^95^*. In the solid component, the ‘extracellular matrix binding’ pathway was upregulated, indicating active ECM remodeling by fibroblasts*^96^* (**Supplementary Fig. 5.2f**). These alterations underscore the role of CAFs role in ECM remodeling, angiogenesis and T-cell exclusion facilitating tumor growth and immune evasion.

### Immune cell abundance and phenotypes in the non-solid and solid components of part-solid nodules corroborated in an independent cohort of patients with pure non-solid and solid nodules

To determine if the non-solid and solid components of a part-solid nodule accurately represent the TME of pure non-solid or solid nodules, we used an independent in-house cohort of bulk RNA-seq samples from both non-solid (n=98) and solid (n=37) nodules, along with matched adjacent normal lung (n=29). For a direct comparison, scRNA-seq cell type abundances were recalculated as a proportion of total cells (**Supplementary Fig. 5.3a**). Deconvolution of bulk RNA-seq and enrichment analysis using gene signatures from the scRNA-seq analysis (**Supplementary Table S15)** revealed similar patterns of changes with T, B, plasma, total and activated dendritic cells increasing; and NK cells, endothelial cells and neutrophils decreasing in the nodules vs. normal lungs (**Supplementary Fig. 5.3b**). CD4 T cells and Tregs exhibited a trend towards increase consistent with that observed with scRNA-seq (**Supplementary Fig. 5.3a-b**). Additionally, there was a significant enrichment of the suppressive Treg signature in both non-solid and solid nodules (**Supplementary Fig. 5.3c**). Moreover, there was an increase in T-cell exhaustion and pre-exhaustion signature enrichment scores and a decrease in T-cell cytotoxicity scores, in agreement with the scRNA-seq results. While we observed an increase in general macrophage abundance, M2 macrophage abundance did not increase. Therefore, we applied the M2-like MoMac cluster signatures from our scRNA-seq data (Mac.2: CHIT1^high^ SELENOP ^high^, Mac.6: SPP1^high^) and found that Mac.6 signature was enriched in solid nodules (**Supplementary Fig. 5.3c)**. Notably, there was a decrease in TRM-AM scores in the nodules, associated with an increase in MoMac signatures, indicating a shift from resident to recruited macrophages consistent with scRNA-seq findings (**Supplementary Fig. 5.3c**). These data suggest that the TME of the non-solid and solid components of the part-solid nodule recapitulates, to a large extent, the TME of pure non-solid and solid nodules.

### Cell-cell communication identifies complex dynamics of angiogenesis modulation and immunosuppression

Tumor progression is supported by dynamic crosstalk between tumor cells and immune/stromal cells within the TME*^97^*. Using the MultiNicheNet platform*^98^*, which computes ligand-receptor interactions and the differential expression of target genes downstream of the receptor, we identified differential cell-cell communications (CCC) in both the non-solid and solid components (**Supplementary Fig. 6.1a-b, Supplementary Table S16**). In the non-solid component compared to normal lungs, the top 20 CCCs (**Fig. 6**) included epithelium as the sender cell, suggesting that the TME alterations are driven majorly by tumor cells. *VEGFA* ligand expressed on epithelial cells (cancer cells and AT1 cells) communicated with *FLT1* (VEGF receptor 1), *KDR* (VEGF receptor 2), and *NRP1* (a coreceptor for *KDR*) expressed on ECs*^99^* (**Fig. 6a-b**). Notably, the VEGF-VEGFR-activated pathway network in the endothelium included the venous EC-specific proangiogenic gene *PLVAP*, suggesting that the angiogenesis-promoting transcriptomic changes observed in the ECs may be initiated by the tumor cells (**Fig. 6c**). Tumor cells also expressed *MDK* (Midkine), a secreted growth factor with an affinity for receptor ITGA4 on monocytes (**Fig. 6a,d**), an interaction that facilitates the recruitment of monocytes to tumor sites*^100^*, consistent with increased abundance of MoMacs in the TME. Additionally, *CXCL16,* expressed by tumor cells and PMN-MDSC, interacted with *CXCR6* expressed by exhausted and pre-exhausted CD8 T cells (**Fig. 6a, d-e**), possibly facilitating their recruitment to generate an immunosuppressive phenotype*^101^*. The non-solid component also showed M2-like MoMac interactions including *APOE/TREM2* and *APOE/SCARB1*, which affect lipid metabolism and contribute to anti-inflammatory phenotype*^102,103^*, and *CCL18/CCR1* interactions which leads to M2 polarization of MoMacs*^104^* (**Fig. 6a, f**). Notably, *CCL18* has been reported to inhibit CCR1-mediated chemotaxis*^105^*.

**Figure 6.**
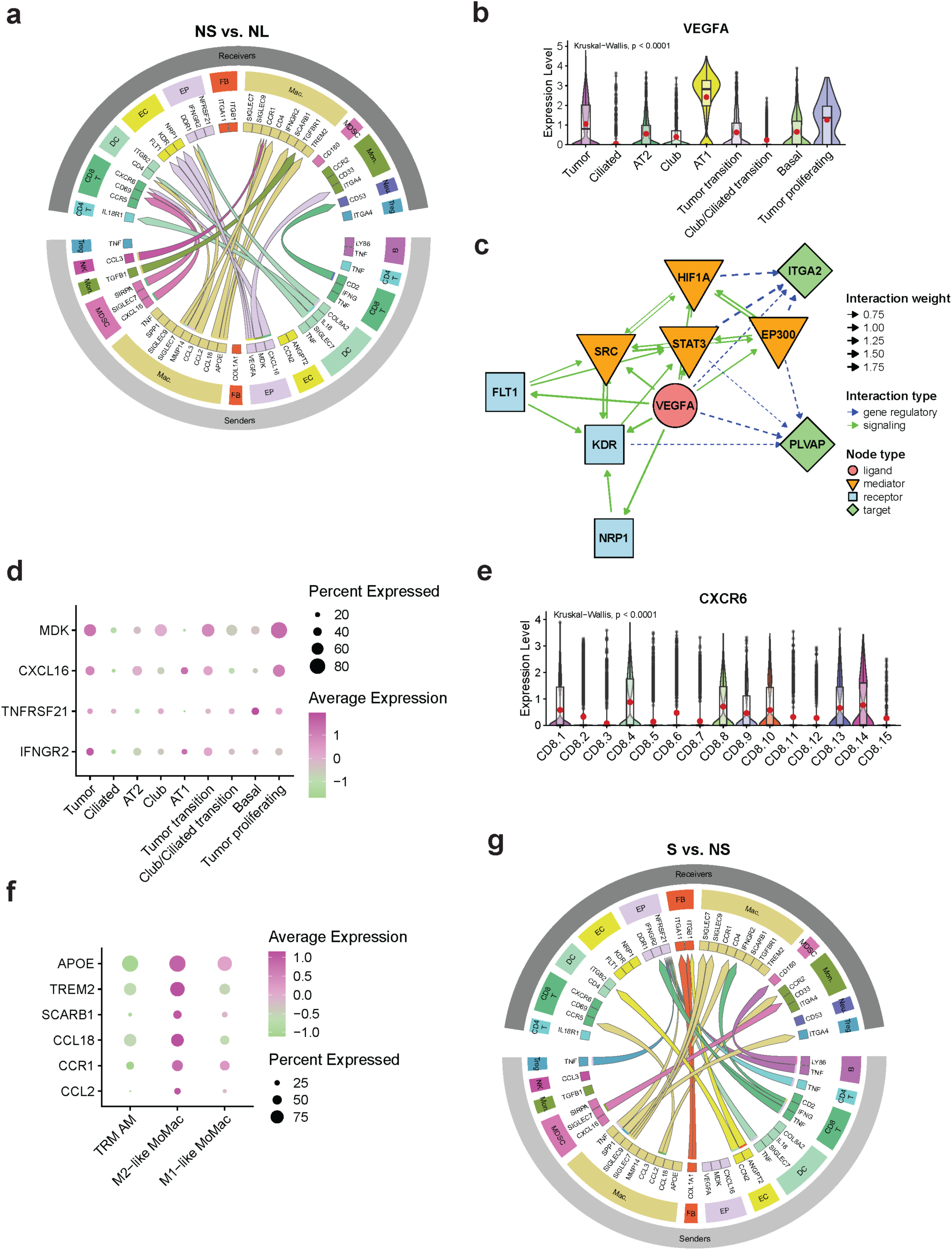
Differential cell-cell communications in non-solid and solid components of part-solid nodules. a. A circos plot displays the top 20 differential ligand-receptor interactions between cells in the TME in non-solid (NS) vs. normal (NL) by MultiNicheNet analysis. Sender and receiver cells are shown. FB, Fibroblasts; EP, Epithelial cells; B, B cells; Exh T, Exhausted T cells, Mac., macrophages; Mon., monocytes, Neu., neutrophils. b. *VEGFA* expression across epithelial cell types. Mean values are indicated by red dots. P-value was calculated using the Kruskal-Wallis test. c. Signaling and regulatory pathways between the *VEGFA* ligand expressed by epithelial cells, and the *FLT1*, *NRP1*, and *KDR* receptors, as well as downstream target genes, expressed in endothelial cells. d. Dot plot displaying the expression of selected ligands and receptors in epithelial cells. Expression level is depicted by color, and the size of the dots represents the proportion of cells expressing the genes. e. *CXCR6* expression across CD8 T cell clusters. Mean values are indicated by red dots. P-value was calculated using the Kruskal-Wallis test. f. Dot plot displaying the expression of selected ligands and receptors in macrophages. Expression level is depicted by color, and the size of the dots represents the proportion of cells expressing the genes. g. A circos plot displays the top 20 differential ligand-receptor interactions between cells in the TME in solid (S) vs. non-solid (NS) by MultiNicheNet analysis. Sender and receiver cells are shown. FB, Fibroblasts; EP, Epithelial cells; B, B cells; Exh T, Exhausted T cells, Mac., macrophages; Mon., monocytes, Neu., neutrophils.

In the solid compared to the non-solid, *TNF* expressed in multiple immune cells (Treg, B, DC, CD8 T, CD4 T and macrophages) interacted with *TNFRSF21* in epithelial cells which may enhance tumor cell survival and immune evasion (**Fig. 6g**)*^106,107^*. Interactions were also observed between M2-like MoMacs expressing CCL2 and the receptors CCR5 and CCR2, expressed on CD8 T cells and monocytes, respectively (**Fig. 6g**). These interactions indicate the recruitment of myeloid and T cells to the solid nodule*^108^*, specifically M2-like macrophages*^104,109^*.

Interestingly, within the solid component there was an interaction between CD8 T cells expressing *IFNG* (mem/eff CD8) and macrophages and epithelial cells expressing *IFNGR2* (**Fig. 6g**). IFN-γ is a cytokine with dual roles in cancer capable of both promoting and inhibiting tumor growth, by virtue of inhibiting proliferation, inducing apoptosis and promoting immune evasion by upregulating PD-L1 on tumor cells*^110^*. These analyses highlight the dynamic cell-cell communications within the TME that drive tumor initiation, progression, and immune regulation.

### Spatial proteomics and transcriptomics analysis show an increased density of suppressive immune cells and a decreased density of cytotoxic T cells proximal to tumor cells

To further elucidate the TME and investigate its spatial landscape during tumor progression, we applied spatial proteomics using multiplexed imaging mass cytometry (IMC) and spatial transcriptomics (ST, 10X Visium) to 8 patients with part-solid nodules from our cohort (**Supplementary Fig. 7.1a**). For IMC, a customized 40-plex antibody panel was used to identify tumor, immune and stromal cells corresponding to our scRNA-seq analysis (**Supplementary Table S17**). Using H&E images as a guide, 71 regions of interest (ROIs) were selected to capture the non-solid (n=30) and solid (n=23) components of the part solid nodules and adjacent normal lung tissue (n=18; **Fig. 7a)**, leading to the identification of 24 cell phenotypes **(Supplementary Fig. 7.1b, 7.2**). Compared to the normal lung, the non-solid and solid components showed an increased density of T cells (CD4 and CD8), myeloid cells (CD33+HLA-DR+), B-cells (CD20+), fibroblasts (αSMA+Pan-CK-), endothelial cells (CD31+PanCK-) and a decrease in NK (CD56+NCR1+) cells (**Supplementary Fig. 7.3a**). More specifically, in the myeloid population there was a decrease in TRM-AMs (CD68+CD11c+) and an increase in M1-like macrophages (CD14+CD68+CD163-), M2-like macrophages (CD14+CD68+CD206+CD163+) and MDSCs (PMN-MDSC: CD11b+CD15+, monocytic MDSC: CD11b+CD14+; **Supplementary Fig. 7.3b**), consistent with the scRNA-seq analysis. The spatial landscape was also characterized using spatial transcriptomics (ST, 10X Visium), and cell type annotations and expression signatures from the scRNA-seq data were used to deconvolute Visium spots (**Supplementary Fig. 7.4a**). A comparison of cell type relative abundances revealed an increased abundance of CD8 T cells, CD4 T cells, Tregs, B/plasma cells and DCs along with a decreased abundance of monocytes and neutrophils in both the non-solid and solid components, consistent with findings from scRNA-seq and IMC analyses (**Supplementary Fig. 7.4b**). We next determined the proximity of immune cells to tumor cells (**Fig. 7b**). Compared to non-solid, the solid component exhibited an increased density of suppressive immune cells, including exhausted CD4 and CD8 T cells (TOX+PD-1+), and Tregs (FoxP3+). In contrast, there was a decreased density of activated CD4 T cells (CD45RA+GITR+TOX-) and CD8 T cells (CD45RA+GzmB+TOX-) in proximity to tumor cells (**Fig. 7b**). Naïve CD4 and CD8 T cells did not show proximity changes relative to tumor cells. Of the myeloid cells, the solid components showed increased density of M2-like and SPP1+ M2-like macrophages and a decreased density of M1-like macrophages and DCs (CD68-CD11c+; **Fig. 7b**). These data suggest that increased density of suppressive immune cells and decreased density of cytotoxic T cells around the tumor cells contributes to tumor initiation and subsequent progression by conferring resistance to immune-mediated clearance.

**Figure 7.**
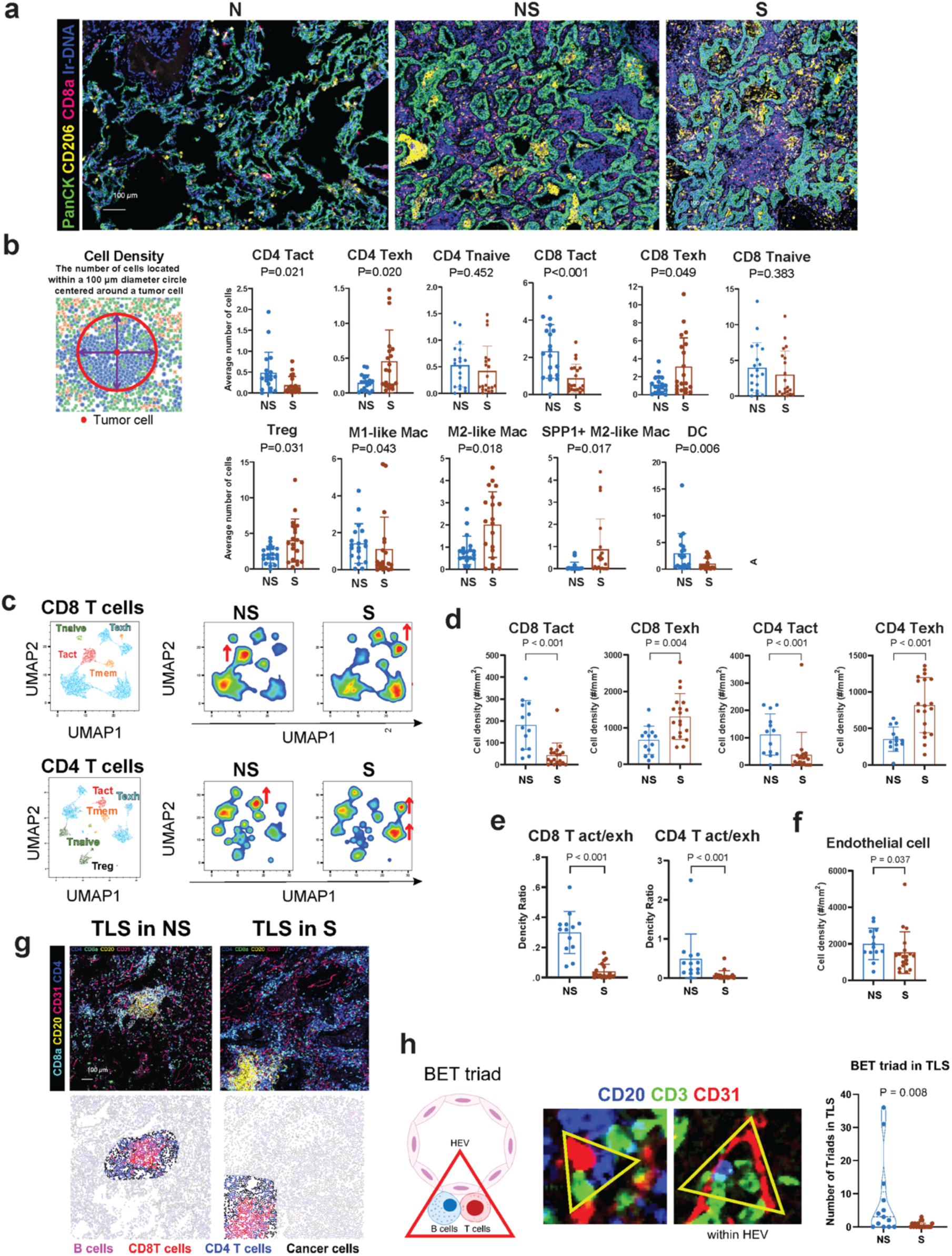
IMC analysis showing an increased density of suppressive immune cells around tumor cells and TLS with B cells surrounded by exhausted T cells. a. Representative multichannel IMC images from normal lung (N), non-solid (NS) and solid (S) regions. PanCK, tumor cells; CD206, macrophages; CD8a, CD8+ T cells. b. Cell density analysis showing immune cells within 100 µm radius centered on a tumor cell. act, activated; exh, exhausted. c. UMAP of CD8 T cells (upper panel) and CD4 T cells (lower panel) showing T naive, T activated (Tact), T memory (Tmem), exhausted T cells (Texh) and Tregs in TLS of NS and S components. Red arrows indicate an increase. d. Cell densities (per mm*^2^*) of specific CD4 and CD8 T cells in the TLS of NS and S components. e. Density ratios of CD8 T act vs CD4 Texh and CD4 T activated vs CD4 Texh. f. Representative multichannel IMC image (upper panel) and identified cells (lower panel) of TLS in NS and S components. CD4 T, CD8 T, endothelial cells and macrophage cells are color-coded. g. BET (B cell, endothelial cell, T cell) triads. Schematic and representative image (left panel) and enumeration in NS and S components (right panel).

### Spatial analysis shows T cell suppressive phenotypes in the TLS

Histopathological analysis of part-solid nodules showed pronounced TLSs both in the non-solid and solid components (**Supplementary Fig. 7.1a)**. Recent studies suggest that the impact of TLSs on survival and response to immunotherapy depends on both their cellular components and maturation stage*^111^*. To gain insights into TLS, we expanded the IMC analysis on TLS (21 ROIs with 32 TLSs) of non-solid and solid components (**Supplementary Fig. 7.5a**). While the TLS size did not differ significantly between the non-solid and solid components (**Supplementary Fig. 7.5b**), their cellular landscape did. Compared to non-solid, TLSs in the solid component showed a decreased density of activated CD8 and CD4 T cells and an increased density of exhausted CD8 and CD4 T cells, associated with a decrease in both CD8 and CD4 activation/exhaustion ratios, indicative of immunosuppressive phenotypes (**Fig. 7c-e, Supplementary Fig. 7.5c)**. These alterations were specific to TLS and not to regions outside the TLS (**Supplementary Fig. 7.5d**). Consistent with the IMC findings, ST analysis showed elevated exhausted T cell scores in the solid TLSs versus non-solid TLSs and in TLSs compared to non-TLS regions (**Supplementary Fig. 7.6a)**. TLS also showed increased expression of *CXCL13* in the solid TLSs versus non-solid TLSs and in TLSs compared to non-TLS regions (**Supplementary Fig. 7.6b**). Notably, *CXCL13*, expressed exclusively by exhausted T cells in the scRNA-seq data is a known driver of TLS formation and organization*^112^*.

Additionally, ST analysis showed an increased abundance of B/plasma cells in TLSs compared to non-TLS regions, suggesting that the observed increase in these cell populations in the scRNA-seq analysis was linked to TLS formation (**Supplementary Fig. 7.6c**). Further characterization with IMC showed abundant B cells surrounded by T cells and CD31+ endothelial cells in the TLSs (**Fig. 7f**). Interestingly, TLSs in the non-solid component had an increased number of triads comprised of closely interacting B cells, T cells and endothelial cells, indicative of high endothelial venules (HEV) and referred to as the ‘BET triad’ (**Fig. 7g**). These newly identified BET triads may facilitate immune cell recruitment from systemic circulation to support TLS function in the context of T cell activation and antigen presentation*^113,114^*. These data suggest that early disease emergence and subsequent progression are associated with increased exhaustion of both CD8 and CD4 T cells in TLS and that the accumulation of CXCL13+ exhausted T cells during tumor progression may promote the recruitment of B cells and facilitate the formation of TLSs.

## Discussion

Microdissection of matched non-solid and solid components from a part-solid nodule, along with adjacent normal lung allowed identification of not only the earliest changes associated with the initiation of the non-solid nodule but also changes associated with the progression of preinvasive to invasive adenocarcinoma (**Fig. 8**). Importantly, analysis of an independent cohort of patients with pure non-solid and solid nodules largely corroborated cell type abundance identified in the non-solid and solid components of part-solid nodules, supporting our unique approach. Early tumor initiation, as determined by the analysis of the non-solid component showed the presence of high-CNV epithelial cells enriched in protumorigenic and metabolic pathways. Trajectory analysis inferred progression of normal AT2 and club cells via the intermediary “tumor-transition cells” to malignant tumor cells. Indeed, assignment of AT2 and club scores of each tumor cell per patient categorized patients with either AT2-like tumors or club-like tumors, indicative of significant complexity and heterogeneity in tumor cells-of-origin. In this context, a recent study exclusively investigating epithelial cell states in early-stage human lung adenocarcinoma implicated “alveolar intermediary cells” as progenitors for mutant KRAS tumors*^115^*, consistent with AT2 cells implicated as the cell-of-origin in Kras-driven mouse lung cancer models*^116^*. Nonetheless, the cell-of-origin issue remains contentious, with evidence suggesting involvement of other cell types*^117^*, as well as the potential for club cells to trans-differentiate into AT2 cells*^118^*. Therefore, further investigations are warranted as identifying the cell(s) of origin for lung cancer is critical for designing better chemoprevention strategies.

**Figure 8.**
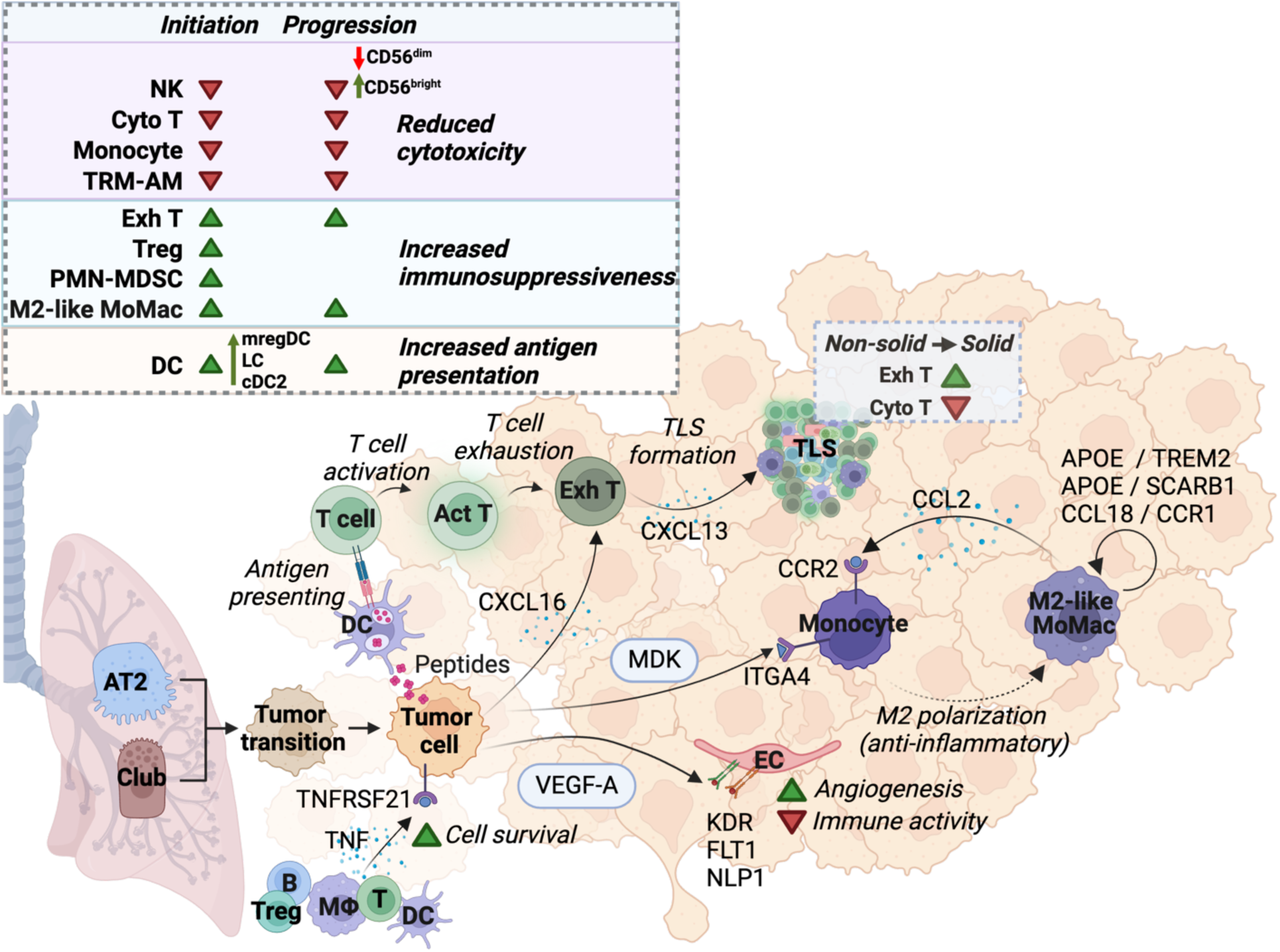
Immune and cellular dynamics in initiation of preinvasive nodules and progression to invasive lung adenocarcinoma. A summary of the observed changes in immune cell populations and their interactions within the TME during the progression of lung adenocarcinoma. The upper panel illustrates changes in key immune cell populations during tumor initiation (non-solid vs. normal) and progression (solid vs. non-solid). A reduction in cytotoxic T cells (Cyto T), NK cells and monocytes contributes to decreased cytotoxicity, while an increase in Tregs, exhausted T cells (Exh T), PMN-MDSCs and M2-like MoMac fosters immunosuppression. DCs exhibit increased antigen presentation, accompanied by a rise in total DCs, as well as mregDC, LC and cDC2 subpopulations. The lower panel highlights cell-cell interactions within the TME that drive immune modulation, angiogenesis, and immune suppression. These dynamics underscore the transition from non-solid to solid tumor states in lung adenocarcinoma.

In the solid component, tumor cells showed further enhancement of protumorigenic genes, metabolic alterations and oxidative stress, suggesting that these pathways may support the progression of non-solid nodules to solid adenocarcinoma. Notably, EGFR-mutant tumors showed markedly increased expression of HLA cluster genes (both MHC class I and II) along with CD74, which is known to regulate the presentation of MHC class II proteins. While these findings contradict reports showing that mutant EGFR suppresses MHC I expression and is negatively correlated with MHC expression in advanced LUAD*^119^*, it is conceivable that during tumor initiation, there is a fine balance between an active immune response and a concomitant tumor-elicited immune suppression. Supporting this concept, studies have shown that non-solid nodules have substantially lower rates of HLA deletions than advanced lung cancer*^10,120^*.

Tumor initiation, as determined by analysis of the non-solid component, revealed two major modes of immunosuppression spanning both the lymphoid and myeloid phenotypes. Particularly, the lymphoid compartment showed an increased abundance of suppressive Tregs and M2-like macrophages along with a reduction in cytotoxic CD8 T and NK cells and accumulation of exhausted CD4 and CD8 T cells, indicative of impaired protective antitumor immunity*^57,121^*. This immunosuppressive microenvironment appeared to constitute critical determinants not only of the earliest changes associated with non-solid nodule formation but also with the progression of non-solid to solid adenocarcinoma. Such immunosuppressive phenotypes have been reported recently in subsolid nodules*^16^*. Similarly, a progressive decrease in tissue-resident alveolar macrophages was associated with a concomitant increase in anti-inflammatory/immunosuppressive M2-like monocyte-derived macrophages as a function of disease progression. There was an increased abundance of DCs with enhanced antigen presentation capacity, perhaps due to the greater prevalence of tumor neoantigens as a function of disease progression*^82^*. Spatial analysis showed that the increased abundance of DCs was actually in the TLS, whereas in the non-TLS regions, there was decreased proximity of DCs to tumor cells in the solid component compared to non-solid. Additionally, we observed a decreased proportion of mregDCs in the non-solid and solid components, which is unexpected, as mregDCs were shown to limit the activity of CD8 T and NK cell anti-tumor phenotypes in NSCLC*^122^*.

The stromal fraction showed a vascular compartment with increased angiogenic activity and reduced immunomodulatory function. Additionally, there was increased expression of α-SMA, *COL4A1* and *MYH11* in pericyte/smooth muscle cells, markers previously implicated in T-cell exclusion by cancer-associated fibroblasts (CAFs) in early-stage lung adenocarcinoma*^89^*. This is consistent with our previous finding that non-solid nodules harbor αSMA-positive CAFs organized in a continuous layer circumscribing the neoplastic epithelial cells and that this CAF morphology is associated with T cell exclusion from the cancer cell nests*^96^*. Future studies are likely to characterize the mechanisms underlying CAF-mediated T- cell exclusion and immunosuppression in non-solid nodules. Importantly, many of the immunosuppressive mechanisms observed in the non-solid components were further augmented in the solid component.

Cell-cell communication analysis identified complex dynamics of angiogenesis modulation and immunosuppression, and together with spatial analysis revealed how the immunosuppressive mechanisms operate within the heterogeneous TME to drive tumor progression and immune regulation. Particularly, there was an increased density of suppressive immune cells and a decreased density of cytotoxic T cells around the tumor cells. Together, these mechanisms are likely to contribute to tumor initiation and subsequent progression by conferring resistance to immune-mediated clearance. TLSs serve as an important component of the immune response occurring in TME and are associated with better prognosis and immunotherapy efficacy. However, recent studies suggest that the effects of TLS on immune control may depend on its cellular composition, maturation state and spatial location (peritumor vs intratumor)*^123^*. Our analysis of TLS from discrete non-solid and solid components showed that disease initiation and subsequent progression were associated with increased exhaustion of both CD8 and CD4 T cells in TLS. This exhaustion was accompanied by increased expression of CXCL13 by the exhausted T cells, which is believed to promote the recruitment of B cells and facilitate the formation of TLSs. While CD8 T cells in lung cancer are reported to preferentially localize in TLSs, which provides a protective niche for CD8 T cells to exert an antitumor effect*^124^*, our study highlights the need to identify mechanisms that blunt T cell activity in the TLS to support tumor growth. Importantly, we identified BET triads as a new three-cell-cluster comprised of B cell, EC and T cell (BET triads), with increased frequency in the non-solid compared to the solid component. We posit that the BET triads in the TLS represent high endothelial venules (HEVs) recruiting B and T cells from systemic circulation and supporting TLS function in the context of T cell activation and antigen presentation.

A major challenge in the clinical management of patients with preinvasive nodules is a lack of reliable biomarkers for nodule progression prediction. Within the Treg population, we identified a unique subset of tumor-infiltrating *TNFRSF9* (*4-1BB*) Tregs*^49^* that increased in the non-solid and solid lesions. Notably, a 260 gene signature from *TNFRSF9*^high^ Tregs was predictive of worse overall survival in TCGA LUAD*^50^*. Tregs also expressed *LAIR2*, a secreted receptor that binds collagen and was found to be adversely prognostic in LUAD*^125^*. These findings suggest that these candidates have the potential to be developed as prognostic biomarkers for the progression of non-solid to invasive adenocarcinoma and may advocate for nodule biopsies.

There is much interest in intercepting the progression of preinvasive nodules to malignant disease. Minimally invasive surgery used in patients with solitary nodules confers superior outcomes. However, nearly 50% of patients presenting with synchronous multifocal ground-glass nodules*^126^* on their initial CT may not qualify for surgery and, therefore, likely to benefit from clinically actionable targets for disease interception. Targets for therapy development include pre-exhausted tumor-infiltrating CD8 T cells expressing inhibitory receptor *NKG2A.* NKG2A+ CD8 T cells abundantly present in lung tumors are associated with a poorer prognosis and response to immunotherapy, and blocking the inhibitory NKG2A receptor was shown to reactivate CD8 T cells and improve the impact of immunotherapies, including cancer vaccines*^127^*. We showed that tumor-infiltrating NK cells have distinct functional roles in tumor immunity, including direct cytotoxic effects on malignant cells and the recruitment of key immune cell populations. Notably, through secretion of the chemokines XCL1, XCL2 and CCL5, NK cells attract cross-presenting XCR1^high^ cDC1s to the TME, which are essential for T cell-mediated immunity. Importantly, a variety of approaches can be considered to enhance NK cell-mediated tumor immunity in preinvasive disease, including antibodies that block inhibitory receptors (NKG2A) or to prevent MICA/B proteolytic shedding by tumor cells*^128,129^*. Additionally, the suppressive tumor-infiltrating 4-1BB+ Treg population can be preferentially depleted by an FcyR-optimized anti-4-1BB monoclonal antibody *in vivo* to promote effector T cell agonism leading to marked tumor control*^130,131^*. Given that intratumoral Tregs have been shown to express CCR8*^133^*, another approach is to consider the Fc-optimized anti-CCR8 antibody, which was shown in murine models to selectively deplete intratumoral Tregs and not peripheral Tregs, resulting in impaired tumor growth*^132,133^*. Similarly, we identified *TREM2* expressing M2-like macrophages, which constrain NK and T cell function*^74^*. Targeting TREM2^high^ macrophages is being developed as a therapeutic modality. Fc domain enhanced anti-TREM2 monoclonal antibody was shown to impair tumor growth in mice by enhancing anti-tumor CD8 T cell responses and altering the balance between immunosuppressive and immunostimulatory macrophages*^134^*. Indeed, a humanized anti-TREM2 mAb currently is being tested in humans (NCT04691375) and may be considered in future clinical trials seeking to intercept preinvasive to invasive disease progression. We also observed ECs with increased angiogenic activity driven by tumor-mediated VEGF-VEGFR activated pathway network involving *PVLAP* and *ANGPT2* in the endothelium with a concomitant reduction in immune activity during non-solid to solid disease progression. Inhibition of *PLVAP* and *ANGPT2* has been shown to suppress tumor growth in mice*^135^*, suggesting the potential of FDA-approved antiangiogenic therapies, including anti-VEGF Avastin and anti-ANGPT2 Vabysmo, to intercept preinvasive to invasive disease progression. In summary, this study identifies determinants of early disease emergence and progression, with potential for development not only as diagnostic/prognostic biomarkers but also as targets for disease interception.

## Methods

### Clinical samples collection and processing

The samples for this Institutional Review Board approved study (#1008011221) were collected at Weill Cornell using the Thoracic Surgery Biobank Protocol. Two separate cohorts of patients were used. *1)* For sc-RNAseq, prospectively collected freshly resected CT-imaged part solid nodules were immediately microdissected into adjacent normal lung, non-solid/preinvasive and solid /invasive components by a trained pulmonary pathologist, and the remaining tissue was storage as formalin-fixed paraffin-embedded tissue. CT images were reviewed by 2 trained radiologists and classified as part-solid or solid nodules on CT attenuation *2)* For validation, bulk RNA-seq data from the Weill Cornell lung nodule cohort, a retrospective collection of CT-imaged nodules was used.

### Preparation of single-cell RNA-sequencing libraries

Single-cell suspension was prepared by the genomics core facility at Weill Cornell following protocols from 10X Genomics. Single-cell libraries were generated using 10X Genomics Chromium Single Cell 2’ Reagent Kits User Guide (v3.1 Chemistry Dual Index) and sequenced on the NovaSeq 6000 sequencing system (Illumina).

### Single-cell RNA-sequencing data pre-processing and quality control

Sequencing reads were aligned to the human transcriptome (GRCh38) using cell ranger count (10x Genomics Cell Ranger v6.0*^136^*) with default parameters. The method decontX was applied to remove RNA contamination*^137^* and a Seurat object was created with the filtered counts (Seurat v4.1.0, SeuratObject v4.0.4)*^138^*. Low-quality cells (i.e., cells with less than 200 or more than 6500 genes and cells with more than 15% mitochondrial genes) were removed. Highly expressed genes (i.e., *MALAT1* and mitochondrial genes) were removed. Doublets were identified and removed using the DoubletFinder R packages (v2.0.3)*^139^*. Cells expressing multiple cell lineage markers were considered doublets as well and filtered out (epithelial: *SFTPC*, *SCGB3A1*, lymphoid: *CD3D*, *CD79A*, *JCHAIN*, *MZB1*, myeloid: *MARCO*, *CD163*, pan-immune cell: *PTPRC*).

### Sample normalization and integration

Samples were merged and the scTransform method was applied with ‘method=” glmGamPoi”’ for normalization, scaling, and variable features identification. Cell-cycle and technical-noise-associated genes (*XIST* and *NEAT1*) were removed from the variable genes list. Principal-component analysis (PCA) was performed, and the first 20 principal components were selected using an elbow plot for integration via Harmony *^140^*.

### Clustering and identification of major cell lineages

The integrated Seurat object was reclustered based on 20 dimensions of the harmony reduction using the ‘FindNeighbors’ and ‘FindClusters’ methods with ‘resolution = 0.3’. A Umap was then created based on the same 20 Harmony dimensions using the ‘RunUMAP’ method. The clusters were then explored manually and were annotated based on known marker expression as epithelial (*EPCAM*), endothelial (*CDH5, VWF*), stroma (*FBN1, COL1A1*), lymphoid (all: *PTPRC*; T: *CD3D, CD3G*’; B/plasma: *CD79A*, *MS4A1*, *JCHAIN*; NK: *GNLY*, *NKG7*) and myeloid cells (all: *PTPRC*; macrophage: *CD68*, *CD163*; monocyte: *CD14*, *FCGR3A*, neutrophils: *S100A8*, *FCGR3B*; dendritic: *FCGR2B*, *HLA-DPB1*).

### Reclustering of identified cell types

The integrated object was separated into five cell lineage-specific objects (i.e., myeloid, lymphoid, epithelial, endothelial and stromal), renormalized, and reclustered in the same manner described above. Cells that were misannotated and added to the incorrect cell lineage object were identified based on the expression of the makers mentioned above and removed iteratively (i.e., clustering, identifying contaminant clusters, eliminating those clusters and reclustering). Cluster markers were identified using Seurat’s FindAllMarkers Method. The final clusters were annotated based on the expression of known cell-type markers curated from the literature (**Supplementary Table S2**) and compared to the identified cluster markers.

### Tumor cell identification

We applied InferCNV*^141^* on epithelial cells from solid and non-solid nodule samples using parameters ‘cutoff=0.1, BayesMaxPNormal = 0.2, analysis_mode = “subclusters”, tumor_subcluster_partition_method = “qnorm, denoise = T”. Epithelial cells from adjacent normal tissue samples served as the reference. We then calculated a CNV score per cell following a procedure described by Tirosh *et al^142^*. Arm-level CNV scores were determined by averaging the squares of CNV values for each chromosomal arm, and then aggregating these scores by taking their arithmetic mean across all arms. Clusters with elevated CNV scores were identified as tumor clusters. Additional evidence supporting the tumor designation included the enrichment of nodule samples (both solid and non-solid) in these clusters and the cells either not expressing known lung epithelial cell markers at all or simultaneously expressing markers from several cell types (especially AT2 and club cells).

### Trajectory analysis

Monocle 2 (v2.22.0)*^37^* was used to reconstruct the differentiation trajectories of epithelial, CD8 T, and Treg cells. Seurat objects were transformed into CellDataSet objects that were projected into a lower dimensional space using the ‘reduceDimension’ method and the DDRTree algorithm.

### Quantifying regulon activity

To compare the TF activity of Treg and endothelial subpopulations, we applied the SCENIC method (v1.3.1)*^143^* that reconstructs gene regulatory networks based on gene co-expression and DNA motif analysis. That included inferring potential TF targets based on the expression data using GRNBoost, selecting potential direct-binding targets (regulons) based on DNA-motif analysis (RcisTarget: TF motif analysis), and scoring the regulons in the cells using AUCell. The hg38 - refseq_r80 data files (‘hg38_10kbp_up_10kbp_down_full_tx_v10_clust. genes_vs_motifs.rankings.feather’, and ‘hg38_500bp_up_100bp_down_full_tx_v10_clust.genes_vs_motifs.rankings.feather’) were downloaded from the cisTarget resources website (https://resources.aertslab.org/cistarget/).

### Ranking cell-cell communications

The MultiNicheNet framework from R package multinichenetr (v2.0.0)*^144^* to analyze and prioritize potential differential cell-cell communication (CCC) events. This analysis incorporated ligand-receptor interactions and their subsequent signaling to target genes. The pre-calculated ligand-receptor network (lr_network_human_21122021.rds) and ligand-target matrix (ligand_target_matrix_nsga2r_final.rds) were retrieved from https://zenodo.org/records/7074291. We conducted two tests: non-solid vs. normal and solid vs. non-solid with default settings. We examined the overall top twenty CCCs. Circos plots were generated using the ‘make_circos_group_comparison’ function in the multinichenetr package.

### Metabolic activity quantification

We quantified the metabolic activity of Tregs using the scMetabolism R package*^145^* and the KEGG*^146^* and REACTOME*^147^* databases. The ‘sc. metabolism.Seurat’ function was employed with its default settings.

### Gene signature score calculation

We utilized the AddModuleScore method from the Seurat R package*^138^* to compute various cellular scores: AT2-like and club-like scores for epithelial cells; cytotoxicity, exhaustion, naïve and TRM scores for T cells; suppressive score for Tregs; cytotoxicity and chemokine-expression scores for NK cells; inflammatory, immunosuppressive and antigen-presenting scores for dendritic cells; immunosuppressive (M2-like), proinflammatory (M1-like), alveolar macrophage, and monocyte-derived scores for macrophages; and scores for immature neutrophils in myeloid cells. The signatures employed for these calculations are depicted in **Supplementary Table S4**.

### Differential expression and gene-set enrichment analysis

The method FindMarkers from the Seurat R package*^138^* was used to test genes for differential expression with ‘min.pct=0.25’. For differential gene expression and gene signature score tests between matched patient samples (e.g., solid vs non-solid), the logistic regression DE test with the patient ID as a latent variable was used. For unpaired samples (e.g., EGFR-pos vs EGFR-neg samples), we used the default Wilcoxon Rank Sum test. For gene set enrichment analysis (GSEA) the threshold for log fold change was set to 0 and for all other tests it was set to 0.25.

### Bulk RNA-sequencing

RNA was extracted from FFPE samples and processed as described*^17^*. After post-capture PCR amplification and purification, the quality of the final libraries was checked and loaded to Illumina NovaSeq6000 for sequencing at PE2×100 cycles. The raw sequencing reads in BCL format were processed through bcl2fastq 2.19 (Illumina) for FASTQ conversion and demultiplexing. All reads were aligned to the human reference genome GRCh38 using STAR aligner (version STAR-2.7.10a)*^148^* with GENCODE (release GRCh38.p14.v44)*^149^*. The aligned reads were subsequently quantified as fragments per kilobase of transcript per million mapped reads (FPKMs) using Cufflinks (version-2.2.1)*^150^*. The bulk expression profiles were used to deconvolve the TME into 64 distinct cell types using xCell*^151^*. Following deconvolution, the results were subset to focus specifically on immune cells and stromal cells. Among these, cell types with statistically significant differences were identified using the Wilcoxon rank-sum test (p-value < 0.05). These selected cell types were visualized through box plots to illustrate the distribution of expression across samples. The TME was also characterized using single-sample Gene Set Enrichment Analysis (ssGSEA)*^152^* implemented using the GSVA package (version 1.46.0) in R (version 4.2.2), with cell type signatures inferred from scRNA-seq profiles for the GGOs.

### Imaging mass cytometry (IMC)

*<u>Tissue Deparaffinization and Antigen Retrieval</u>* FFPE tissue samples were sectioned at a 5-μm thickness for IMC. Prior to staining, paraffinized tissue slides were incubated on a slide warmer set at 65°C for 2 hours. pH9 Tris-ETDA buffer antigen retrieval solution was prepared and aliquoted in a 50mL Falcon tube at a volume of 40mL to preheat in a water-bath set to 95°C. After incubation on the slide warmer, the slides were deparaffinized in 100% CitriSolv, 2X for 10 minutes each with gentle agitation every 3 minutes, then underwent rehydration in a descending series of ethanol solutions: 2X in 100%, 1X in 80%, 1X in 70%; all for 5 minutes each, with gentle agitation every 2 minutes. After the ethanol series, the slides were washed in MilliQ water for 5 minutes, with gentle agitation every 2 minutes. Two slides were placed in a back-to-back configuration in the 95°C preheated tube to incubate in the water-bath for 30 minutes. After removal from the bath to cool at room temperature for 15 minutes, the slides were removed from the antigen retrieval solution and washed in 1X TBS on a shaker 2X for 10 minutes each. *<u>Tissue Staining</u>*. The slides were transferred to a hydration chamber, with brief drying around the tissue. A PAP hydrophobic pen was used to closely outline each tissue core. SuperBlock (PBS) Blocking Buffer was applied to the tissue regions, and the slides were incubated in the hydration chamber at RT for 1 hour. Prior to experimentation, control testing of dilutions of all antibodies was performed to optimize the antibody panel and protocol. The antibody cocktail was made with the respective conjugated antibody dilutions, 1X PBS, and 10% BSA volumes. The concentration of the antibody cocktail used was 1:100, and of each antibody, 0.5mg/mL. After removing the blocking solution, the antibody mix was applied to the tissue sections and incubated overnight at 4°C in the hydration chamber. The following day, the slides were washed in 0.2% Triton X in 1X PBS twice for 10 minutes each with agitation on the shaker, followed by two washes in 1X TBS for 10 minutes each. The tissues were then stained with iridium-intercalator solution, made 1:300 in 1X PBS, to incubate for 30 min at RT in the hydration chamber. The slides were washed for 5 minutes in 1X TBS, followed by another 5-minute wash in MilliQ water. After washing, the slides were left to dry in open air at RT for at least 20 minutes before subjection to Hyperion acquisition. *<u>Hyperion Image</u> <u>Acquisition.</u>* Upon successful completion of tuning and quality control, one image per slide was acquired using the Hyperion. The slides were scanned at 200 Hz in the Hyperion Imaging System (Standard BioTools). Panoramic images were created to encompass the region of tissue that was readable by the CyTOF software, and regions of interest were selected in accordance with H&E-stained images of the respective tissues. The antibody panel with its respective conjugated metals was imported and selected for each ROI before commencing ablation and raw data acquisition.

### Characterization of cell phenotypes and cell density analysis in IMC

For cellular phenotype characterization and cell density analysis, raw data, stored as .mcd files, were transformed into TIFF format using the IMC Python Package (https://github.com/ElementoLab/imc). The Ilastik model was used to predict cellular locations and boundaries via pixel and object classification*^153^*. Single-cell segmentation for each image was achieved using the DeepCell package*^154^*. Nuclear analysis was integrated with cell boundary data to determine cell segmentation. The mean expressions of panels from segmented single cells were retrieved using the IMC package by overlaying segmentation masks onto the corresponding TIFF images. Resulting segmented images and data files were cataloged into CSV and AnnData formats. For enhanced precision in cell protein expression metrics, we employed the PowerTransform (Yeo-Johnson) normalization tools designed for skewed and bimodal condition interpretations. This normalization was executed for each region of interest and individual sample. To compare the characteristics of cellular phenotypes, CD4 or CD8 T cells were isolated from TLSs, or regions devoid of these structures. The IMC images were transposed into topological neighborhood graphs, wherein cells were depicted as nodes. Direct cell-cell neighboring pairs, within a 100 µm range between centroids, were depicted as edges. The quantification of neighborhoods is predicated upon the cellular density surrounding the center of cancer cells, facilitating comparisons between grouped data sets.

### B cell-Endothelium-T cell (BET) TRIAD Analysis from IMC

A new triad composed of B cells, endothelium, and T cells was proposed in this study. Its quantification, based on IMC, was performed as previously described*^155^*. Briefly, endothelial cells (ECs) were identified by the expression of CD31+Pan- CK- markers, while B cells were identified as CD20+HLA-DR+. T cells were categorized into CD4 T cells (CD3+CD4+) and CD8 T cells (CD3+CD8). For each EC, an EC-centric boundary was established based on its X and Y coordinates and the major axis length (the longest diameter of the EC). This boundary defined the spatial extent of interactions by determining which B or T cells were close enough to be considered in contact with the EC. Each cell was assigned a unique identifier for tracking within the dataset. Next, the distances between ECs and B or T cells were calculated using their X and Y spatial coordinates. The distance between the EC and the lymphocytes was checked to confirm if it fell within 2X the major axis length of the EC. Only interactions within this boundary were included in the analysis. When a B cell was in contact with an EC, the interaction was labeled as contact B, and similarly, contact T was used when a T cell was in contact with an EC. A BET TRIAD was defined when both B cells and T cells were in contact with the same EC. Data entries with only ‘single contact’ or ‘no contact’ between ECs and B or T cells were removed, ensuring that only relevant interactions were analyzed. Once contacts were calculated and interactions classified, a result table was generated. The distances between ECs and lymphocytes (both B and T cells) were recalculated and verified against the major axis length to ensure accuracy.

### Statistical analysis for IMC

The details of the statistical analyses performed for each experiment are provided in the respective figure legends. To compare continuous variables between groups, normality was tested using Levene’s test for equality of variance. If the p-value for normality was less than 0.05, the variables were considered non-parametric, and a Mann-Whitney U test (Wilcoxon rank sum) was performed. If the p-value was greater than 0.05, the variables were considered parametric, and a Student’s t-test was applied. Data are presented as mean ± SD. A two-sided p-value < 0.05 was considered statistically significant.

### 10X Visium Spatial transcriptomics sequencing

Library preparation for CytAssist Spatial Gene Expression, sequencing and post-processing of the raw data was performed at the Epigenomics Core at Weill Cornell Medicine. Spatial gene expression libraries were prepared using the 10x Genomics CytAssist platform (10x Genomics, Pleasanton, CA). FFPE tissue sections from eight tissue blocks were sectioned at 5 μm thickness, placed on histology slides, dried at 42°C for 3 hours, and stored in a desiccator overnight at room temperature before proceeding with the CytAssist Spatial Gene Expression protocol (guides CG000520 and CG000495). The deparaffinized and H&E-stained tissue slides were then imaged using an EVOS M7000 Automated Imaging System (10x objective, 3.45 µm/pixel - Thermo Fisher Scientific, CA). Following imaging, the slides were de-coverslipped and assembled in the 10x Tissue Slide Cassette, ensuring that the tumor areas were positioned within the 6.5 mm x 6.5 mm capture area. The tissue sections were subsequently destained, de-crosslinked, and hybridized overnight with the Visium Human Transcriptome Probe Set v2.0 (PN-1000466), which contains probe pairs specific to each of the 18,049 targeted genes. After hybridization, the probe pairs were ligated, and the slides were loaded onto the Visium CytAssist instrument, where the regions of interest were adjusted, and ligated probes were transferred and captured on the spatially barcoded oligos of the Visium CytAssist SD slide. Full-length probes were eluted from the Visium SD slide and pre-amplified through 10 cycles of PCR before generating indexed libraries. The libraries were pooled at 10 nM concentration, clustered on an Illumina NovaSeq 6000 paired-end read flow cell and sequenced with 28 cycles on Read 1 (10x barcode and UMI), 10 cycles for the I7 Index (sample index), and 50 bases on Read 2 (probe), achieving a coverage of approximately 150 million reads per sample. Primary processing of the sequencing images was performed using Illumina’s Real-Time Analysis (RTA) software. The 10x Genomics Space Ranger software (v 2.0.0) was subsequently used to demultiplex the data, align reads to the human genome (hg38), assign reads to their original spatial location using the spatial barcodes, quantify transcripts using Unique Molecular Identifiers (UMIs), generate a gene expression matrix, and integrate the spatial gene expression data with the tissue morphology.

### 10X Visium data processing and deconvolution

Visium spots were deconvoluted via the BayesTME*^156^* method using the scRNA-seq data as the cell type expression guidance. The scRNA-seq gene expression was averaged per cell type and the standard preprocessing steps specified in BayesTME were used. Spatial significance scores were calculated by taking the spatial standard deviation of each gene. Genes with the top spatial significance score (n=5000) were picked as the candidate spatial genes for each sample. The intersection of the picked candidate spatial genes of all 8 samples (n=3764) was used for the cohort spatial analysis and cell type deconvolution. BayesTME model was trained with the cohort of 8 patient samples jointly and devolved the aggregated spatial transcriptomics signals into 22 cell types of interest. Based on the cell type deconvolution results, spatial coexistence scores were calculated using Pearson’s r correlation on the cell counts of each cell type. To evaluate gene signature enrichment AUCell scores*^143^* were calculated per Visium spot based on the gene signatures of exhausted T cells (*LAG3*, *PDCD1*, *CTLA4*, *HAVCR2*, *TIGIT*, *TOX*, *LAYN* and *CXCL13*) and B cells (*CD19*, *MS4A1*, *CD79A* and *CD79B*).

## Acknowledgments

We thank Cathy Spinelli, Joyce Gakuria, Abu Nasar, and Murtaza Malbari for clinical support. We thank Dr. Jenny Xiang of the Genomics Resources Core Facility for professional advice. This work was supported in part by NIH grant UH3CA244697, and funds from The Neuberger Berman Foundation Lung Cancer Research Center; generous gifts/donations from Jay and Vicky Furman; Patricia Donnington Research Fund, and patients in the Division of Thoracic Surgery to N.K.A. H-S.L was supported by US Department of Defense Impact Award (W81XWH-22-1-0657), Cancer Prevention and Research Institute of Texas grant (CPRIT RP200443), NIH R21 (R21AI159379), and the Helis Medical Research Foundation. The funding organizations played no role in experimental design, data analysis or manuscript preparation.

## Author contributions

V.M., and N.K.A. supervised this study. L.Y., N.K.A., and V.M. designed the experiments and wrote the manuscript. L.Y. B.B, S.W.K, H.Z, W.T, H-S.L and J.K performed computational analysis. A.B and C.Z performed histopathological analysis and microdissection of part-solid nodules. A.S. coordinated the clinical sample acquisition and biobanking. J.K, T.M, O.E, A.B, G.J.M, M.M provided suggestions and edited the manuscript. All authors discussed the results and conclusions drawn from the studies.

## References

1. Pao W, Girard N. New driver mutations in non-small-cell lung cancer. Lancet Oncol. Feb 2011;12(2):175–80. doi:10.1016/S1470-2045(10)70087-5

2. Network CGAR. Comprehensive molecular profiling of lung adenocarcinoma. Nature. Jul 2014;511(7511):543–50. doi:10.1038/nature13385

3. Aberle DR, Adams AM, Berg CD, et al. Reduced lung-cancer mortality with low-dose computed tomographic screening. N Engl J Med. Aug 2011;365(5):395–409. doi:10.1056/NEJMoa1102873

4. Henschke CI, Yankelevitz DF, Mirtcheva R, et al. CT screening for lung cancer: frequency and significance of part-solid and nonsolid nodules. AJR Am J Roentgenol. May 2002;178(5):1053–7. doi:10.2214/ajr.178.5.1781053

5. Aokage K, Saji H, Suzuki K, et al. A non-randomized confirmatory trial of segmentectomy for clinical T1N0 lung cancer with dominant ground glass opacity based on thin-section computed tomography (JCOG1211). Gen Thorac Cardiovasc Surg. May 2017;65(5):267–272. doi:10.1007/s11748-016-0741-1

6. Kobayashi Y, Mitsudomi T. Management of ground-glass opacities: should all pulmonary lesions with ground-glass opacity be surgically resected? Transl Lung Cancer Res. Oct 2013;2(5):354–63. doi:10.3978/j.issn.2218-6751.2013.09.03

7. Sivakumar S, Lucas FAS, McDowell TL, et al. Genomic Landscape of Atypical Adenomatous Hyperplasia Reveals Divergent Modes to Lung Adenocarcinoma. Cancer Res. 11 15 2017;77(22):6119–6130. doi:10.1158/0008-5472.CAN-17-1605

8. Hu X, Fujimoto J, Ying L, et al. Multi-region exome sequencing reveals genomic evolution from preneoplasia to lung adenocarcinoma. Nat Commun. 07 05 2019;10(1):2978. doi:10.1038/s41467-019-10877-8

9. Hu X, Estecio MR, Chen R, et al. Evolution of DNA methylome from precancerous lesions to invasive lung adenocarcinomas. Nat Commun. 01 29 2021;12(1):687. doi:10.1038/s41467-021-20907-z

10. Chen H, Carrot-Zhang J, Zhao Y, et al. Genomic and immune profiling of pre-invasive lung adenocarcinoma. Nat Commun. Nov 29 2019;10(1):5472. doi:10.1038/s41467-019-13460-3

11. Wang S, Du M, Zhang J, et al. Tumor evolutionary trajectories during the acquisition of invasiveness in early stage lung adenocarcinoma. Nat Commun. 11 27 2020;11(1):6083. doi:10.1038/s41467-020-19855-x

12. Teixeira VH, Pipinikas CP, Pennycuick A, et al. Deciphering the genomic, epigenomic, and transcriptomic landscapes of pre-invasive lung cancer lesions. Nat Med. Mar 2019;25(3):517–525. doi:10.1038/s41591-018-0323-0

13. Kadara H, Choi M, Zhang J, et al. Whole-exome sequencing and immune profiling of early-stage lung adenocarcinoma with fully annotated clinical follow-up. Ann Oncol. Jan 01 2017;28(1):75–82. doi:10.1093/annonc/mdw436

14. Dejima H, Hu X, Chen R, et al. Immune evolution from preneoplasia to invasive lung adenocarcinomas and underlying molecular features. Nat Commun. 05 11 2021;12(1):2722. doi:10.1038/s41467-021-22890-x

15. Krysan K, Tran LM, Grimes BS, et al. The Immune Contexture Associates with the Genomic Landscape in Lung Adenomatous Premalignancy. Cancer Res. Oct 01 2019;79(19):5022–5033. doi:10.1158/0008-5472.CAN-19-0153

16. Yanagawa J, Tran LM, Salehi-Rad R, et al. Single-cell characterization of pulmonary nodules implicates suppression of immunosurveillance across early stages of lung adenocarcinoma. Cancer Res. Jul 21 2023;doi:10.1158/0008-5472.CAN-23-0128

17. Altorki NK, Borczuk AC, Harrison S, et al. Global evolution of the tumor microenvironment associated with progression from preinvasive invasive to invasive human lung adenocarcinoma. Cell Rep. 04 05 2022;39(1):110639. doi:10.1016/j.celrep.2022.110639

18. Chen Y, Toth R, Chocarro S, et al. Club cells employ regeneration mechanisms during lung tumorigenesis. Nat Commun. Aug 05 2022;13(1):4557. doi:10.1038/s41467-022-32052-2

19. Meyer N, Penn LZ. Reflecting on 25 years with MYC. Nature Reviews Cancer. 2008/12/01 2008;8(12):976–990. doi:10.1038/nrc2231

20. Zhou D, Duan Z, Li Z, Ge F, Wei R, Kong L. The significance of glycolysis in tumor progression and its relationship with the tumor microenvironment. Front Pharmacol. 2022;13:1091779. doi:10.3389/fphar.2022.1091779

21. Mehner C, Radisky ES. Bad Tumors Made Worse: SPINK1. Opinion. Frontiers in Cell and Developmental Biology. 2019-February-04 2019;710.3389/fcell.2019.00010

22. Filippou PS, Karagiannis GS, Constantinidou A. Midkine (MDK) growth factor: a key player in cancer progression and a promising therapeutic target. Oncogene. 2020/03/01 2020;39(10):2040–2054. doi:10.1038/s41388-019-1124-8

23. Lin G, Chen L, Lin L, et al. Comprehensive Analysis of Aquaporin Superfamily in Lung Adenocarcinoma. Original Research. Frontiers in Molecular Biosciences. 2021-October-11 2021;8doi:10.3389/fmolb.2021.736367

24. Burgos M, Cavero-Redondo I, Álvarez-Bueno C, et al. Prognostic value of the immune target CEACAM6 in cancer: a meta-analysis. Ther Adv Med Oncol. 2022;14:17588359211072621. doi:10.1177/17588359211072621

25. Ferone G, Lee MC, Sage J, Berns A. Cells of origin of lung cancers: lessons from mouse studies. Genes Dev. Aug 1 2020;34(15-16):1017–1032. doi:10.1101/gad.338228.120

26. Li X, Sun X, Kan C, et al. COL1A1: A novel oncogenic gene and therapeutic target in malignancies. Pathology - Research and Practice. 2022/08/01/ 2022;236:154013. 10.1016/j.prp.2022.154013

27. Hou L, Lin T, Wang Y, Liu B, Wang M. Collagen type 1 alpha 1 chain is a novel predictive biomarker of poor progression-free survival and chemoresistance in metastatic lung cancer. J Cancer. 2021;12(19):5723–5731. doi:10.7150/jca.59723

28. Zhao Y, Lu H, Yan A, et al. ABCC3 as a marker for multidrug resistance in non-small cell lung cancer. Sci Rep. Nov 1 2013;3:3120. doi:10.1038/srep03120

29. Meng Y, Wang L, Chen D, et al. LAPTM4B: an oncogene in various solid tumors and its functions. Oncogene. Dec 15 2016;35(50):6359–6365. doi:10.1038/onc.2016.189

30. Chen HZ, Tsai SY, Leone G. Emerging roles of E2Fs in cancer: an exit from cell cycle control. Nat Rev Cancer. Nov 2009;9(11):785–97. doi:10.1038/nrc2696

31. Vousden KH, Lu X. Live or let die: the cell’s response to p53. Nature Reviews Cancer. 2002/08/01 2002;2(8):594–604. doi:10.1038/nrc864

32. Li Y, Miao L, Yu M, et al. α1-antitrypsin promotes lung adenocarcinoma metastasis through upregulating fibronectin expression. Int J Oncol. 2017/06/01 2017;50(6):1955–1964. doi:10.3892/ijo.2017.3962

33. Chang TM, Chiang YC, Lee CW, et al. CXCL14 promotes metastasis of non-small cell lung cancer through ACKR2-depended signaling pathway. Int J Biol Sci. 2023;19(5):1455–1470. doi:10.7150/ijbs.79438

34. Wang Y, Qiu S, Wang H, et al. Transcriptional Repression of Ferritin Light Chain Increases Ferroptosis Sensitivity in Lung Adenocarcinoma. Original Research. Frontiers in Cell and Developmental Biology. 2021-October-26 2021;9doi:10.3389/fcell.2021.719187

35. Yang Y, Zhang X, Gao Y, et al. Research Progress in Immunotherapy of NSCLC With EGFR-Sensitive Mutations. Oncol Res. May 4 2022;29(1):63–74. doi:10.3727/096504022x16462176651719

36. Hamarsheh Sa, Groß O, Brummer T, Zeiser R. Immune modulatory effects of oncogenic KRAS in cancer. Nature Communications. 2020/10/28 2020;11(1):5439. doi:10.1038/s41467-020-19288-6

37. Cao J, Spielmann M, Qiu X, et al. The single-cell transcriptional landscape of mammalian organogenesis. Nature. 2019;566(7745):496–502.

38. van der Leun AM, Thommen DS, Schumacher TN. CD8+ T cell states in human cancer: insights from single-cell analysis. Nature Reviews Cancer. 2020/04/01 2020;20(4):218–232. doi:10.1038/s41568-019-0235-4

39. Parry EM, Lemvigh CK, Deng S, et al. ZNF683 marks a CD8. Cancer Cell. Oct 09 2023;41(10):1803–1816.e8. doi:10.1016/j.ccell.2023.08.013

40. Chen Y, Xin Z, Huang L, et al. CD8(+) T Cells Form the Predominant Subset of NKG2A(+) Cells in Human Lung Cancer. Front Immunol. 2019;10:3002. doi:10.3389/fimmu.2019.03002

41. Guo X, Zhang Y, Zheng L, et al. Global characterization of T cells in non-small-cell lung cancer by single-cell sequencing. Nature Medicine. 2018/07/01 2018;24(7):978–985. doi:10.1038/s41591-018-0045-3

42. Wherry EJ, Kurachi M. Molecular and cellular insights into T cell exhaustion. Nat Rev Immunol. Aug 2015;15(8):486–99. doi:10.1038/nri3862

43. Cardenas MA, Prokhnevska N, Sobierajska E, et al. Differentiation fate of a stem-like CD4 T cell controls immunity to cancer. Nature. Oct 23 2024;doi:10.1038/s41586-024-08076-7

44. Tanemoto S, Sujino T, Miyamoto K, et al. Single-cell transcriptomics of human gut T cells identifies cytotoxic CD4+CD8A+ T cells related to mouse CD4 cytotoxic T cells. Original Research. Frontiers in Immunology. 2022-October-24 2022;13doi:10.3389/fimmu.2022.977117

45. Clarke J, Panwar B, Madrigal A, et al. Single-cell transcriptomic analysis of tissue-resident memory T cells in human lung cancer. J Exp Med. Sep 2 2019;216(9):2128–2149. doi:10.1084/jem.20190249

46. Toribio-Fernández R, Herrero-Fernandez B, Zorita V, et al. Lamin A/C deficiency in CD4+ T-cells enhances regulatory T-cells and prevents inflammatory bowel disease. The Journal of Pathology. 2019;249(4):509–522. 10.1002/path.5332

47. Chinen T, Kannan AK, Levine AG, et al. An essential role for the IL-2 receptor in T. Nat Immunol. Nov 2016;17(11):1322–1333. doi:10.1038/ni.3540

48. Burchill MA, Yang J, Vogtenhuber C, Blazar BR, Farrar MA. IL-2 receptor beta-dependent STAT5 activation is required for the development of Foxp3+ regulatory T cells. J Immunol. Jan 01 2007;178(1):280–90. doi:10.4049/jimmunol.178.1.280

49. Bhairavabhotla R, Kim YC, Glass DD, et al. Transcriptome profiling of human FoxP3+ regulatory T cells. Hum Immunol. Feb 2016;77(2):201–13. doi:10.1016/j.humimm.2015.12.004

50. Guo X, Zhang Y, Zheng L, et al. Global characterization of T cells in non-small-cell lung cancer by single-cell sequencing. Nat Med. 07 2018;24(7):978–985. doi:10.1038/s41591-018-0045-3

51. Li L, Liu X, Sanders KL, et al. TLR8-Mediated Metabolic Control of Human Treg Function: A Mechanistic Target for Cancer Immunotherapy. Cell Metabolism. 2019/01/08/ 2019;29(1):103–123.e5. 10.1016/j.cmet.2018.09.020

52. Carriche GM, Almeida L, Stüve P, et al. Regulating T-cell differentiation through the polyamine spermidine. Journal of Allergy and Clinical Immunology. 2021/01/01/ 2021;147(1):335–348.e11. 10.1016/j.jaci.2020.04.037

53. Angelin A, Gil-de-Gómez L, Dahiya S, et al. Foxp3 Reprograms T Cell Metabolism to Function in Low-Glucose, High-Lactate Environments. Cell Metab. Jun 6 2017;25(6):1282–1293.e7. doi:10.1016/j.cmet.2016.12.018

54. Pacella I, Procaccini C, Focaccetti C, et al. Fatty acid metabolism complements glycolysis in the selective regulatory T cell expansion during tumor growth. Proc Natl Acad Sci U S A. Jul 10 2018;115(28):E6546–e6555. doi:10.1073/pnas.1720113115

55. Shi LZ, Wang R, Huang G, et al. HIF1alpha-dependent glycolytic pathway orchestrates a metabolic checkpoint for the differentiation of TH17 and Treg cells. J Exp Med. Jul 4 2011;208(7):1367–76. doi:10.1084/jem.20110278

56. Skadow M, Penna VR, Galant-Swafford J, Shevach EM, Thornton AM. Helios Deficiency Predisposes the Differentiation of CD4+Foxp3− T Cells into Peripherally Derived Regulatory T Cells. The Journal of Immunology. 2019;203(2):370–378. doi:10.4049/jimmunol.1900388

57. Wu SY, Fu T, Jiang YZ, Shao ZM. Natural killer cells in cancer biology and therapy. Mol Cancer. Aug 6 2020;19(1):120. doi:10.1186/s12943-020-01238-x

58. Tang F, Li J, Qi L, et al. A pan-cancer single-cell panorama of human natural killer cells. Cell. Sep 14 2023;186(19):4235–4251.e20. doi:10.1016/j.cell.2023.07.034

59. de Andrade LF, Lu Y, Luoma A, et al. Discovery of specialized NK cell populations infiltrating human melanoma metastases. JCI Insight. 12/05/ 2019;4(23)doi:10.1172/jci.insight.133103

60. Lugthart G, Melsen JE, Vervat C, et al. Human Lymphoid Tissues Harbor a Distinct CD69+CXCR6+ NK Cell Population. The Journal of Immunology. 2016;197(1):78–84. doi:10.4049/jimmunol.1502603

61. Marquardt N, Kekäläinen E, Chen P, et al. Unique transcriptional and protein-expression signature in human lung tissue-resident NK cells. Nature communications. 2019/08/26 2019;10(1):3841. doi:10.1038/s41467-019-11632-9

62. Marquardt N, Kekäläinen E, Chen P, et al. Human lung natural killer cells are predominantly comprised of highly differentiated hypofunctional CD69−CD56dim cells. Journal of Allergy and Clinical Immunology. 2017/04/01/ 2017;139(4):1321–1330.e4. 10.1016/j.jaci.2016.07.043

63. Böttcher JP, Bonavita E, Chakravarty P, et al. NK Cells Stimulate Recruitment of cDC1 into the Tumor Microenvironment Promoting Cancer Immune Control. Cell. 02 2018;172(5):1022–1037.e14. doi:10.1016/j.cell.2018.01.004

64. Gordon S, Plüddemann A, Martinez Estrada F. Macrophage heterogeneity in tissues: phenotypic diversity and functions. Immunol Rev. Nov 2014;262(1):36–55. doi:10.1111/imr.12223

65. Casanova-Acebes M, Dalla E, Leader AM, et al. Tissue-resident macrophages provide a pro-tumorigenic niche to early NSCLC cells. Nature. 2021/07/01 2021;595(7868):578–584. doi:10.1038/s41586-021-03651-8

66. Yang Q, Zhang H, Wei T, et al. Single-Cell RNA Sequencing Reveals the Heterogeneity of Tumor-Associated Macrophage in Non-Small Cell Lung Cancer and Differences Between Sexes. Original Research. Frontiers in Immunology. 2021-November-05 2021;12doi:10.3389/fimmu.2021.756722

67. Chistiakov DA, Myasoedova VA, Revin VV, Orekhov AN, Bobryshev YV. The impact of interferon-regulatory factors to macrophage differentiation and polarization into M1 and M2. Immunobiology. Jan 2018;223(1):101–111. doi:10.1016/j.imbio.2017.10.005

68. Tanaka T, Murakami K, Bando Y, Yoshida S. Interferon regulatory factor 7 participates in the M1-like microglial polarization switch. Glia. Apr 2015;63(4):595–610. doi:10.1002/glia.22770

69. Bosschaerts T, Guilliams M, Noel W, et al. Alternatively Activated Myeloid Cells Limit Pathogenicity Associated with African Trypanosomiasis through the IL-10 Inducible Gene Selenoprotein P1. The Journal of Immunology. 2008;180(9):6168–6175. doi:10.4049/jimmunol.180.9.6168

70. Short SP, Pilat JM, Williams CS. Roles for selenium and selenoprotein P in the development, progression, and prevention of intestinal disease. Free Radical Biology and Medicine. 2018/11/01/ 2018;127:26–35. 10.1016/j.freeradbiomed.2018.05.066

71. Zhang Y, Du W, Chen Z, Xiang C. Upregulation of PD-L1 by SPP1 mediates macrophage polarization and facilitates immune escape in lung adenocarcinoma. Experimental Cell Research. 2017/10/15/ 2017;359(2):449–457. 10.1016/j.yexcr.2017.08.028

72. Matsubara E, Komohara Y, Esumi S, et al. SPP1 Derived from Macrophages Is Associated with a Worse Clinical Course and Chemo-Resistance in Lung Adenocarcinoma. Cancers (Basel). Sep 8 2022;14(18)doi:10.3390/cancers14184374

73. Dong B, Wu C, Huang L, Qi Y. Macrophage-Related SPP1 as a Potential Biomarker for Early Lymph Node Metastasis in Lung Adenocarcinoma. Front Cell Dev Biol. 2021;9:739358. doi:10.3389/fcell.2021.739358

74. Park MD, Reyes-Torres I, LeBerichel J, et al. TREM2 macrophages drive NK cell paucity and dysfunction in lung cancer. Nat Immunol. May 2023;24(5):792–801. doi:10.1038/s41590-023-01475-4

75. Wculek SK, Cueto FJ, Mujal AM, Melero I, Krummel MF, Sancho D. Dendritic cells in cancer immunology and immunotherapy. Nat Rev Immunol. Jan 2020;20(1):7–24. doi:10.1038/s41577-019-0210-z

76. Li K, Fazekasova H, Wang N, et al. Expression of complement components, receptors and regulators by human dendritic cells. Mol Immunol. May 2011;48(9-10):1121–7. doi:10.1016/j.molimm.2011.02.003

77. Liu S, Wu J, Zhang T, et al. Complement C1q chemoattracts human dendritic cells and enhances migration of mature dendritic cells to CCL19 via activation of AKT and MAPK pathways. Mol Immunol. Dec 2008;46(2):242–9. doi:10.1016/j.molimm.2008.08.279

78. Li D, Yu H, Hu J, et al. Comparative profiling of single-cell transcriptome reveals heterogeneity of tumor microenvironment between solid and acinar lung adenocarcinoma. J Transl Med. Sep 23 2022;20(1):423. doi:10.1186/s12967-022-03620-3

79. Roberts EW, Broz ML, Binnewies M, et al. Critical Role for CD103+/CD141+ Dendritic Cells Bearing CCR7 for Tumor Antigen Trafficking and Priming of T Cell Immunity in Melanoma. Cancer cell. 2016/08/08/ 2016;30(2):324–336. 10.1016/j.ccell.2016.06.003

80. Yan Y, Chen R, Wang X, et al. CCL19 and CCR7 Expression, Signaling Pathways, and Adjuvant Functions in Viral Infection and Prevention. Review. Frontiers in Cell and Developmental Biology. 2019-October-01 2019;7doi:10.3389/fcell.2019.00212

81. Gobert M, Treilleux I, Bendriss-Vermare N, et al. Regulatory T Cells Recruited through CCL22/CCR4 Are Selectively Activated in Lymphoid Infiltrates Surrounding Primary Breast Tumors and Lead to an Adverse Clinical Outcome. Cancer Research. 2009;69(5):2000–2009. doi:10.1158/0008-5472.Can-08-2360

82. Krysan K, Tran LM, Grimes BS, et al. The Immune Contexture Associates with the Genomic Landscape in Lung Adenomatous Premalignancy. Cancer Research. 2019;79(19):5022–5033. doi:10.1158/0008-5472.Can-19-0153

83. Schupp JC, Adams TS, Cosme C, et al. Integrated Single-Cell Atlas of Endothelial Cells of the Human Lung. Circulation. 2021;144(4):286–302. doi:doi:10.1161/CIRCULATIONAHA.120.052318

84. Goveia J, Rohlenova K, Taverna F, et al. An Integrated Gene Expression Landscape Profiling Approach to Identify Lung Tumor Endothelial Cell Heterogeneity and Angiogenic Candidates. Cancer Cell. Jan 13 2020;37(1):21–36.e13. doi:10.1016/j.ccell.2019.12.001

85. Piper PJ, Vane JR, Wyllie JH. Inactivation of prostaglandins by the lungs. Nature. Feb 14 1970;225(5233):600–4. doi:10.1038/225600a0

86. Lambrechts D, Wauters E, Boeckx B, et al. Phenotype molding of stromal cells in the lung tumor microenvironment. Nature Medicine. 2018/08/01 2018;24(8):1277–1289. doi:10.1038/s41591-018-0096-5

87. Jeong H-W, Hernández-Rodríguez B, Kim J, et al. Transcriptional regulation of endothelial cell behavior during sprouting angiogenesis. Nature Communications. 2017/09/28 2017;8(1):726. doi:10.1038/s41467-017-00738-7

88. Lin YJ, Shyu WC, Chang CW, et al. Tumor Hypoxia Regulates Forkhead Box C1 to Promote Lung Cancer Progression. Theranostics. 2017;7(5):1177–1191. doi:10.7150/thno.17895

89. Grout JA, Sirven P, Leader AM, et al. Spatial Positioning and Matrix Programs of Cancer-Associated Fibroblasts Promote T-cell Exclusion in Human Lung Tumors. Cancer Discov. Nov 2 2022;12(11):2606–2625. doi:10.1158/2159-8290.Cd-21-1714

90. She Z, Chen H, Lin X, Li C, Su J. POSTN Regulates Fibroblast Proliferation and Migration in Laryngotracheal Stenosis Through the TGF-β/RHOA Pathway. Laryngoscope. Sep 2024;134(9):4078–4087. doi:10.1002/lary.31505

91. Hu Y, Recouvreux MS, Haro M, et al. INHBA(+) cancer-associated fibroblasts generate an immunosuppressive tumor microenvironment in ovarian cancer. npj Precision Oncology. 2024/02/15 2024;8(1):35. doi:10.1038/s41698-024-00523-y

92. Sampson N, Zenzmaier C, Heitz M, et al. Stromal Insulin-Like Growth Factor Binding Protein 3 (IGFBP3) Is Elevated in the Diseased Human Prostate and Promotes ex Vivo Fibroblast-to-Myofibroblast Differentiation. Endocrinology. 2013;154(8):2586–2599. doi:10.1210/en.2012-2259

93. Shi X, Young CD, Zhou H, Wang X. Transforming Growth Factor-β Signaling in Fibrotic Diseases and Cancer-Associated Fibroblasts. Biomolecules. Dec 12 2020;10(12)doi:10.3390/biom10121666

94. Di Gregorio J, Robuffo I, Spalletta S, et al. The Epithelial-to-Mesenchymal Transition as a Possible Therapeutic Target in Fibrotic Disorders. Review. Frontiers in Cell and Developmental Biology. 2020-December-21 2020;8doi:10.3389/fcell.2020.607483

95. Cavagnero KJ, Gallo RL. Essential immune functions of fibroblasts in innate host defense. Review. Frontiers in Immunology. 2022-December-15 2022;13doi:10.3389/fimmu.2022.1058862

96. Altorki NK, Borczuk AC, Harrison S, et al. Global evolution of the tumor microenvironment associated with progression from preinvasive invasive to invasive human lung adenocarcinoma. Cell Reports. 2022/04/05/ 2022;39(1):110639. 10.1016/j.celrep.2022.110639

97. de Visser KE, Joyce JA. The evolving tumor microenvironment: From cancer initiation to metastatic outgrowth. Cancer Cell. 2023/03/13/ 2023;41(3):374–403. 10.1016/j.ccell.2023.02.016

98. Browaeys R, Gilis J, Sang-Aram C, et al. MultiNicheNet: a flexible framework for differential cell-cell communication analysis from multi-sample multi-condition single-cell transcriptomics data. bioRxiv. 2023:2023.06.13.544751. doi:10.1101/2023.06.13.544751

99. Pérez-Gutiérrez L, Ferrara N. Biology and therapeutic targeting of vascular endothelial growth factor A. Nature Reviews Molecular Cell Biology. 2023/11/01 2023;24(11):816–834. doi:10.1038/s41580-023-00631-w

100. Hayward S, Gachehiladze M, Badr N, et al. The CD151-midkine pathway regulates the immune microenvironment in inflammatory breast cancer. The Journal of Pathology. 2020;251(1):63–73. 10.1002/path.5415

101. Chia T-Y, Billingham LK, Boland L, et al. The CXCL16-CXCR6 axis in glioblastoma modulates T-cell activity in a spatiotemporal context. Original Research. Frontiers in Immunology. 2024-January-17 2024;14doi:10.3389/fimmu.2023.1331287

102. Song GJ, Kim S-M, Park K-H, Kim J, Choi I, Cho K-H. SR-BI mediates high density lipoprotein (HDL)- induced anti-inflammatory effect in macrophages. Biochemical and Biophysical Research Communications. 2015/01/30/ 2015;457(1):112–118. 10.1016/j.bbrc.2014.12.028

103. Khantakova D, Brioschi S, Molgora M. Exploring the Impact of TREM2 in Tumor-Associated Macrophages. Vaccines. 2022;10(6):943.

104. Schraufstatter IU, Zhao M, Khaldoyanidi SK, Discipio RG. The chemokine CCL18 causes maturation of cultured monocytes to macrophages in the M2 spectrum. Immunology. Apr 2012;135(4):287–98. doi:10.1111/j.1365-2567.2011.03541.x

105. Krohn SC, Bonvin P, Proudfoot AEI. CCL18 Exhibits a Regulatory Role through Inhibition of Receptor and Glycosaminoglycan Binding. PLOS ONE. 2013;8(8):e72321. doi:10.1371/journal.pone.0072321

106. Benschop R, Wei T, Na S. Tumor necrosis factor receptor superfamily member 21: TNFR-related death receptor-6, DR6. Adv Exp Med Biol. 2009;647:186–94. doi:10.1007/978-0-387-89520-8_13

107. Ma J, Huang L, Gao Y-B, Li M-X, Chen L-L, Yang L. Circ_TNFRSF21 promotes cSCC metastasis and M2 macrophage polarization via miR-214-3p/CHI3L1. Journal of Dermatological Science. 2023/08/01/ 2023;111(2):32–42. 10.1016/j.jdermsci.2023.06.001

108. Xu R, Li Y, Yan H, et al. CCL2 promotes macrophages-associated chemoresistance via MCPIP1 dual catalytic activities in multiple myeloma. Cell Death & Disease. 2019/10/14 2019;10(10):781. doi:10.1038/s41419-019-2012-4

109. Fridlender ZG, Kapoor V, Buchlis G, et al. Monocyte Chemoattractant Protein–1 Blockade Inhibits Lung Cancer Tumor Growth by Altering Macrophage Phenotype and Activating CD8+ Cells. American Journal of Respiratory Cell and Molecular Biology. 2011;44(2):230–237. doi:10.1165/rcmb.2010-0080OC

110. Gocher AM, Workman CJ, Vignali DAA. Interferon-γ: teammate or opponent in the tumour microenvironment? Nat Rev Immunol. Mar 2022;22(3):158–172. doi:10.1038/s41577-021-00566-3

111. Petroni G, Pillozzi S, Antonuzzo L. Exploiting Tertiary Lymphoid Structures to Stimulate Antitumor Immunity and Improve Immunotherapy Efficacy. Cancer Res. Apr 15 2024;84(8):1199–1209. doi:10.1158/0008-5472.CAN-23-3325

112. Workel HH, Lubbers JM, Arnold R, et al. A Transcriptionally Distinct CXCL13. Cancer Immunol Res. May 2019;7(5):784–796. doi:10.1158/2326-6066.CIR-18-0517

113. Asrir A, Tardiveau C, Coudert J, et al. Tumor-associated high endothelial venules mediate lymphocyte entry into tumors and predict response to PD-1 plus CTLA-4 combination immunotherapy. Cancer Cell. Mar 14 2022;40(3):318–334.e9. doi:10.1016/j.ccell.2022.01.002

114. Lee HS, Jang HJ, Ramineni M, et al. A Phase II Window of Opportunity Study of Neoadjuvant PD-L1 versus PD-L1 plus CTLA-4 Blockade for Patients with Malignant Pleural Mesothelioma. Clin Cancer Res. Feb 1 2023;29(3):548–559. doi:10.1158/1078-0432.Ccr-22-2566

115. Han G, Sinjab A, Rahal Z, et al. An atlas of epithelial cell states and plasticity in lung adenocarcinoma. Nature. Mar 2024;627(8004):656–663. doi:10.1038/s41586-024-07113-9

116. Lin C, Song H, Huang C, et al. Alveolar type II cells possess the capability of initiating lung tumor development. PLoS One. 2012;7(12):e53817. doi:10.1371/journal.pone.0053817

117. Osborne JK, Minna JD. Lung Cancer Cell of Origin: Controversy and Clinical Translational Implications. Cancer Res. Mar 15 2022;82(6):972–973. doi:10.1158/0008-5472.CAN-22-0301

118. Rowbotham SP, Kim CF. Diverse cells at the origin of lung adenocarcinoma. Proc Natl Acad Sci U S A Apr 01 2014;111(13):4745–6. doi:10.1073/pnas.1401955111

119. Brea EJ, Oh CY, Manchado E, et al. Kinase Regulation of Human MHC Class I Molecule Expression on Cancer Cells. Cancer Immunol Res. Nov 2016;4(11):936–947. doi:10.1158/2326-6066.CIR-16-0177

120. Zhang C, Zhang J, Xu FP, et al. Genomic Landscape and Immune Microenvironment Features of Preinvasive and Early Invasive Lung Adenocarcinoma. J Thorac Oncol. Nov 2019;14(11):1912–1923. doi:10.1016/j.jtho.2019.07.031

121. Shimasaki N, Jain A, Campana D. NK cells for cancer immunotherapy. Nat Rev Drug Discov. Mar 2020;19(3):200–218. doi:10.1038/s41573-019-0052-1

122. Maier B, Leader AM, Chen ST, et al. A conserved dendritic-cell regulatory program limits antitumour immunity. Nature. Apr 2020;580(7802):257–262. doi:10.1038/s41586-020-2134-y

123. Zhang Y, Xu M, Ren Y, et al. Tertiary lymphoid structural heterogeneity determines tumour immunity and prospects for clinical application. Mol Cancer. Apr 06 2024;23(1):75. doi:10.1186/s12943-024-01980-6

124. Im SJ, Obeng RC, Nasti TH, et al. Characteristics and anatomic location of PD-1. Proc Natl Acad Sci U S A. Oct 10 2023;120(41):e2221985120. doi:10.1073/pnas.2221985120

125. Ly D, Li Q, Navab R, et al. Tumor-Associated Regulatory T Cell Expression of LAIR2 Is Prognostic in Lung Adenocarcinoma. Cancers. 2022;14(1):205.

126. Heuvelmans MA, Walter JE, Peters RB, et al. Relationship between nodule count and lung cancer probability in baseline CT lung cancer screening: The NELSON study. Lung Cancer. Nov 2017;113:45–50. doi:10.1016/j.lungcan.2017.08.023

127. van Montfoort N, Borst L, Korrer MJ, et al. NKG2A Blockade Potentiates CD8 T Cell Immunity Induced by Cancer Vaccines. Cell. Dec 13 2018;175(7):1744–1755.e15. doi:10.1016/j.cell.2018.10.028

128. André P, Denis C, Soulas C, et al. Anti-NKG2A mAb Is a Checkpoint Inhibitor that Promotes Anti-tumor Immunity by Unleashing Both T and NK Cells. Cell. Dec 13 2018;175(7):1731–1743.e13. doi:10.1016/j.cell.2018.10.014

129. Ferrari de Andrade L, Tay RE, Pan D, et al. Antibody-mediated inhibition of MICA and MICB shedding promotes NK cell-driven tumor immunity. Science. Mar 30 2018;359(6383):1537–1542. doi:10.1126/science.aao0505

130. Freeman ZT, Nirschl TR, Hovelson DH, et al. A conserved intratumoral regulatory T cell signature identifies 4-1BB as a pan-cancer target. J Clin Invest. Mar 02 2020;130(3):1405–1416. doi:10.1172/JCI128672

131. Buchan SL, Dou L, Remer M, et al. Antibodies to Costimulatory Receptor 4-1BB Enhance Anti-tumor Immunity via T Regulatory Cell Depletion and Promotion of CD8 T Cell Effector Function. Immunity. Nov 20 2018;49(5):958–970.e7. doi:10.1016/j.immuni.2018.09.014

132. Kidani Y, Nogami W, Yasumizu Y, et al. CCR8-targeted specific depletion of clonally expanded Treg cells in tumor tissues evokes potent tumor immunity with long-lasting memory. Proc Natl Acad Sci U S A. Feb 15 2022;119(7)doi:10.1073/pnas.2114282119

133. Campbell JR, McDonald BR, Mesko PB, et al. Fc-Optimized Anti-CCR8 Antibody Depletes Regulatory T Cells in Human Tumor Models. Cancer Res. Jun 01 2021;81(11):2983–2994. doi:10.1158/0008-5472.CAN-20-3585

134. Molgora M, Esaulova E, Vermi W, et al. TREM2 Modulation Remodels the Tumor Myeloid Landscape Enhancing Anti-PD-1 Immunotherapy. Cell. Aug 20 2020;182(4):886–900.e17. doi:10.1016/j.cell.2020.07.013

135. Wang YH, Cheng TY, Chen TY, Chang KM, Chuang VP, Kao KJ. Plasmalemmal Vesicle Associated Protein (PLVAP) as a therapeutic target for treatment of hepatocellular carcinoma. BMC Cancer. Nov 6 2014;14:815. doi:10.1186/1471-2407-14-815

136. Zheng GXY, Terry JM, Belgrader P, et al. Massively parallel digital transcriptional profiling of single cells. Nature Communications. 2017/01/16 2017;8(1):14049. doi:10.1038/ncomms14049

137. Yang S, Corbett SE, Koga Y, et al. Decontamination of ambient RNA in single-cell RNA-seq with DecontX. Genome Biology. 2020/03/05 2020;21(1):57. doi:10.1186/s13059-020-1950-6

138. Yuhan H, Stephanie H, Erica A-N, et al. Integrated analysis of multimodal single-cell data. Cell. 2021;184(13):3573–3587.e29. 10.1016/j.cell.2021.04.048

139. McGinnis CS, Murrow LM, Gartner ZJ. DoubletFinder: Doublet Detection in Single-Cell RNA Sequencing Data Using Artificial Nearest Neighbors. Cell Systems. 2019;8(4):329–337.e4. doi:10.1016/j.cels.2019.03.003

140. Korsunsky I, Millard N, Fan J, et al. Fast, sensitive and accurate integration of single-cell data with Harmony. Nature Methods. 2019;16(12):1289–1296. doi:10.1038/s41592-019-0619-0

141. inferCNV of the Trinity CTAT Project. https://github.com/broadinstitute/inferCNV

142. Tirosh I, Venteicher AS, Hebert C, et al. Single-cell RNA-seq supports a developmental hierarchy in human oligodendroglioma. Nature. 2016/11/01 2016;539(7628):309–313. doi:10.1038/nature20123

143. Aibar S, González-Blas CB, Moerman T, et al. SCENIC: single-cell regulatory network inference and clustering. Nat Methods. Nov 2017;14(11):1083–1086. doi:10.1038/nmeth.4463

144. Browaeys R, Gilis J, Sang-Aram C, et al. MultiNicheNet: a flexible framework for differential cell-cell communication analysis from multi-sample multi-condition single-cell transcriptomics data. bioRxiv. 2023:2023.06.13.544751. doi:10.1101/2023.06.13.544751

145. Wu Y, Yang S, Ma J, et al. Spatiotemporal Immune Landscape of Colorectal Cancer Liver Metastasis at Single-Cell Level. Cancer Discov. Jan 2022;12(1):134–153. doi:10.1158/2159-8290.Cd-21-0316

146. Kanehisa M, Goto S. KEGG: kyoto encyclopedia of genes and genomes. Nucleic Acids Res. Jan 1 2000;28(1):27–30. doi:10.1093/nar/28.1.27

147. Fabregat A, Sidiropoulos K, Viteri G, et al. Reactome pathway analysis: a high-performance in-memory approach. BMC Bioinformatics. Mar 2 2017;18(1):142. doi:10.1186/s12859-017-1559-2

148. Dobin A, Davis CA, Schlesinger F, et al. STAR: ultrafast universal RNA-seq aligner. Bioinformatics. Jan 01 2013;29(1):15–21. doi:10.1093/bioinformatics/bts635

149. Frankish A, Diekhans M, Ferreira AM, et al. GENCODE reference annotation for the human and mouse genomes. Nucleic Acids Res. Jan 08 2019;47(D1):D766–D773. doi:10.1093/nar/gky955

150. Trapnell C, Williams BA, Pertea G, et al. Transcript assembly and quantification by RNA-Seq reveals unannotated transcripts and isoform switching during cell differentiation. Nat Biotechnol. May 2010;28(5):511–5. doi:10.1038/nbt.1621

151. Aran D, Hu Z, Butte AJ. xCell: digitally portraying the tissue cellular heterogeneity landscape. Genome Biol. 11 2017;18(1):220. doi:10.1186/s13059-017-1349-1

152. Barbie DA, Tamayo P, Boehm JS, et al. Systematic RNA interference reveals that oncogenic KRAS- driven cancers require TBK1. Nature. Nov 05 2009;462(7269):108–12. doi:10.1038/nature08460

153. Berg S, Kutra D, Kroeger T, et al. ilastik: interactive machine learning for (bio)image analysis. Nat Methods. Dec 2019;16(12):1226–1232. doi:10.1038/s41592-019-0582-9

154. Greenwald NF, Miller G, Moen E, et al. Whole-cell segmentation of tissue images with human-level performance using large-scale data annotation and deep learning. Nat Biotechnol. Apr 2022;40(4):555–565. doi:10.1038/s41587-021-01094-0

155. Espinosa-Carrasco G, Chiu E, Scrivo A, et al. Intratumoral immune triads are required for immunotherapy-mediated elimination of solid tumors. Cancer Cell. Jul 8 2024;42(7):1202–1216.e8. doi:10.1016/j.ccell.2024.05.025

156. Zhang H, Hunter MV, Chou J, et al. BayesTME: An end-to-end method for multiscale spatial transcriptional profiling of the tissue microenvironment. Cell Syst. Jul 19 2023;14(7):605–619.e7. doi:10.1016/j.cels.2023.06.003

