## Supplemental figures for "Acquisition of discrete immune suppressive barriers contributes to the initiation and progression of preinvasive to invasive human lung cancer"

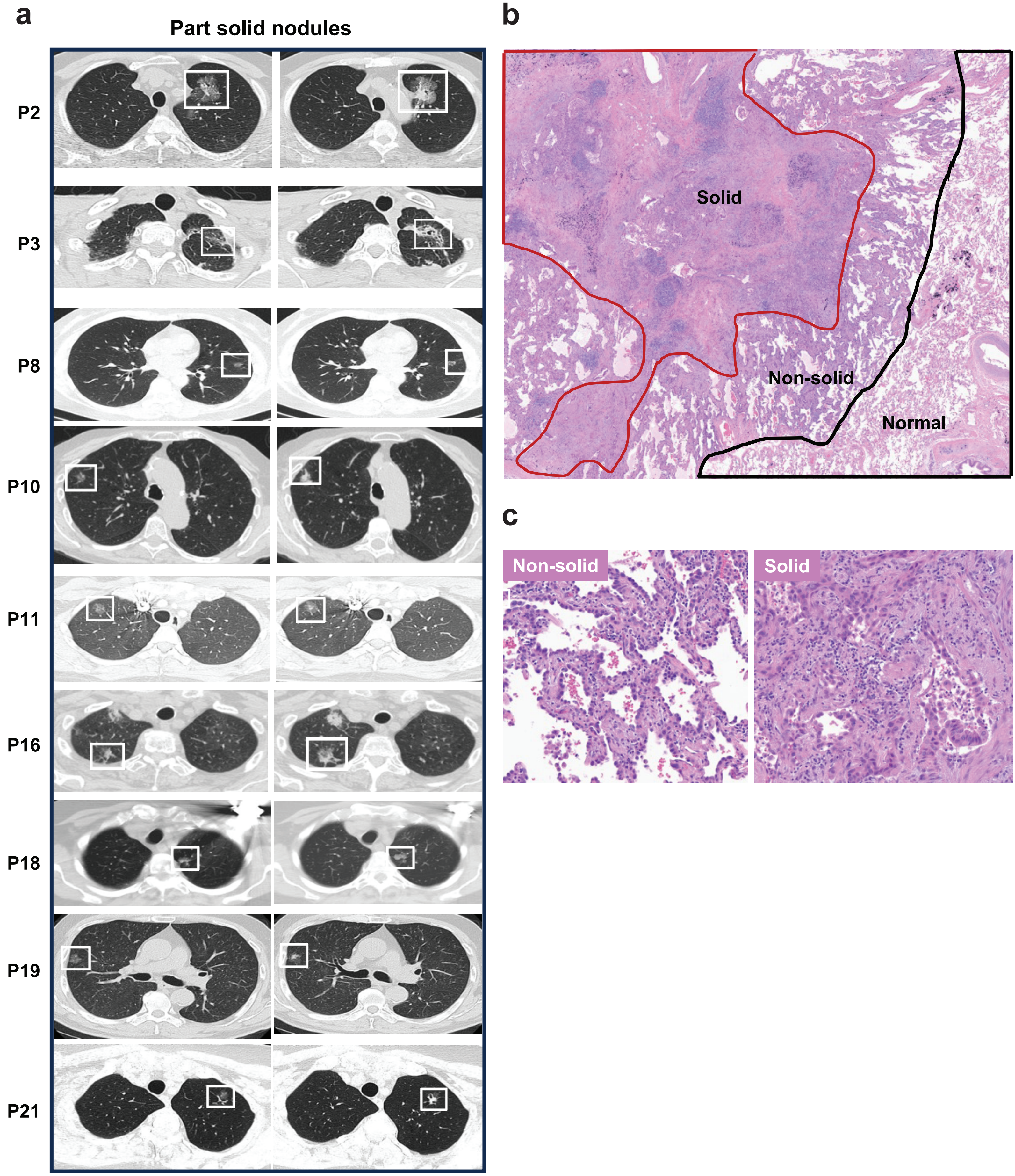


**Supplementary Figure 1.1. Representative CT scan and IHC images of patients with part-solid nodules.**

a. CT scans showing lung nodules (boxed) in individual patients.
b. Representative H&E image of a part-solid nodule showing adjacent normal lung, non-solid and invasive solid regions.
c. H&E images of non-solid (lepidic histology, left panel) and solid (right panel) regions.

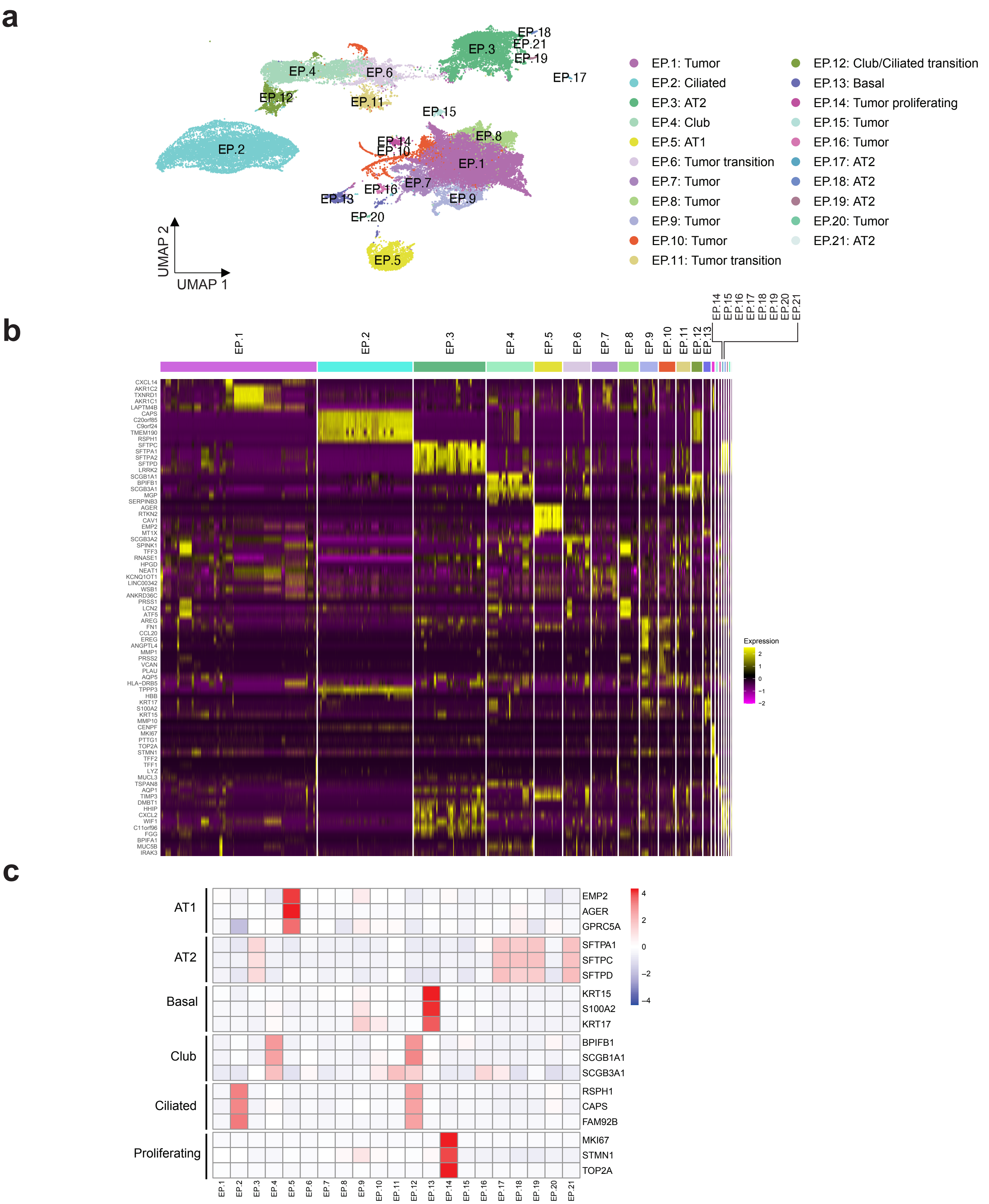


**Supplementary Figure 2.1. Characterization of the epithelial cell clusters.**

a. UMAP visualization of reclustered epithelial cells, illustrating 21 distinct clusters.
b. Heatmap showing the relative expression of the top five cluster markers with the highest fold-change values for each epithelial cell cluster.
c. Heatmaps showing mean expression levels of established markers for epithelial cell types within the epithelial cell clusters.

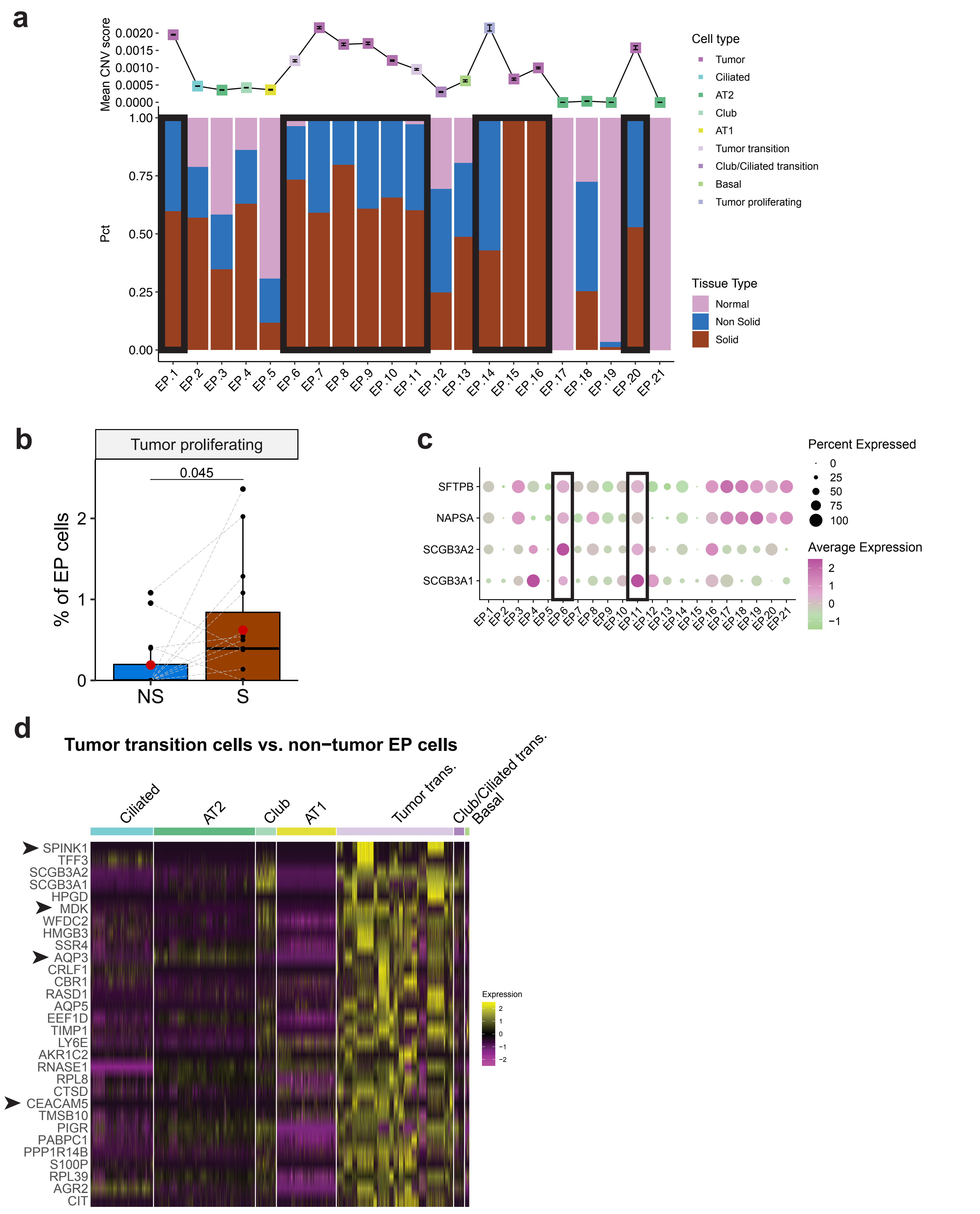
a
Mean CNV score
0.0020
0.0015
0.0010
0.0005
0.0000
Pct
1.00
0.75
0.50
0.25
0.00
EP.1
EP.2
EP.3
EP.4
EP.5
EP.6
EP.7
EP.8
EP.9
EP.10
EP.11
EP.12
EP.13
EP.14
EP.15
EP.16
EP.17
EP.18
EP.19
EP.20
EP.21
Cell type
Tumor
Ciliated
AT2
Club
AT1
Tumor transition
Club/Ciliated transition
Basal
Tumor proliferating
Tissue Type
Normal
Non Solid
Solid
b
Tumor proliferating
0.045
% of EP cells
2
1
0
NS
S
c
SFTPB
NAPSA
SCGB3A2
SCGB3A1
EP1
EP2
EP3
EP4
EP5
EP6
EP7
EP8
EP9
EP10
EP11
EP12
EP13
EP14
EP15
EP16
EP17
EP18
EP19
EP20
EP21
Percent Expressed
0
25
50
75
100
Average Expression
2
1
0
−1
d
Tumor transition cells vs. non−tumor EP cells
Ciliated
AT2
Club
AT1
Tumor trans.
Club/Ciliated trans.
Basal
SPINK1
TFF3
SCGB3A2
SCGB3A1
HPGD
MDK
WFDC2
HMGB3
SSR4
AQP3
CRLF1
CBR1
RASD1
AQP5
EEF1D
TIMP1
LY6E
AKR1C2
RNASE1
RPL8
CTSD
CEACAM5
TMSB10
PIGR
PABPC1
PPP1R14B
S100P
RPL39
AGR2
CIT
Expression
2
1
0
−1
−2

**Supplementary Figure 2.2. Epithelial cells transition to tumor cells.**

a. CNV scores (upper panel) and the percentage of epithelial cells across normal lung, non-solid, and solid components (lower panel) within each epithelial cells subtype. High-CNV tumor cell clusters (including 'tumor', 'tumor-transition', and 'tumor proliferating') are indicated by black rectangles. Pct, percentage
b. Proportions of tumor proliferating cells in non-solid (NS) and solid (S) components. Percentages are based on the entire epithelial cell population. P-values were obtained using the Wilcoxon matched-pairs signed rank test on paired samples from the same patients, with matched samples connected by a line. Mean values are indicated by red dots.
c. Dot plot showing the expression levels of AT2 and club cell markers in epithelial cell populations. Expression level is depicted by color, and the size of the dots represents the proportion of cells expressing these markers.
d. Heatmap displaying the relative expression of the top 30 genes with the highest fold-change values that are significantly upregulated (adjusted p-value $\leq 0.05$) in tumor transition cells compared to non-tumor epithelial cells. Protumorigenic genes highlighted in the text are depicted with arrowheads.

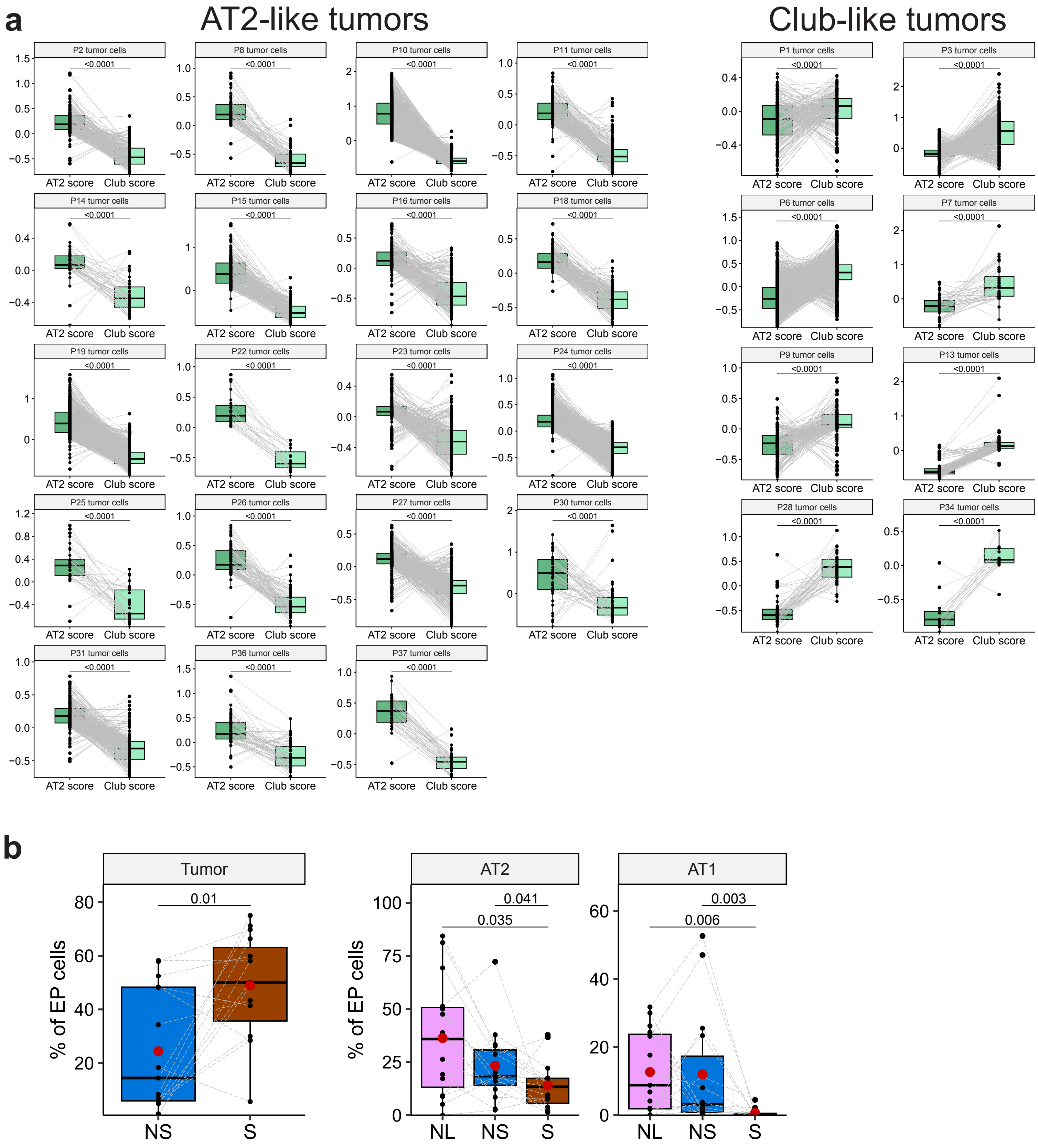


**Supplementary Figure 2.3. Characterization of AT2-like and club-like tumors in individual patients.**

a. A comparison between the AT2 scores and club scores of tumor cells across all patients. Each dot represents a score of one cell and dots are connected by a grey line if those scores are of the same cell. P-values were obtained using the Wilcoxon matched-pairs signed rank test on matched scores. Patients with AT2-like tumors are grouped on the left, and patients with club-like tumors are grouped on the right.

b. Proportions of Tumor, AT2 and AT1 cells in adjacent normal tissue (NL), non-solid (NS) and solid (S) components. Percentages are based on the entire epithelial cell population. P-values were obtained using the Wilcoxon matched-pairs signed rank test on paired samples from the same patients, with matched samples connected by a line. Mean values are indicated by red dots. Only significant p-values (< 0.05) are shown.

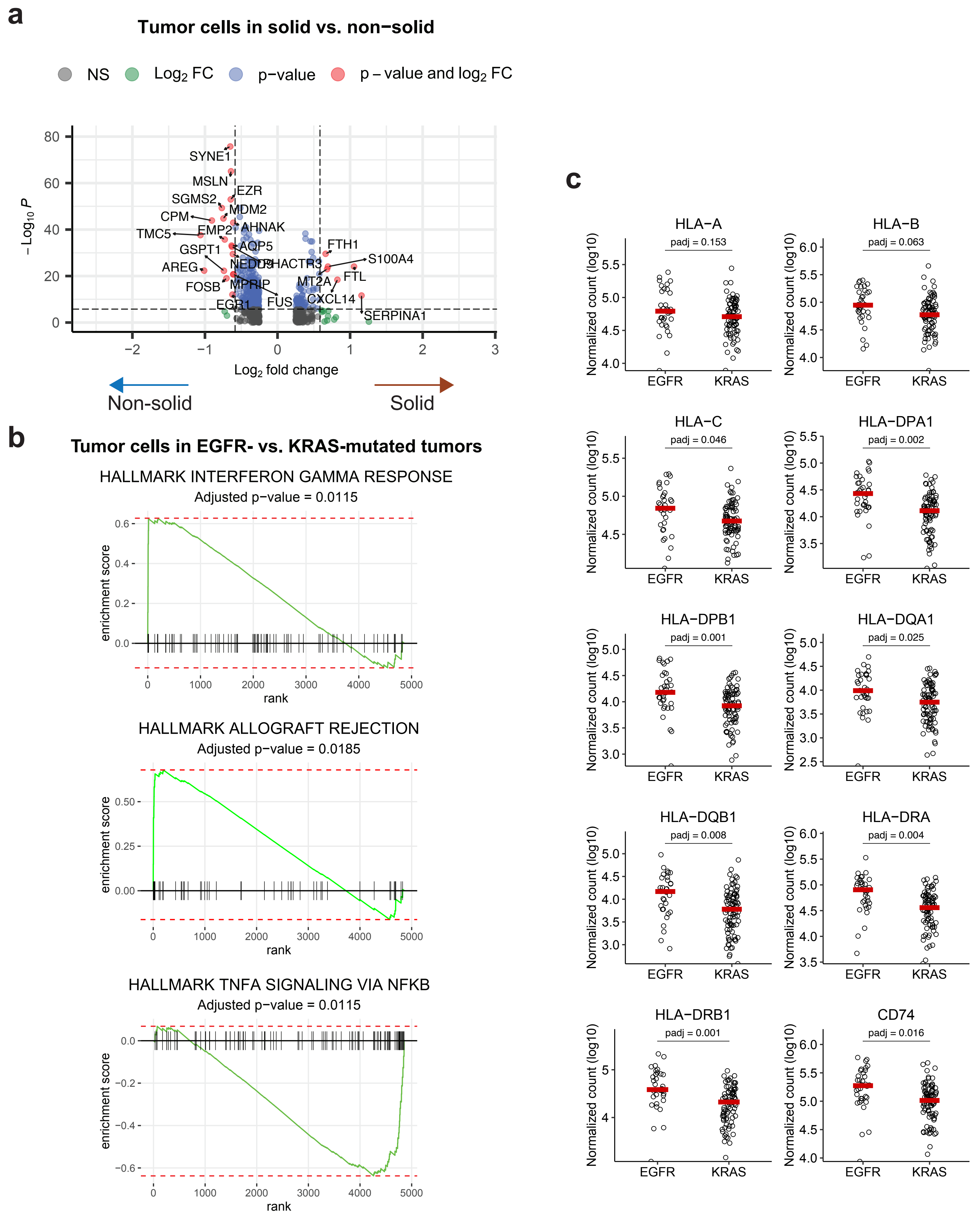

a
Tumor cells in solid vs. non−solid
NS
Log2 FC
p−value
p – value and log2 FC
SYNE1
MSLN
EZR
SGMS2
MDM2
CPM
AHNAK
TMC5
EMP2
AQP5
GSPT1
FTH1
S100A4
AREG
FTL
FOSB
MPRIP
MT2A
FUS
CXCL14
EGR1
SERPINA1
Log2 fold change
– Log10 P
Non-solid
Solid
b
Tumor cells in EGFR- vs. KRAS-mutated tumors
HALLMARK INTERFERON GAMMA RESPONSE
Adjusted p−value = 0.0115
HALLMARK ALLOGRAFT REJECTION
Adjusted p−value = 0.0185
HALLMARK TNFA SIGNALING VIA NFKB
Adjusted p−value = 0.0115
enrichment score
rank
c
HLA−A
padj = 0.153
HLA−B
padj = 0.063
HLA−C
padj = 0.046
HLA−DPA1
padj = 0.002
HLA−DPB1
padj = 0.001
HLA−DQA1
padj = 0.025
HLA−DQB1
padj = 0.008
HLA−DRA
padj = 0.004
HLA−DRB1
padj = 0.001
CD74
padj = 0.016
Normalized count (log10)
EGFR
KRAS

**Supplementary Figure 2.4. Transcriptomic changes in tumor cells from solid and non-solid components and from EGFR and KRAS-mutated tumors.**

a. Volcano plot comparing gene expression between tumor cells in solid and non-solid samples. The x-axis represents the log2 fold change in gene expression, with positive values indicating higher expression in solid samples and negative values indicating higher expression in non-solid samples. The y-axis represents the -log10 of the p-value, with higher values indicating greater statistical significance. Green dots indicate genes with significant fold changes (≥ 1.5), blue dots indicate genes with significant adjusted p-values (≤ 0.05), and red dots indicate genes significant in both fold change and p-value.
b. Enrichment plots of upregulated (upper and middle panels) and downregulated (lower panel) Hallmark pathways in tumor cells in EGFR-mutated tumors vs. KRAS-mutated tumors. Enrichment scores and p-values were obtained using GSEA with an FDR-adjusted p-value shown.
c. Gene expression of MHC class I and MHC class II genes in EGFR-mutated vs KRAS-mutated tumors from the RNA-seq dataset of the TCGA-LUAD cohort (tumor purity > 70%). P-values were derived from a differential expression test on all expressed genes using the DESeq2 R package, with adjusted p-values shown after FDR multiple test correction.

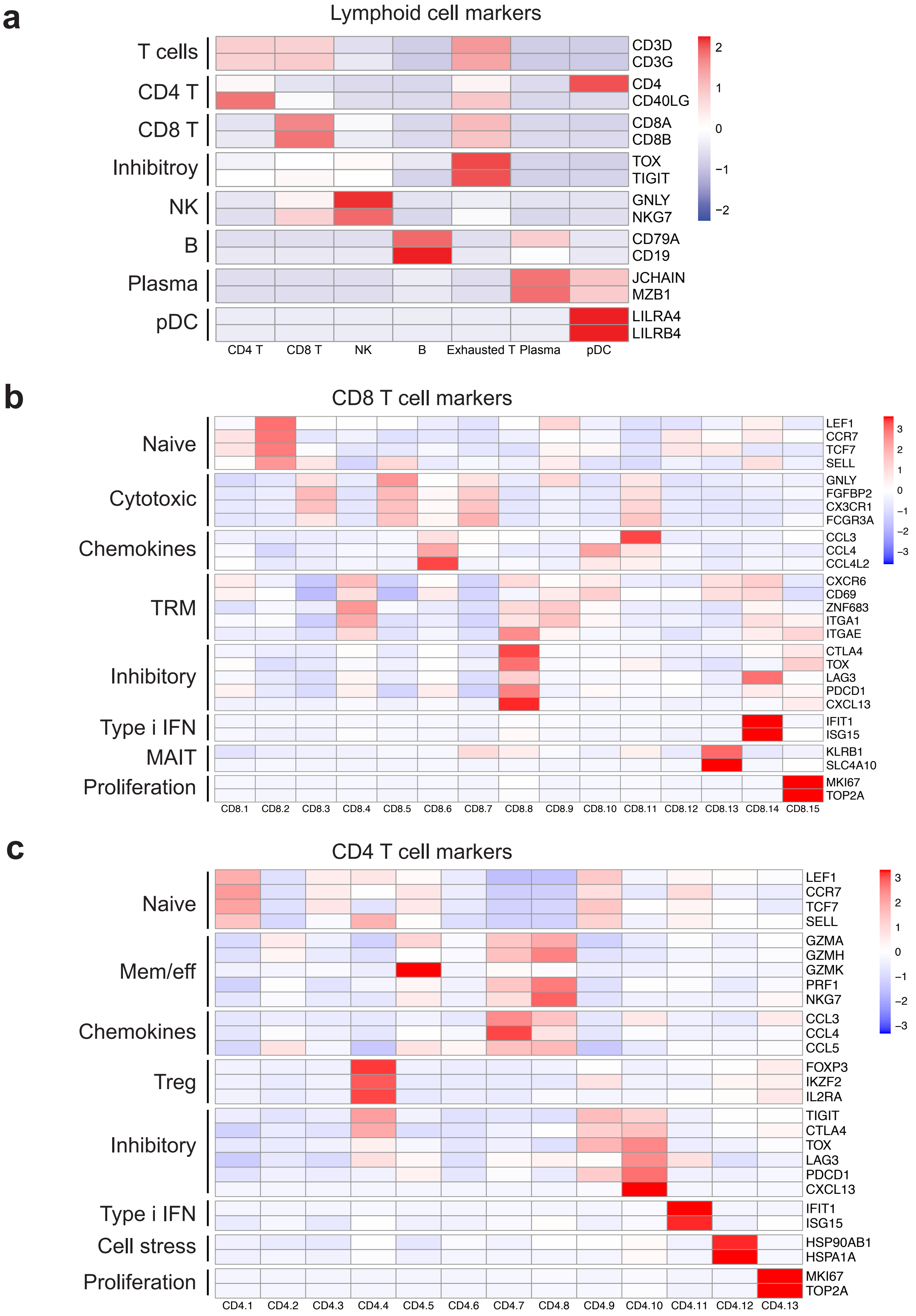


**Supplementary Figure 3.1. Expression of canonical markers in lymphoid cell populations.**

Heatmaps showing mean expression of canonical markers for lymphoid cell types within clusters of lymphoid cells (a), CD8 T cells (b) and CD4 T cells (c).

**a**

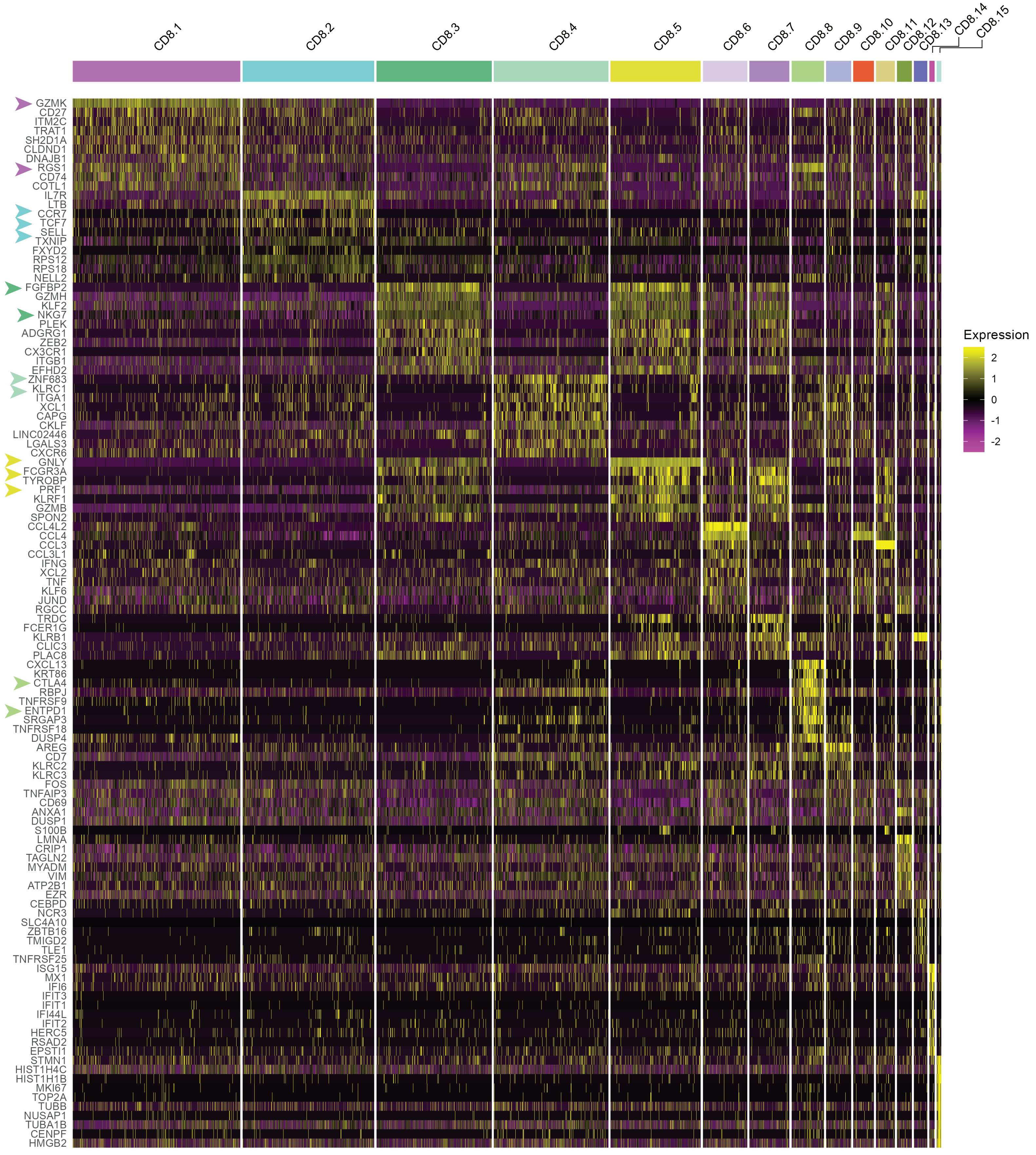


**Supplementary Figure 3.2. CD8 T cells top cluster markers expression.**

a. Heatmap showing the relative expression of the top ten cluster markers with the highest fold-change values for each CD8 T cell cluster. Key genes in each cluster are highlighted with arrowheads.

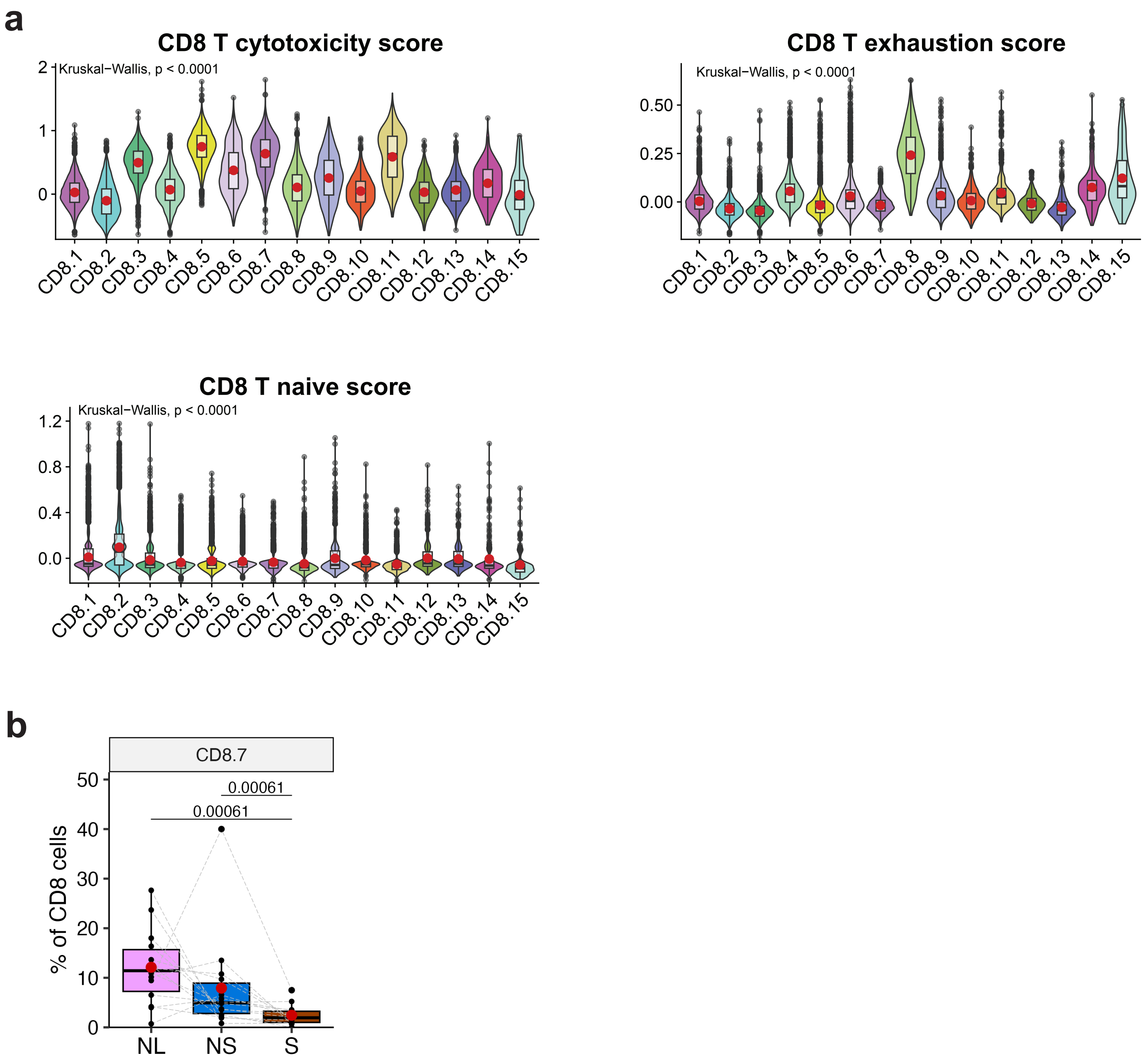


**Supplementary Figure 3.3. Phenotypic characterization of CD8 T cells.**

a. Violin and box plots representing the cytotoxicity (upper left), exhaustion (upper right), and naïve (lower panel) scores across clusters of CD8 T cells, with mean values marked by red dots. P-values were calculated using the Kruskal-Wallis test.
b. Proportion of CD8.7 in adjacent normal tissue (NL), non-solid (NS) and solid (S) lesion components. Percentages are based on the entire CD8 T cell population. P-values were obtained using the Wilcoxon matched-pairs signed rank test on paired samples from the same patients, with matched samples connected by a line. Mean values are indicated by red dots. Only significant p-values (< 0.05) are shown.

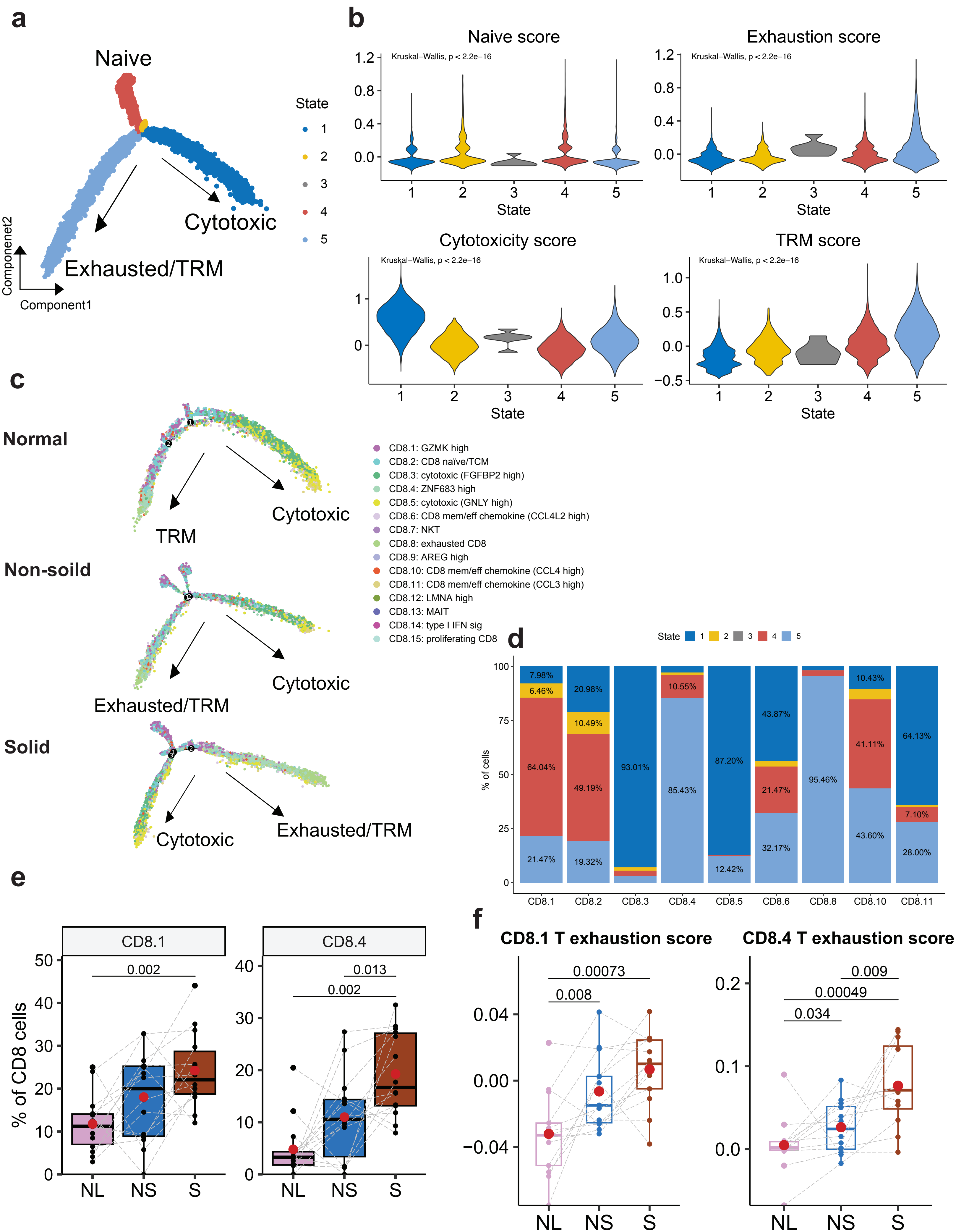

a
Naive
State
1
2
3
4
5
Cytotoxic
Exhausted/TRM
Componenet2
Component1
b
Naive score
Kruskal–Wallis, p < 2.2e−16
State
Exhaustion score
Kruskal–Wallis, p < 2.2e−16
State
Cytotoxicity score
Kruskal–Wallis, p < 2.2e−16
State
TRM score
Kruskal–Wallis, p < 2.2e−16
State
c
Normal
Cytotoxic
TRM
Non-soild
Cytotoxic
Exhausted/TRM
Solid
Cytotoxic
Exhausted/TRM
CD8.1: GZMK high
CD8.2: CD8 naïve/TCM
CD8.3: cytotoxic (FGFBP2 high)
CD8.4: ZNF683 high
CD8.5: cytotoxic (GNLY high)
CD8.6: CD8 mem/eff chemokine (CCL4L2 high)
CD8.7: NKT
CD8.8: exhausted CD8
CD8.9: AREG high
CD8.10: CD8 mem/eff chemokine (CCL4 high)
CD8.11: CD8 mem/eff chemokine (CCL3 high)
CD8.12: LMNA high
CD8.13: MAIT
CD8.14: type I IFN sig
CD8.15: proliferating CD8
d
State 1 2 3 4 5
% of cells
7.98%
6.46%
64.04%
21.47%
20.98%
10.49%
49.19%
19.32%
93.01%
10.55%
85.43%
87.20%
12.42%
43.87%
21.47%
32.17%
95.46%
10.43%
41.11%
43.60%
64.13%
7.10%
28.00%
CD8.1
CD8.2
CD8.3
CD8.4
CD8.5
CD8.6
CD8.8
CD8.10
CD8.11
e
CD8.1
CD8.4
% of CD8 cells
0.002
0.013
0.002
NL
NS
S
f
CD8.1 T exhaustion score
CD8.4 T exhaustion score
0.00073
0.008
0.009
0.00049
0.034
NL
NS
S

**Supplementary Figure 3.4. CD8 T cells trajectories to exhaustion and cytotoxicity.**

a. Trajectory plot corresponding to Fig. 3h, showing the inferred trajectory of CD8 T cells. Cells are color-coded according to the trajectory segments referred to as ‘states’ as determined by Monocle2 analysis.
b. Violin plots represent the naïve (upper left), exhaustion (upper right), cytotoxicity (lower left), and TRM (lower right) scores across CD8 T trajectory states. P-values were calculated using the Kruskal-Wallis test.
c. Trajectory plots illustrating the inferred progression of CD8 T cells from naïve to cytotoxic or exhausted/TRM states based on Monocle2 analysis of samples from adjacent normal tissue (upper panel), non-solid (middle), and solid (lower) components. Cells are color-coded by cluster.
d. Percentage of cells across the five defined trajectory states within each CD8 T cells subtype.
e. Proportion of pre-exhausted CD8 T cell populations (CD8.1 and CD8.4) in adjacent normal tissue (NL), non-solid (NS) and solid (S) lesion components. Percentages are based on the entire CD8 T cell population. P-values were obtained using the Wilcoxon matched-pairs signed rank test on paired samples from the same patients, with matched samples connected by a line. Mean values are indicated by red dots. Only significant p-values (< 0.05) are shown.

**a**

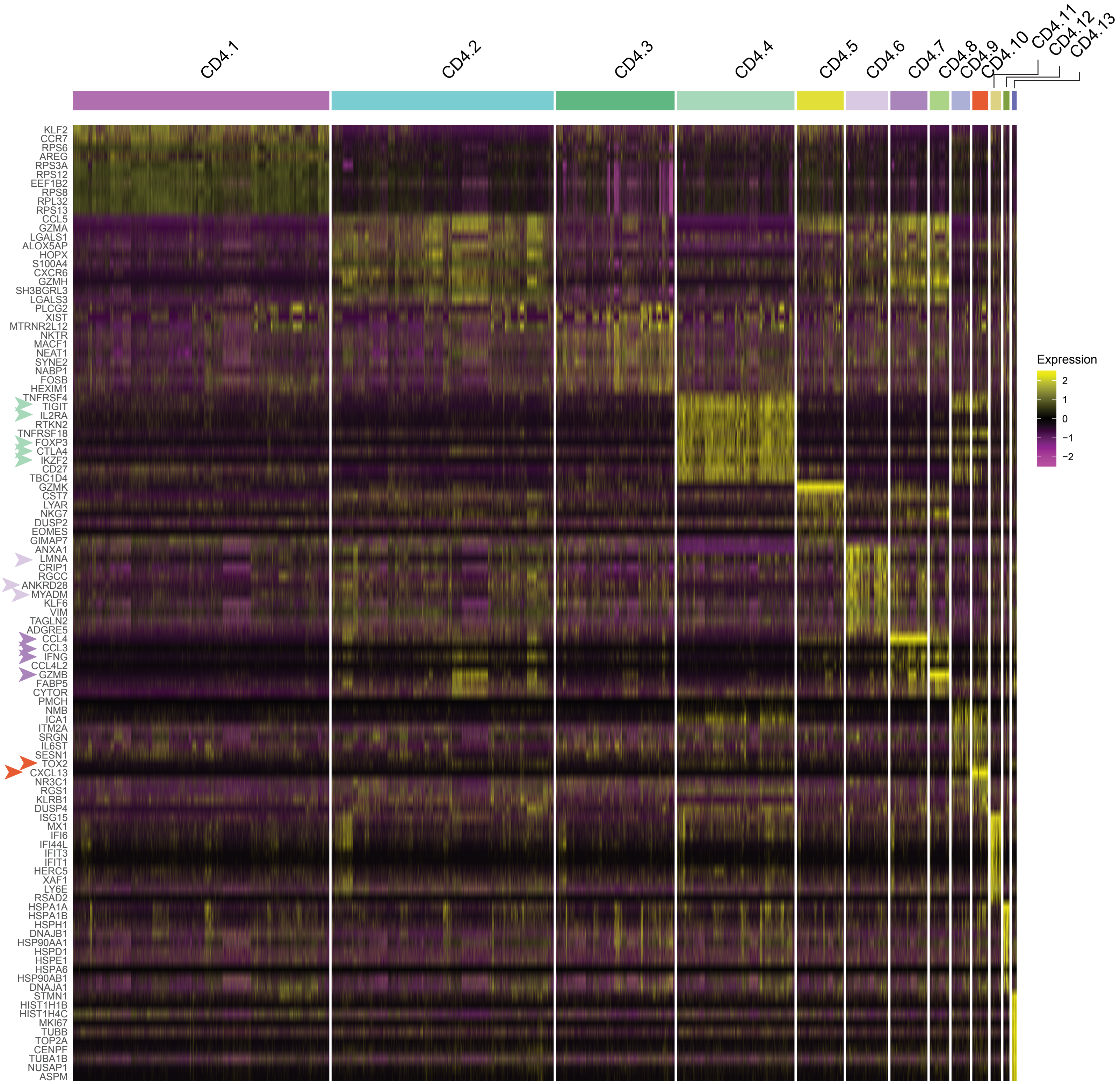


**Supplementary Figure 3.5. CD4 T cell top cluster marker expression.**

a. Heatmap showing the relative expression of the top ten cluster markers with the highest fold-change values for each CD4 T cell cluster.

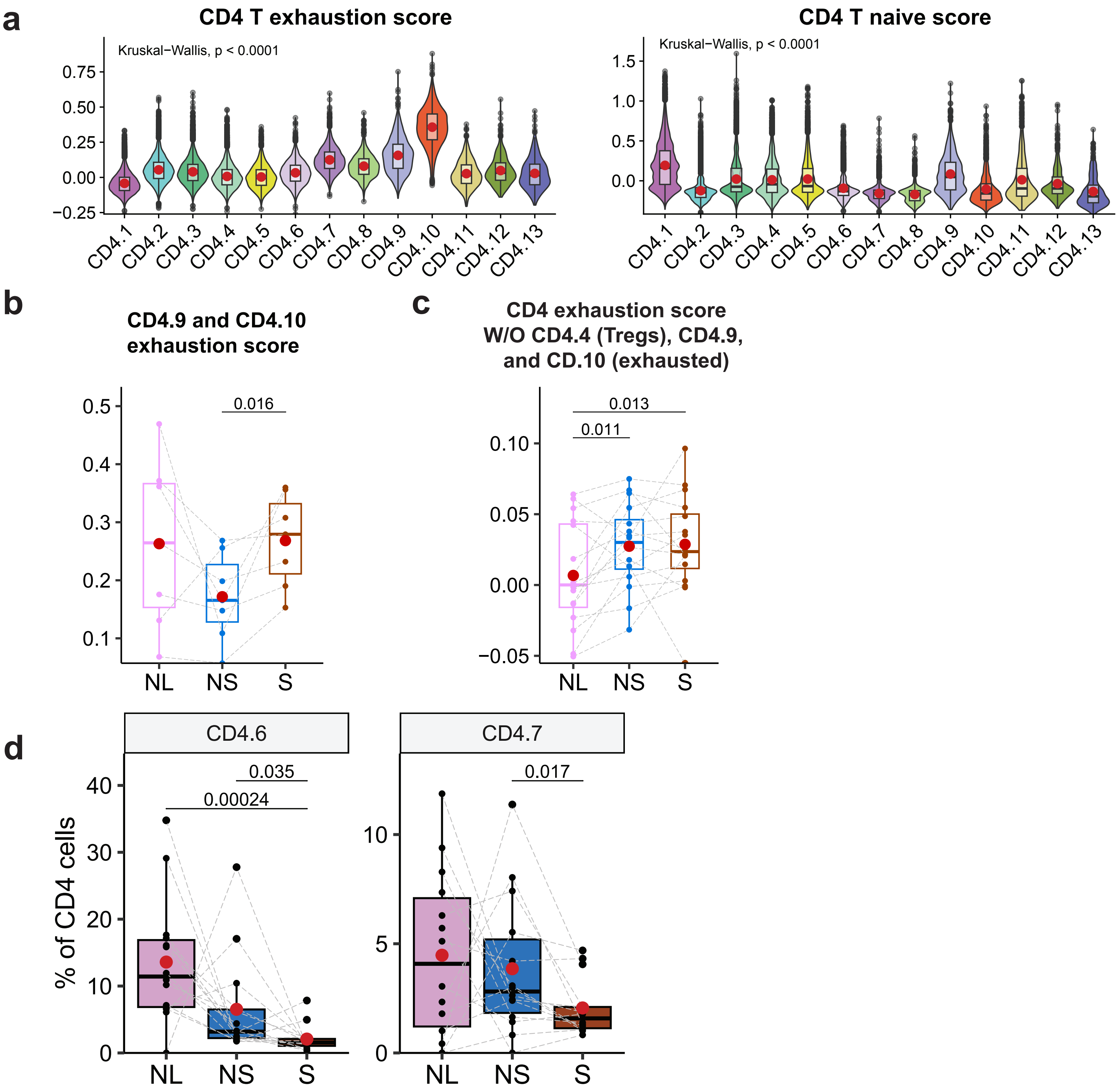


**Supplementary Figure 3.6. Phenotypic characterization of CD4 T cells.**

a. Violin and box plots represent the exhaustion (left) and naïve (right) scores across CD4 T cell clusters, with mean values marked by red dots. P-values were calculated using the Kruskal-Wallis test.
b. Exhaustion scores of CD4.9 and CD4.10 T cells in adjacent normal tissue (NL), non-solid (NS) and solid (S) components. Scores are averaged per sample. P-values were obtained using the Wilcoxon matched-pairs signed rank test on paired samples from the same patients, with matched samples connected by a line. Mean values are indicated by red dots. Only significant p-values (< 0.05) are shown.
c. Mean exhaustion scores per sample of CD4 T cells excluding Tregs (CD4.4) and exhausted cells (CD4.9 and CD4.10) across adjacent normal tissue (NL), non-solid (NS) and solid (S) components. Mean values per tissue type are marked by red dots. P-values were calculated using the Wilcoxon matched-pairs signed rank test on paired samples from the same patients with matched samples connected by a line. Only significant p-values (< 0.05) are shown.
d. Proportions of clusters CD4.6 and CD4.7 in adjacent normal tissue (NL), non-solid (NS) and solid (S) lesion components. Percentages are based on the entire CD4 T cell population. P-values were obtained using the Wilcoxon matched-pairs signed rank test on paired samples from the same patients with matched samples connected by a line. Mean values are indicated by red dots. Only significant p-values (< 0.05) are shown.

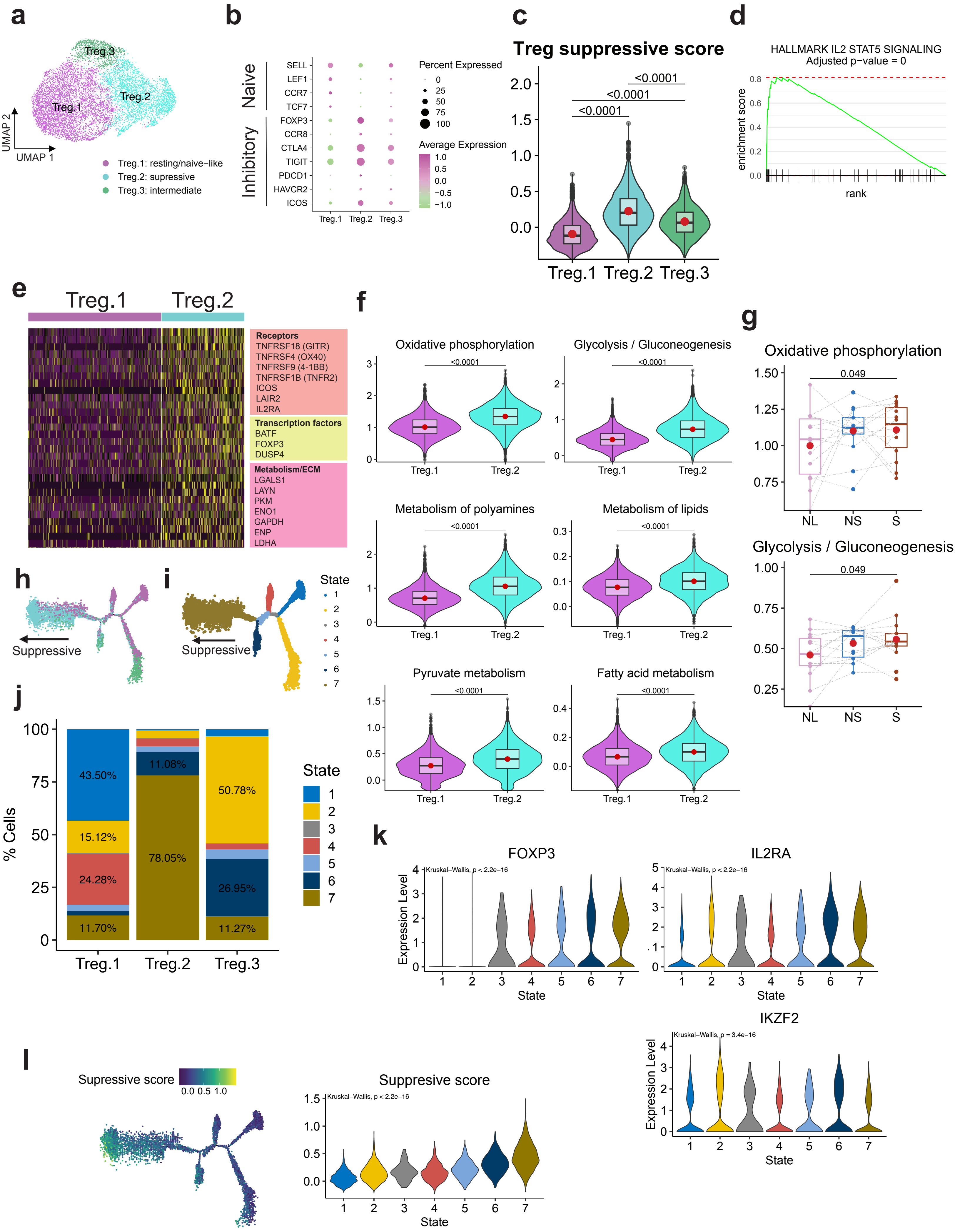
a
Treg.3
Treg.2
Treg.1
UMAP 2
UMAP 1
Treg.1: resting/naive−like
Treg.2: supressive
Treg.3: intermediate
b
Naive
Inhibitory
SELL
LEF1
CCR7
TCF7
FOXP3
CCR8
CTLA4
TIGIT
PDCD1
HAVCR2
ICOS
Treg.1 Treg.2 Treg.3
Percent Expressed
0
25
50
75
100
Average Expression
1.0
0.5
0.0
−0.5
−1.0
c
Treg suppressive score
<0.0001
<0.0001
<0.0001
2.0
1.5
1.0
0.5
0.0
Treg.1 Treg.2 Treg.3
d
HALLMARK IL2 STAT5 SIGNALING
Adjusted p−value = 0
enrichment score
0.8
0.6
0.4
0.2
0.0
rank
e
Treg.1
Treg.2
Receptors
TNFRSF18 (GITR)
TNFRSF4 (OX40)
TNFRSF9 (4-1BB)
TNFRSF1B (TNFR2)
ICOS
LAIR2
IL2RA
Transcription factors
BATF
FOXP3
DUSP4
Metabolism/ECM
LGALS1
LAYN
PKM
ENO1
GAPDH
ENP
LDHA
f
Oxidative phosphorylation
<0.0001
Glycolysis / Gluconeogenesis
<0.0001
Metabolism of polyamines
<0.0001
Metabolism of lipids
<0.0001
Pyruvate metabolism
<0.0001
Fatty acid metabolism
<0.0001
Treg.1 Treg.2
g
Oxidative phosphorylation
0.049
1.50
1.25
1.00
0.75
NL NS S
Glycolysis / Gluconeogenesis
0.049
1.00
0.75
0.50
0.25
NL NS S
h
Suppressive
i
Suppressive
State
1
2
3
4
5
6
7
j
100
75
50
25
0
% Cells
43.50%
15.12%
24.28%
11.70%
11.08%
78.05%
50.78%
26.95%
11.27%
Treg.1 Treg.2 Treg.3
State
1
2
3
4
5
6
7
k
FOXP3
Kruskal−Wallis, p < 2.2e−16
Expression Level
State
IL2RA
Kruskal−Wallis, p < 2.2e−16
State
IKZF2
Kruskal−Wallis, p = 3.4e−16
Expression Level
State
l
Supressive score
0.0 0.5 1.0
Suppresive score
Kruskal−Wallis, p < 2.2e−16
1.5
1.0
0.5
0.0
1 2 3 4 5 6 7
State

**Supplementary Figure 3.7. Characterization of Treg cells and metabolic activities.**

a. UMAP visualization of reclustered Treg cells.
b. Dot plot showing the expression levels of established naïve and inhibitory T cell markers in Treg populations. Expression level is depicted by color, and the size of the dots represents the proportion of cells expressing these markers.
c. Violin and box plots represent the suppressive Treg scores across clusters of Treg cells, with mean values marked by red dots. P-values were calculated using the Wilcoxon signed-rank test.
d. Enrichment plot of the Hallmark pathways 'IL2 STAT5 signaling' up-regulated in the suppressive Tregs (Treg.2) compared to the resting /naïve -like Tregs (Treg.1). Enrichment scores and p-values were obtained using GSEA with an FDR-adjusted p-value reported.
e. Heatmap displaying the relative expression of the top 30 genes with the highest fold-change values that are significantly upregulated (adjusted p-value ≤ 0.05) in suppressive Tregs (Treg.2) compared to resting/naive-like Tregs (Treg.1). Highlighted genes include receptors, transcription factors, and metabolic/ECM genes associated with suppression activity.
f. Comparative analysis of metabolic pathway activity scores between Treg.1 and Treg.2 clusters. The scores were derived using the scMetabolism R package, incorporating KEGG and REACTOM pathways. P-values were determined through the Wilcoxon rank sum test.
g. Activity scores of the 'oxidative phosphorylation' and 'Glycolysis/Gluconeogenesis' pathways of Tregs in adjacent normal tissue (NL), non-solid (NS) and solid (S) components. Scores are averaged per sample. P-values were obtained using the Wilcoxon matched-pairs signed rank test on paired samples from the same patients, with matched samples connected by a line. Mean values are indicated by red dots. Only significant p-values (< 0.05) are shown.
h. Trajectory plot illustrating the inferred progression of Treg cells from resting/naïve-like to suppressive state based on Monocle2 analysis. Cells are color-coded by cluster.
i. Trajectory plot corresponding to panel (h), showing the inferred trajectory of Treg cells. Cells are color-coded according to the trajectory segments, referred to as 'states', as determined by Monocle2 analysis.
j. Percentage of cells across the seven defined trajectory states within each Treg cells subtype.
k. Expression of FOXP3, IL2RA, and IKZF2 across Treg trajectory states. P-values were calculated using the Kruskal-Wallis test.
l. Treg suppressive score along the trajectory corresponding to panel (e) (left) and across the trajectory states (right).

**a**

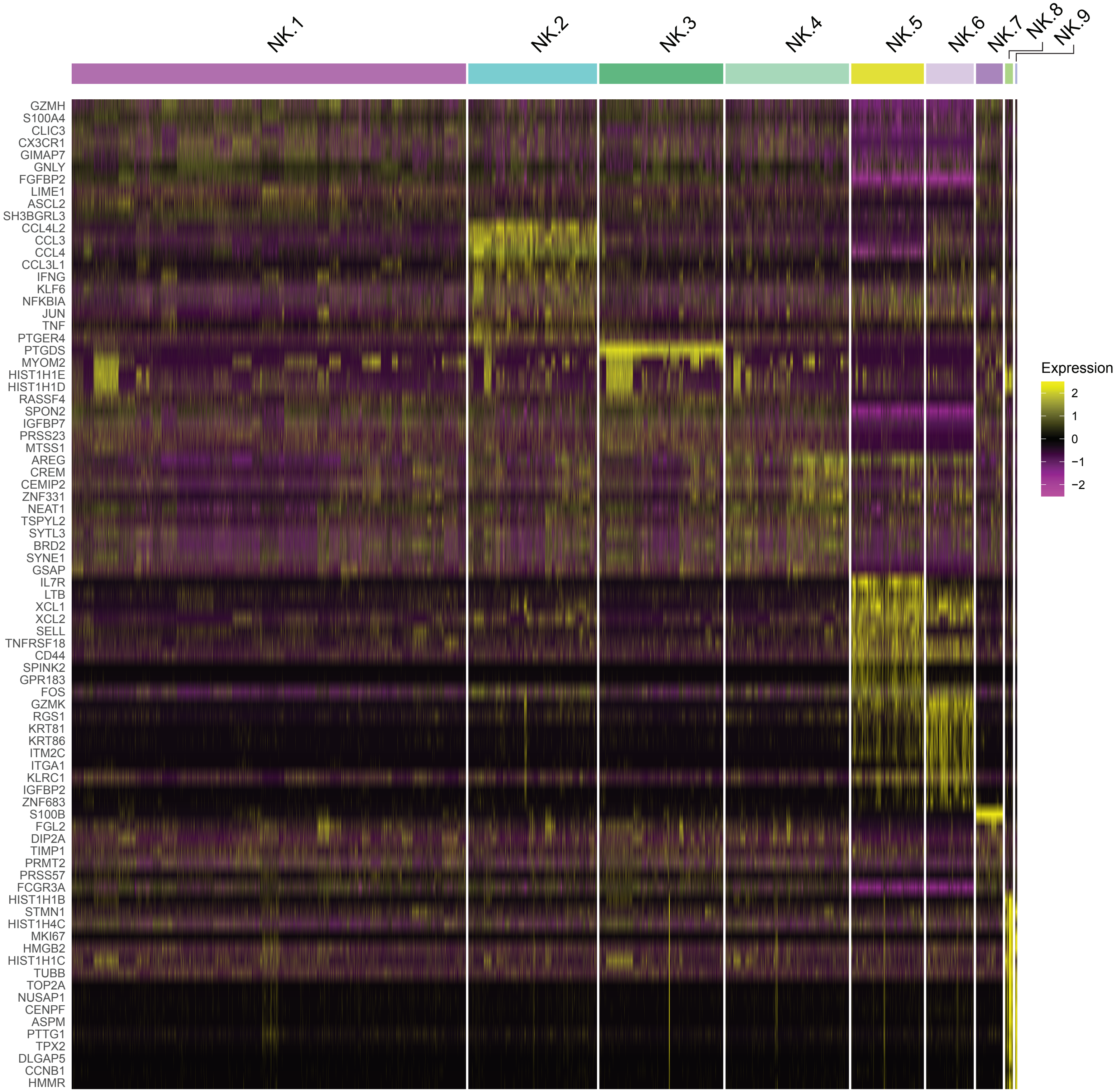


**Supplementary Figure 3.8. Identification of NK cell subpopulations.**

a. Heatmap showing the relative expression of the top ten cluster markers with the highest fold-change values for each NK cluster.

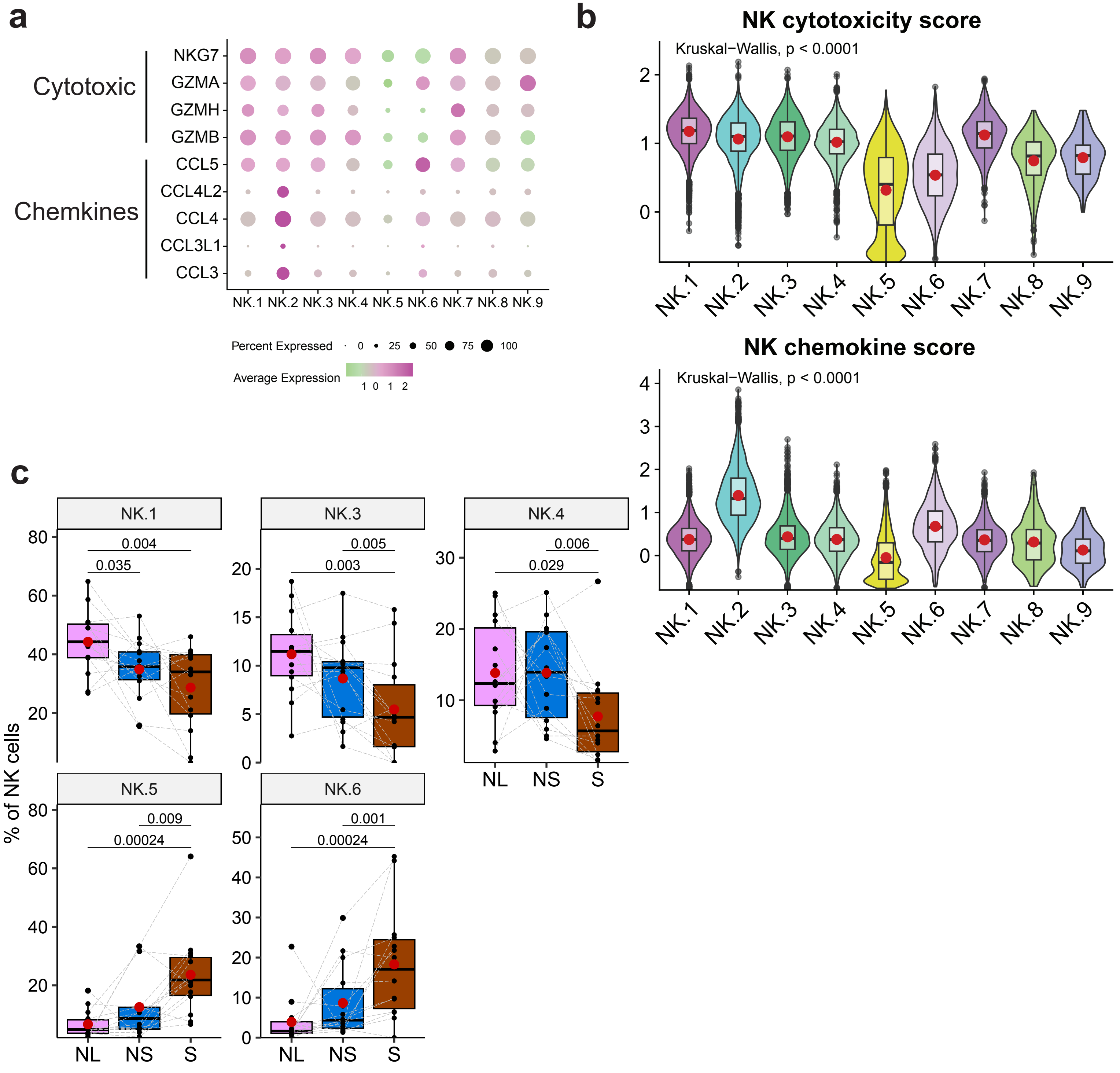


**Supplementary Figure 3.9. Phenotypic characterization of NK cell subpopulations**

a. Dot plot showing the expression levels of cytotoxic markers and chemokines in NK populations. Expression level is depicted by color, and the size of the dots represents the proportion of cells expressing these markers.
b. Violin and box plots represent the cytotoxicity (upper panel) and chemokine expression (lower panel) scores across clusters of NK cells, with mean values marked by red dots. P-values were calculated using the Kruskal-Wallis test.
c. Proportions of NK cell clusters in adjacent normal tissue (NL), non-solid (NS) and solid (S) lesion components. Percentages are based on the entire NK cell population. P-values were obtained using the Wilcoxon matched-pairs signed rank test on paired samples from the same patients, with matched samples connected by a line. Mean values are indicated by red dots. Only significant p-values (< 0.05) are shown.

**Supplementary Fig. 3.10**

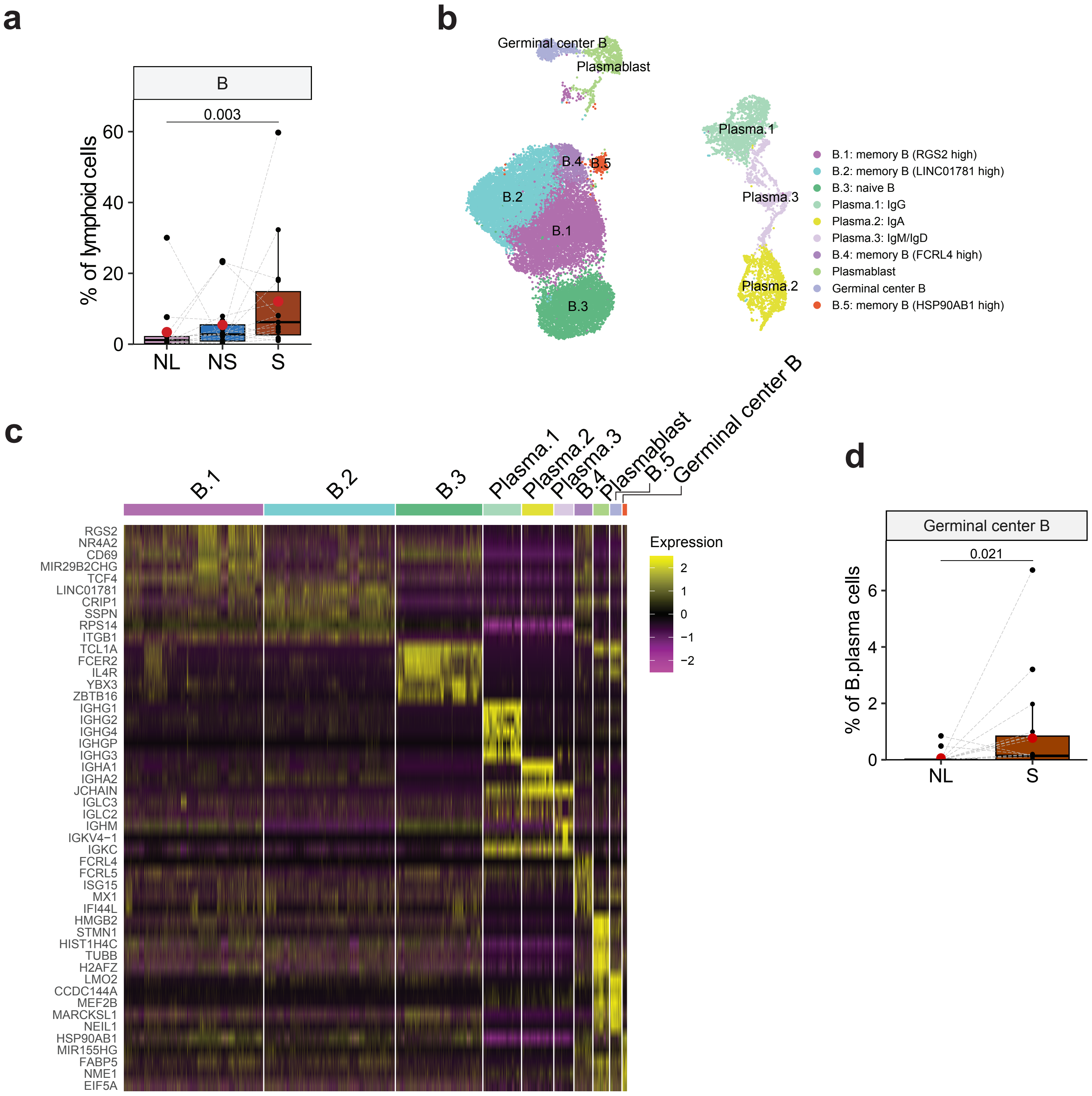


**Supplementary Figure 3.10. Characterization of B and plasma cells.**

a. Proportions of B cells in adjacent normal tissue (NL), non-solid (NS) and solid (S) lesion components. Percentages are based on the entire lymphoid cell population. P-values were obtained using the Wilcoxon matched-pairs signed rank test on paired samples from the same patients, with matched samples connected by a line. Mean values are indicated by red dots. Only significant p-values (< 0.05) are shown.
b. UMAP visualization of reclustered B and plasma cells, illustrating distinct subsets.
c. Heatmap showing the relative expression of the top ten cluster markers with the highest fold-change values for each B/plasma cluster.
d. Proportions of germinal center B cells in adjacent normal tissue (NL) and solid (S) lesion component. Percentages are based on the entire B/plasma cell population. P-values were obtained using the Wilcoxon matched-pairs signed rank test on paired samples from the same patients, with matched samples connected by a line. Mean values are indicated by red dots. Only significant p-values (< 0.05) are shown.

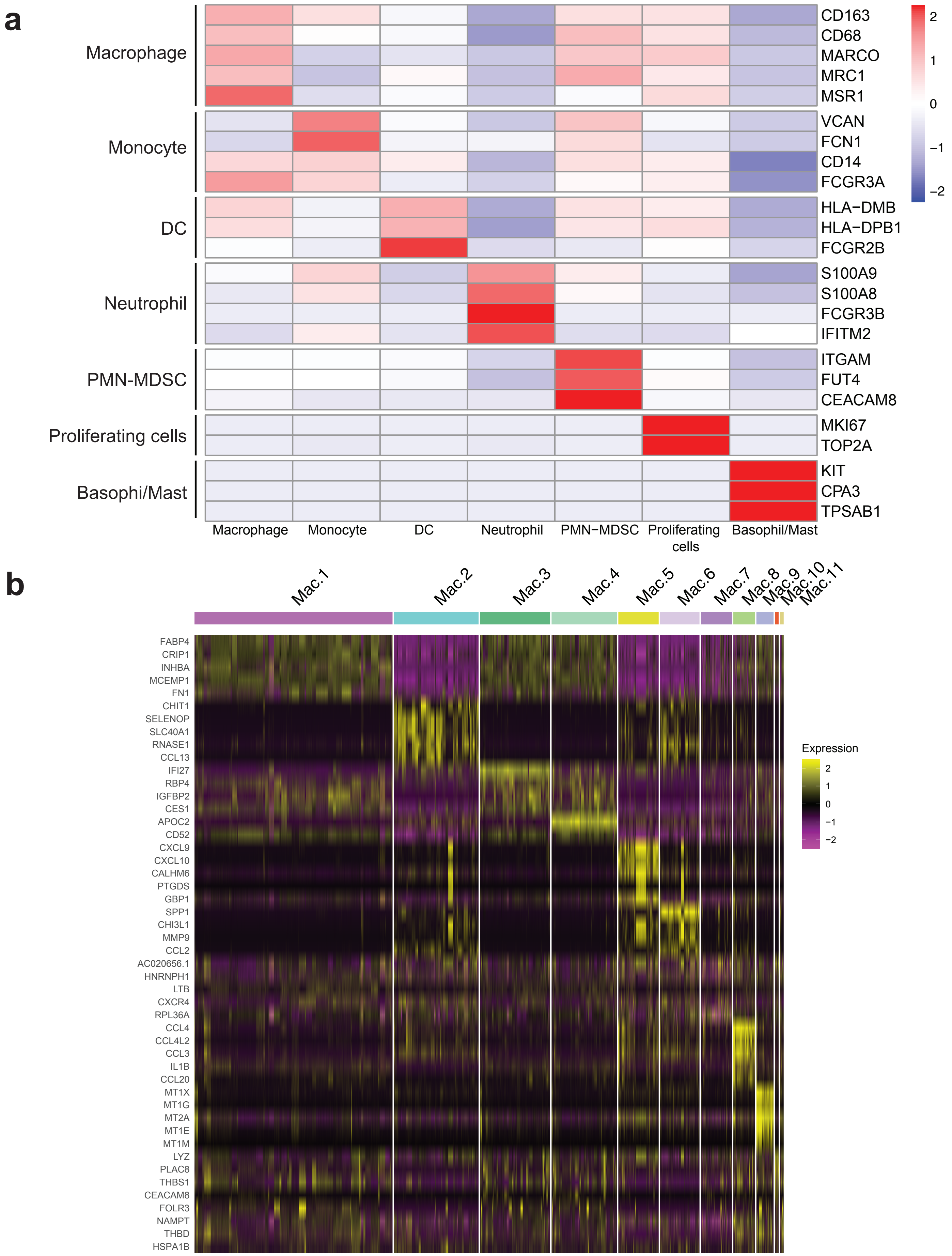


**Supplementary Figure 4.1. Expression of canonical markers in myeloid cell types and expression of macrophage cluster markers.**

a. Heatmaps showing mean expression levels of established markers for myeloid cell types within clusters of myeloid cells.

b. Heatmap showing the relative expression of the top five cluster markers with the highest fold-change values for each macrophage cluster.

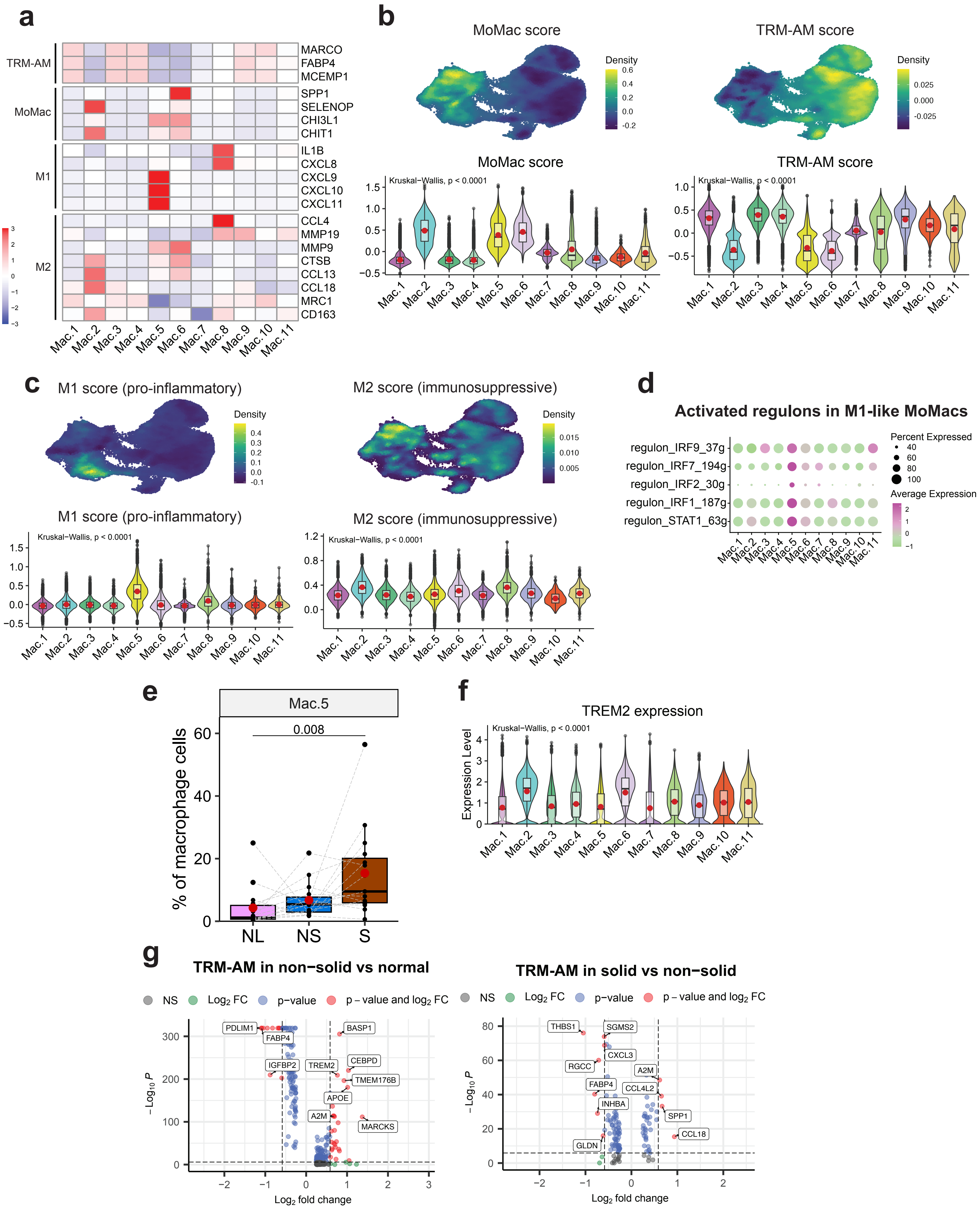
a
TRM-AM
MoMac
M1
M2
MARCO
FABP4
MCEMP1
SPP1
SELENOP
CHI3L1
CHIT1
IL1B
CXCL8
CXCL9
CXCL10
CXCL11
CCL4
MMP19
MMP9
CTSB
CCL13
CCL18
MRC1
CD163
Mac.1 Mac.2 Mac.3 Mac.4 Mac.5 Mac.6 Mac.7 Mac.8 Mac.9 Mac.10 Mac.11
b
MoMac score
TRM-AM score
Density
Kruskal−Wallis, p < 0.0001
c
M1 score (pro-inflammatory)
M2 score (immunosuppressive)
Density
Kruskal−Wallis, p < 0.0001
d
Activated regulons in M1-like MoMacs
regulon_IRF9_37g
regulon_IRF7_194g
regulon_IRF2_30g
regulon_IRF1_187g
regulon_STAT1_63g
Percent Expressed
Average Expression
e
Mac.5
0.008
% of macrophage cells
NL NS S
f
TREM2 expression
Kruskal−Wallis, p < 0.0001
Expression Level
g
TRM-AM in non−solid vs normal
TRM-AM in solid vs non−solid
NS
Log2 FC
p−value
p – value and log2 FC
PDLIM1
FABP4
IGFBP2
TREM2
BASP1
CEBPD
TMEM176B
APOE
A2M
MARCKS
THBS1
SGMS2
CXCL3
RGCC
A2M
FABP4
CCL4L2
INHBA
SPP1
CCL18
GLDN
− Log10 P
Log2 fold change

**Supplementary Figure 4.2. Characterization of macrophage populations.**

a. Heatmaps showing mean expression levels of established markers for TRM-AM, MoMac, M1 and M2 macrophages across macrophage clusters.
b. MoMac (left) and TRM-AM (right) scores of myeloid cells are presented in density plots (upper panel) and violin and box plots (lower panel). Mean values are indicated by red dots. P-values were calculated using the Kruskal-Wallis test.
c. M2 (left) and M1 (right) scores of myeloid cells are presented in density plots (upper panel) and violin and box plots (lower panel). Mean values are indicated by red dots. P-values were calculated using the Kruskal-Wallis test.
d. Dot plot showing the expression levels of regulon activity in macrophage populations. Expression level is depicted by color, and the size of the dots represents the proportion of cells expressing the genes of the regulons.
e. Proportions of M1-like MomMac cluster, Mac.5, in adjacent normal tissue (NL) and solid (S) lesion component. Percentages are based on the entire macrophage population. P-values were obtained using the Wilcoxon matched-pairs signed rank test on paired samples from the same patients, with matched samples connected by a line. Mean values are indicated by red dots. Only significant p-values ($< 0.05$) are shown.
f. TREM2 expression across macrophage clusters. Mean values are indicated by red dots. P-value was calculated using the Kruskal-Wallis test.
g. Volcano plot comparing gene expression in TRM-AMs in non-solid vs. normal (upper panel) and in solid vs. non-solid (lower panel). The x-axis represents the log2 fold change in gene expression, with positive values indicating higher expression in non-solid (upper panel) and solid (lower panel) samples and negative values indicating higher expression in normal (upper panel) and non-solid (lower panel) samples. The y-axis represents the -log10 of the p-value, with higher values indicating greater statistical significance. Green dots indicate genes with significant fold changes ($\geq 1.5$), blue dots indicate genes with significant adjusted p-values ($\leq 0.05$), and red dots indicate genes significant in both fold change and p-value.

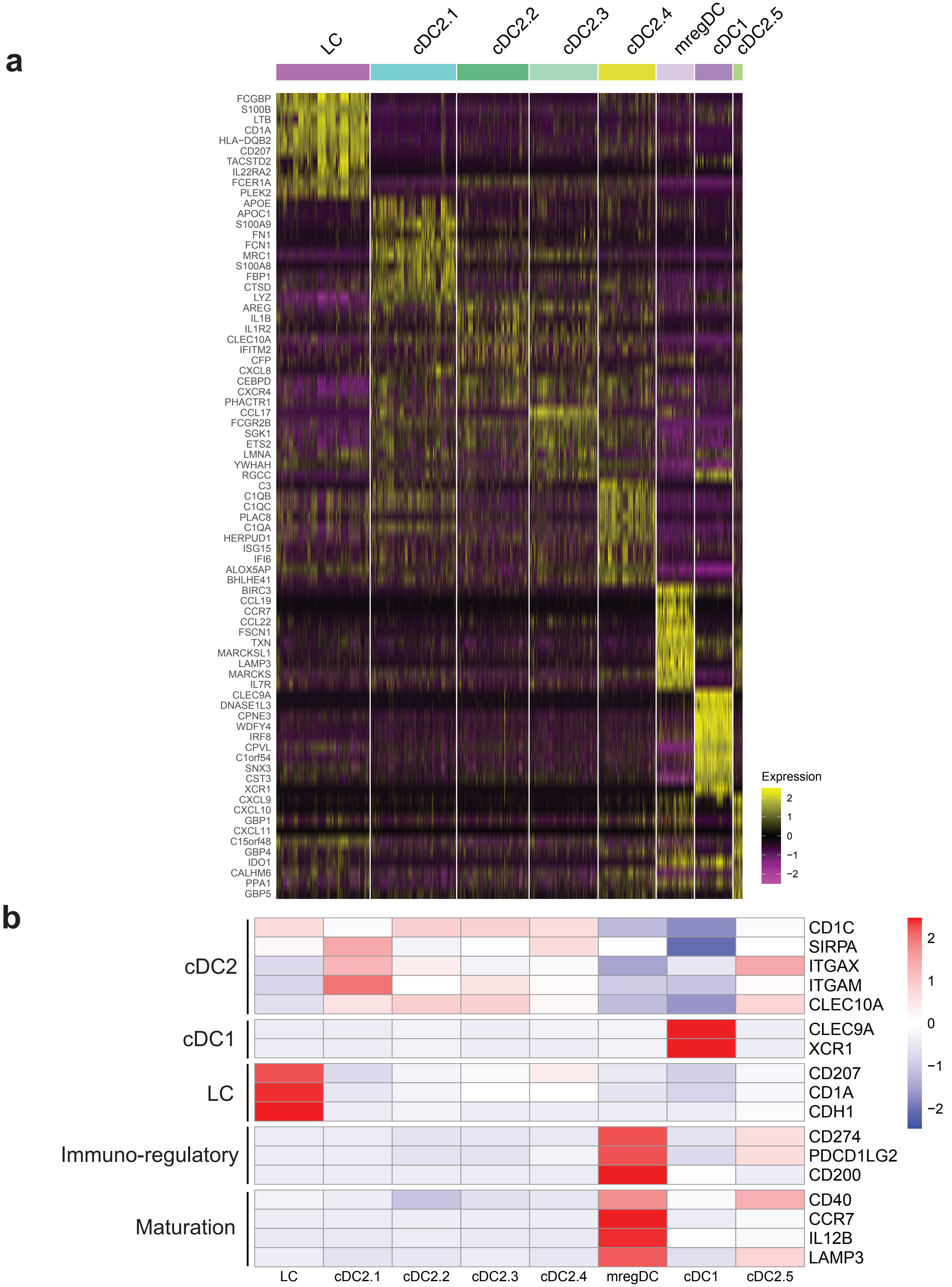


**Supplementary Figure 4.3. Characterization of major DC populations.**

a. Heatmap showing the relative expression of the top ten cluster markers with the highest fold-change values for each DC cluster.
b. Heatmaps showing mean expression levels of established markers for DC cell types within DC clusters.

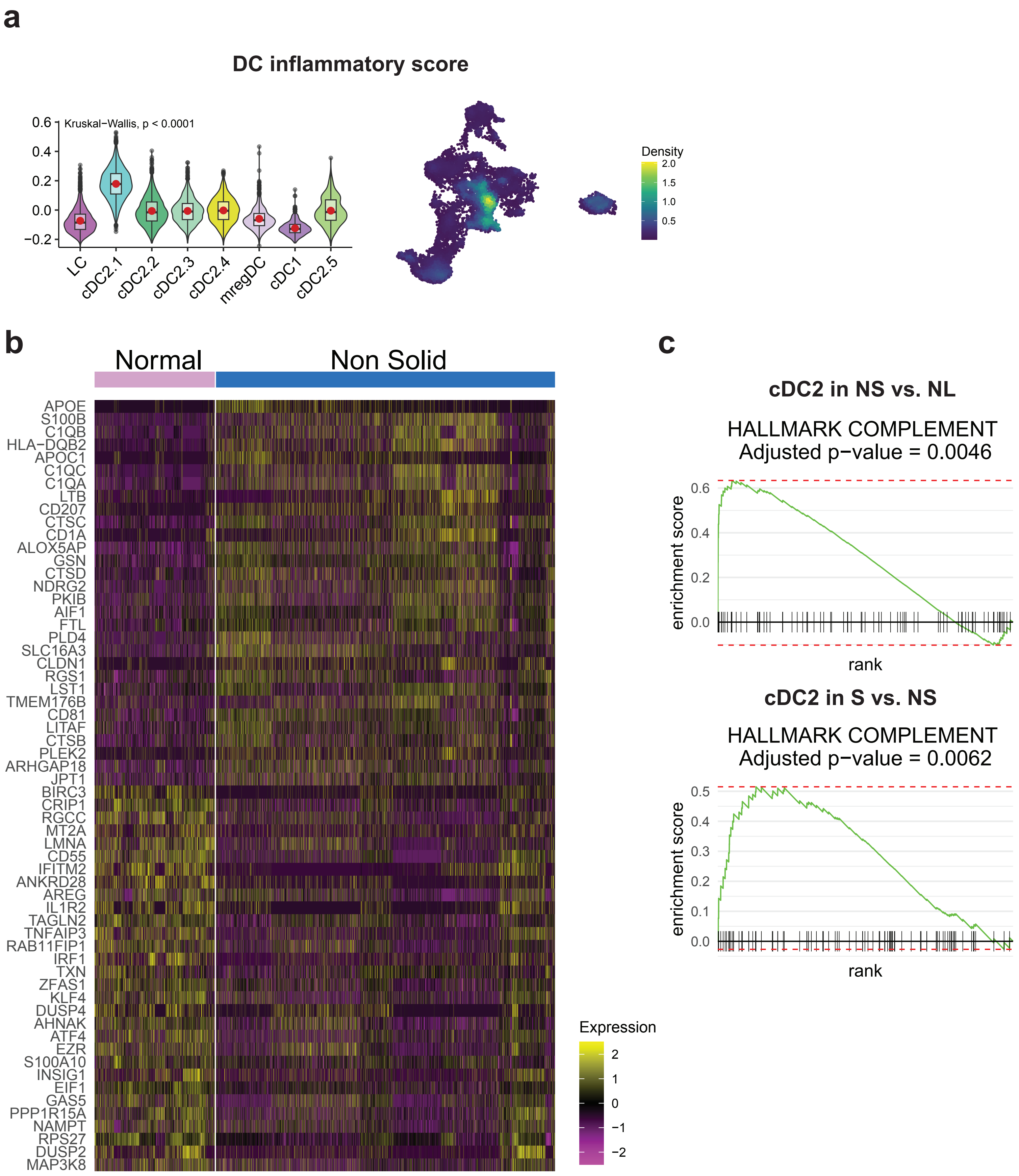


**Supplementary Figure 4.4. Phenotypic characterization of major DC populations.**

a. Inflammatory scores of DCs are presented in violin and box plots (left) and density plots (right). Mean values are indicated by red dots. P-value was calculated using the Kruskal-Wallis test.
b. Heatmap showing the relative expression of the top 30 up and down-regulated genes of cDC2 cells between the non-solid component and adjacent normal tissue.
c. Enrichment plot of the Hallmark pathways ‘complement’ that was up-regulated in cDC2 cells in the non-solid (NS) component compared to adjacent normal tissue (NL; upper panel) and in the solid (S) compared to the non-solid (NS; lower panel). Enrichment scores and p-values were obtained using GSEA, with an FDR-adjusted p-value shown.

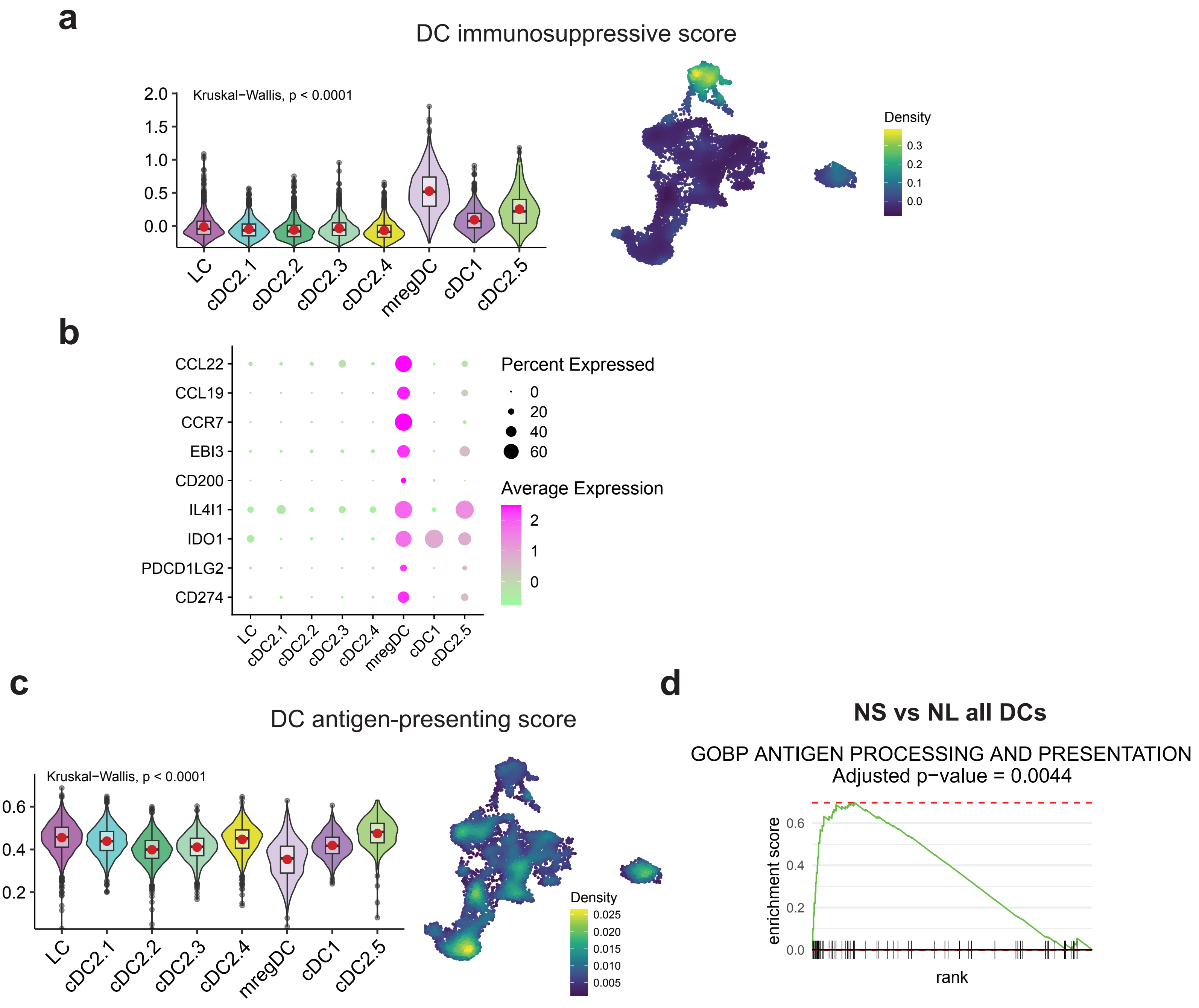


**Supplementary Figure 4.5. Phenotypic characterization of DC populations in non-solid and solid components of part-solid nodules.**

a. Immunosuppressive scores of DCs are presented in violin and box plots (left) and density plots (right). Mean values are indicated by red dots. P-value was calculated using the Kruskal-Wallis test.
b. Dot plot showing the expression levels of mregDC markers in DC populations. Expression level is depicted by color, and the size of the dots represents the proportion of cells expressing the genes of the regulons.
c. (Antigen-presenting scores of DCs are presented in violin and box plots (left) and density plots (right). Mean values are indicated by red dots. P-value was calculated using the Kruskal-Wallis test.
d. Enrichment plot of the GO BP pathways ‘antigen processing and presentation’ that was up-regulated in DCs in the non-solid (NS) component compared to adjacent normal tissue (NL). Enrichment score and p-value were obtained using GSEA, with an FDR-adjusted p-value shown.

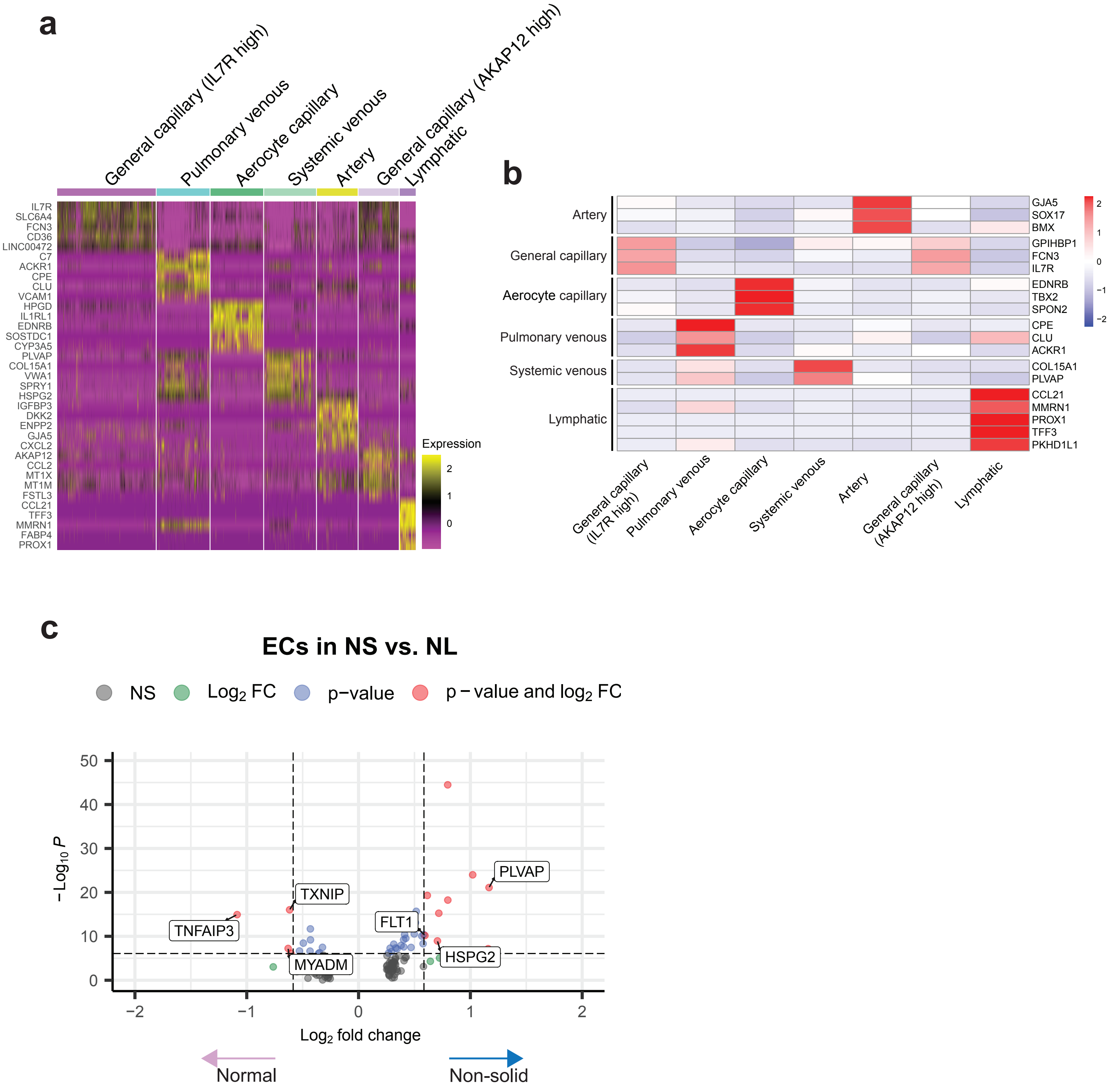


**Supplementary Figure 5.1. Transcriptomic characterization of EC clusters.**

a. Heatmap showing the relative expression of the top five cluster markers with the highest fold-change values for each EC cluster.
b. Heatmaps showing mean expression levels of established markers for EC types within the identified clusters of ECs.
c. Volcano plot comparing gene expression in ECs in non-solid (NS) vs. normal (NL). The x-axis represents the log2 fold change in gene expression, with positive values indicating higher expression in NS samples and negative values indicating higher expression in NL samples. The y-axis represents the -log10 of the p-value, with higher values indicating greater statistical significance. Green dots indicate genes with significant fold changes (≥ 1.5), blue dots indicate genes with significant adjusted p-values (≤ 0.05), and red dots indicate genes significant in both fold change and p-value.

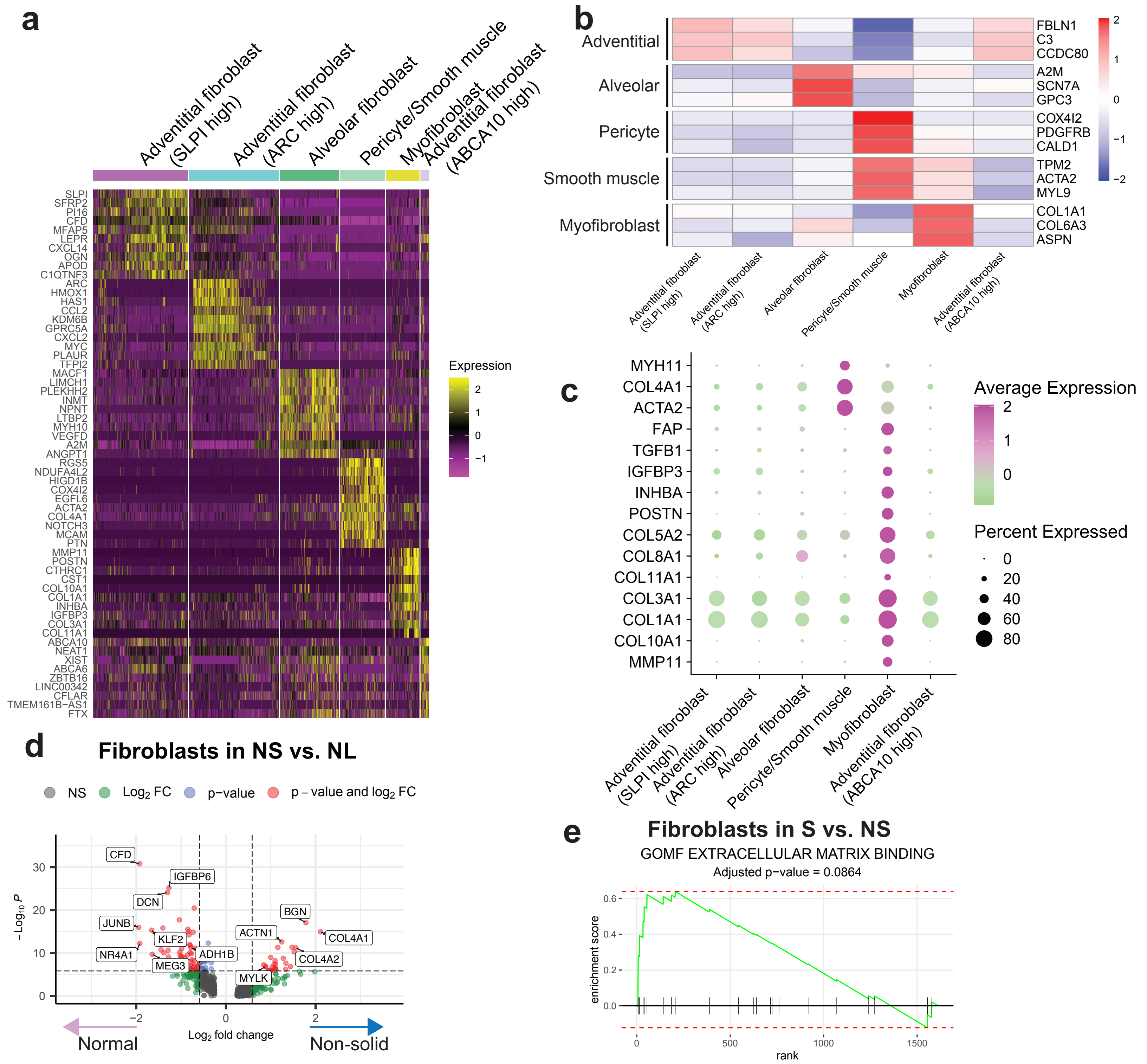


**Supplementary Figure 5.2. Transcriptomic characterization of fibroblast populations**

a. Heatmap showing the relative expression of the top five cluster markers with the highest fold-change values for each fibroblast cluster.
b. Heatmaps showing mean expression levels of established markers for fibroblast types within the identified clusters of ECs.
c. Dot plot showing the expression levels of genes in the fibroblast subtypes. Expression level is depicted by color, and the size of the dots represents the proportion of cells expressing these markers.
d. Volcano plot comparing gene expression between fibroblasts in non-solid (NS) component vs. adjacent normal lung tissue (NL). The x-axis represents the log2 fold change in gene expression, with positive values indicating higher expression in solid samples and negative values indicating higher expression in non-solid samples. The y-axis represents the -log10 of the p-value, with higher values indicating greater statistical significance. Green dots indicate genes with significant fold changes (≥ 1.5), blue dots indicate genes with significant adjusted p-values (≤ 0.05), and red dots indicate genes significant in both fold change and p-value. Upregulated epithelial-mesenchymal transition markers referenced in the text are indicated.
e. Enrichment plots of the up-regulated ‘extracellular matrix binding’ pathway in fibroblasts in the solid (S) compared to non-solid (NS). Enrichment scores and p-values were obtained using GSEA, with an FDR-adjusted p-value shown.

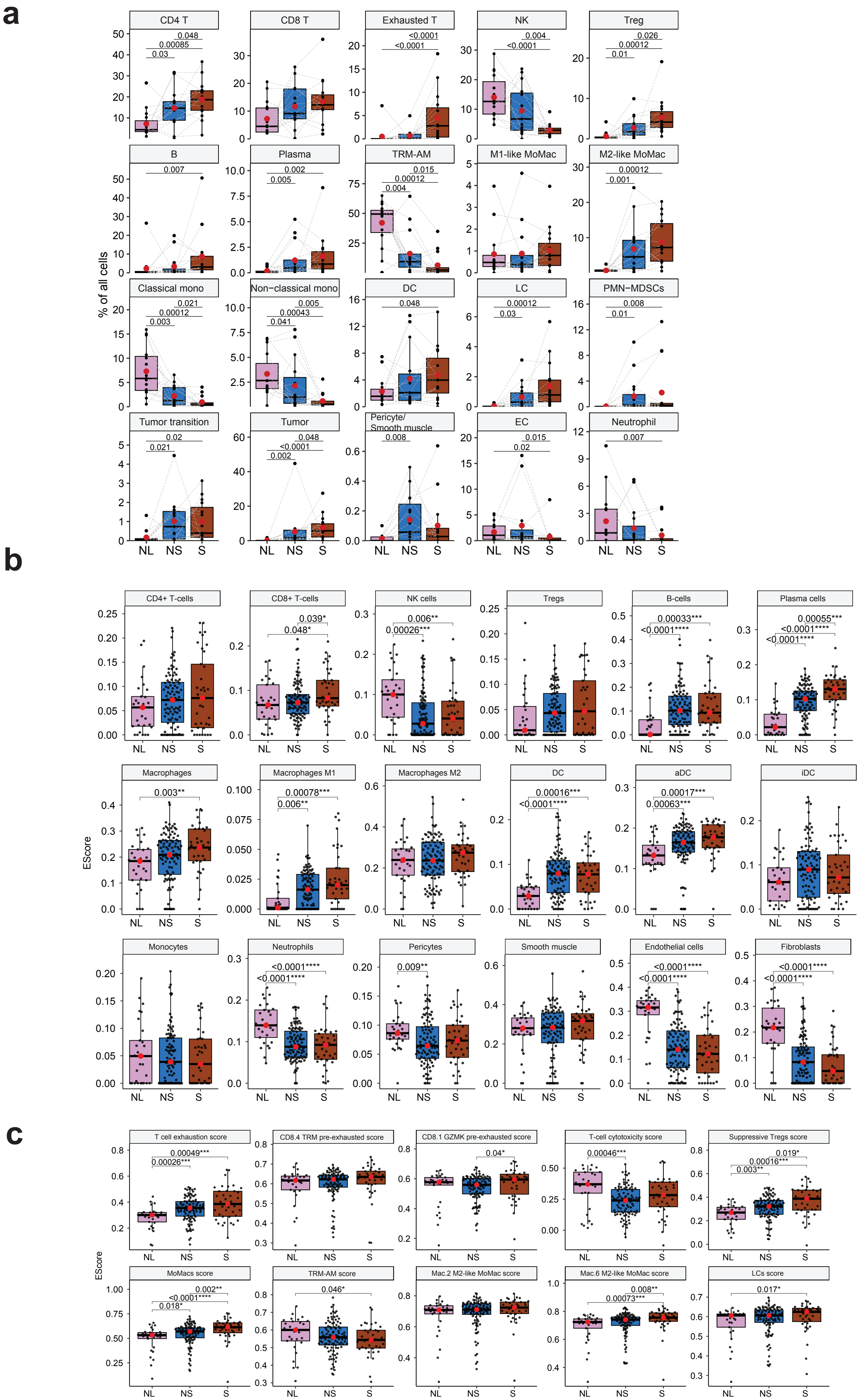
a
% of all cells
CD4 T
0.048
0.00085
0.03
CD8 T
Exhausted T
<0.0001
<0.0001
NK
0.004
<0.0001
Treg
0.026
0.00012
0.01
B
0.007
Plasma
0.002
0.005
TRM-AM
0.015
0.00012
0.004
M1-like MoMac
M2-like MoMac
0.00012
0.001
Classical mono
0.021
0.00012
0.003
Non-classical mono
0.005
0.00043
0.041
DC
0.048
LC
0.00012
0.03
PMN−MDSCs
0.008
0.01
Tumor transition
0.02
0.021
Tumor
0.048
<0.0001
0.002
Pericyte/
Smooth muscle
0.008
EC
0.015
0.02
Neutrophil
0.007
NL
NS
S
b
EScore
CD4+ T-cells
CD8+ T-cells
0.039*
0.048*
NK cells
0.006**
0.00026***
Tregs
B-cells
0.00033***
<0.0001****
Plasma cells
0.00055***
<0.0001****
<0.0001****
Macrophages
0.003**
Macrophages M1
0.00078***
0.006**
Macrophages M2
DC
0.00016***
<0.0001****
aDC
0.00017***
0.00063***
iDC
Monocytes
Neutrophils
<0.0001****
<0.0001****
Pericytes
0.009**
Smooth muscle
Endothelial cells
<0.0001****
<0.0001****
Fibroblasts
<0.0001****
<0.0001****
NL
NS
S
c
EScore
T cell exhaustion score
0.00049***
0.00026***
CD8.4 TRM pre-exhausted score
CD8.1 GZMK pre-exhausted score
0.04*
T-cell cytotoxicity score
0.00046***
Suppressive Tregs score
0.019*
0.00016***
0.003**
MoMacs score
0.002**
<0.0001****
0.018*
TRM-AM score
0.046*
Mac.2 M2-like MoMac score
Mac.6 M2-like MoMac score
0.008**
0.00073***
LCs score
0.017*
NL
NS
S

**Supplementary Figure 5.3. Enumeration of cell abundance and phenotypes in an independent cohort of patients with pure non-solid and solid nodules.**

a. Proportions of major cell populations in adjacent normal tissue (NL), non-solid (NS) and solid (S) lesion components. Percentages are based on all annotated cells. P-values were obtained using the Wilcoxon matched-pairs signed rank test on paired samples from the same patients, with matched samples connected by a line. Mean values are indicated by red dots. Only significant p-values (< 0.05) are shown.
b. Comparison of deconvolution abundance scores estimated by xCell between adjacent normal tissue (NL), non-solid (NS), and solid (S) lesion components. P-values were calculated using the Wilcoxon rank sum test. Median values are indicated by red dots. Only significant p-values (< 0.05) are shown.
c. Comparison of ssGSEA scores between adjacent normal tissue (NL), non-solid (NS) and solid (S) lesion components. The scores were calculated based on gene signatures from the scRNA-seq analysis as detailed in Supplementary Table S15. P-values were obtained using the Wilcoxon rank sum. Median values are marked by red dots. Only significant p-values (< 0.05) are shown.

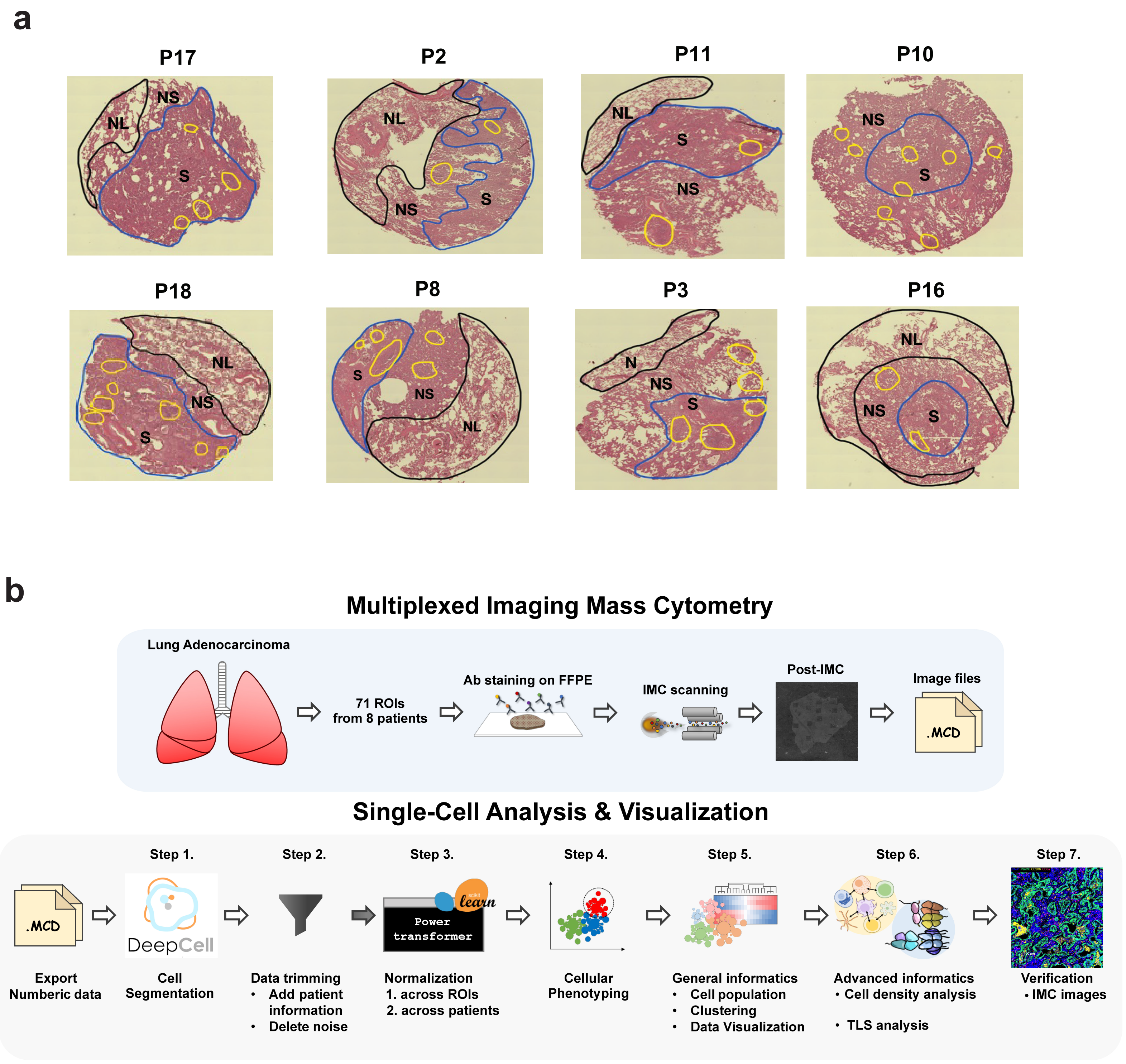


**Supplementary Figure 7.1. Pathologist annotated H&E images of part-solid nodules and a schematic of IMC workflow**.

a. H&E images of patients with part-solid lesions used for spatial analysis. Pathologist-annotated normal lung (NL), non-solid (NS) and solid (S) components are depicted. Tertiary lymphoid structures (yellow circles)
b. Schematic representation of the IMC analysis workflow.

### Supplementary Fig. 7.2

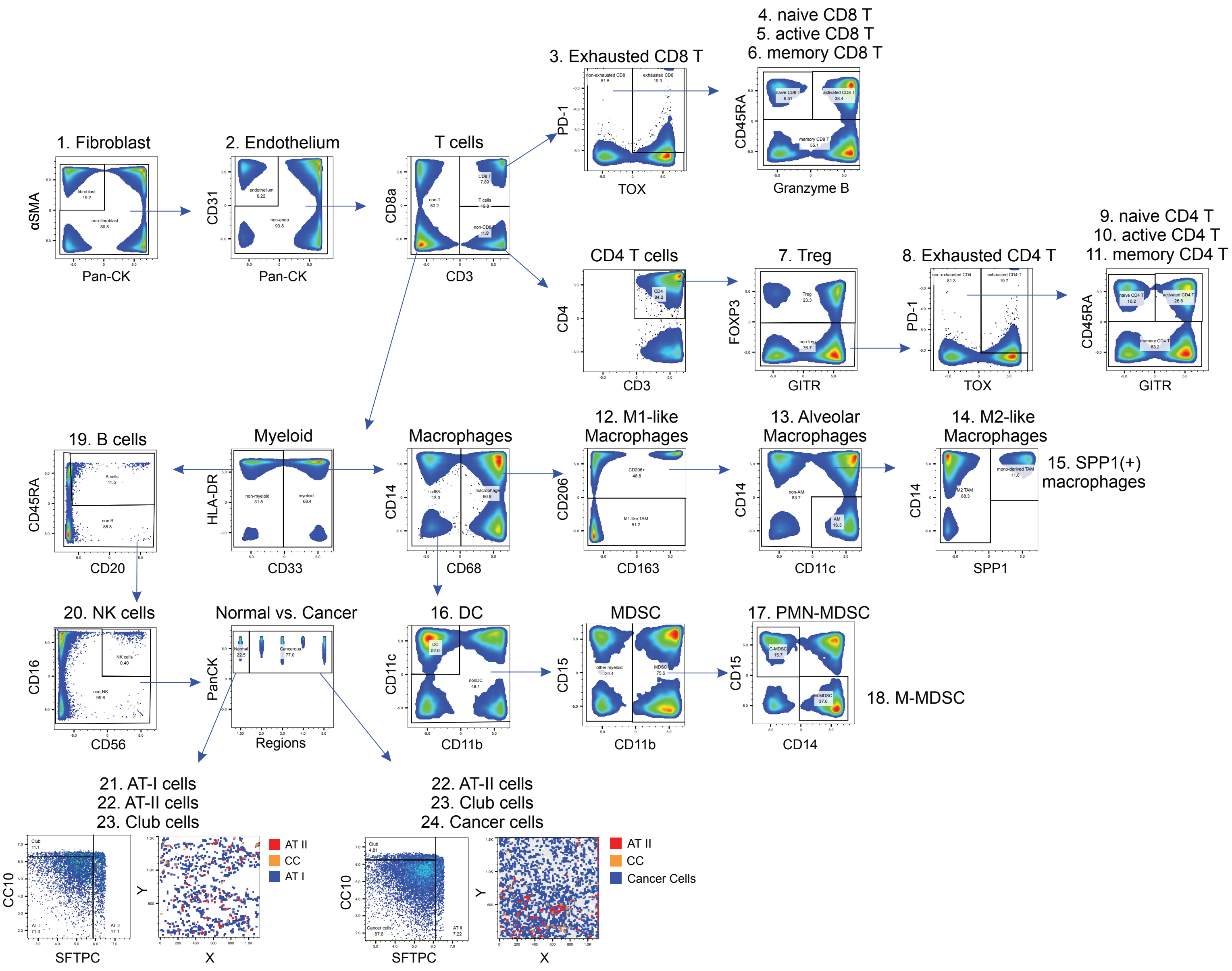


**Supplementary Figure 7.2. Gating strategies for the identification of 24 cell phenotypes.**

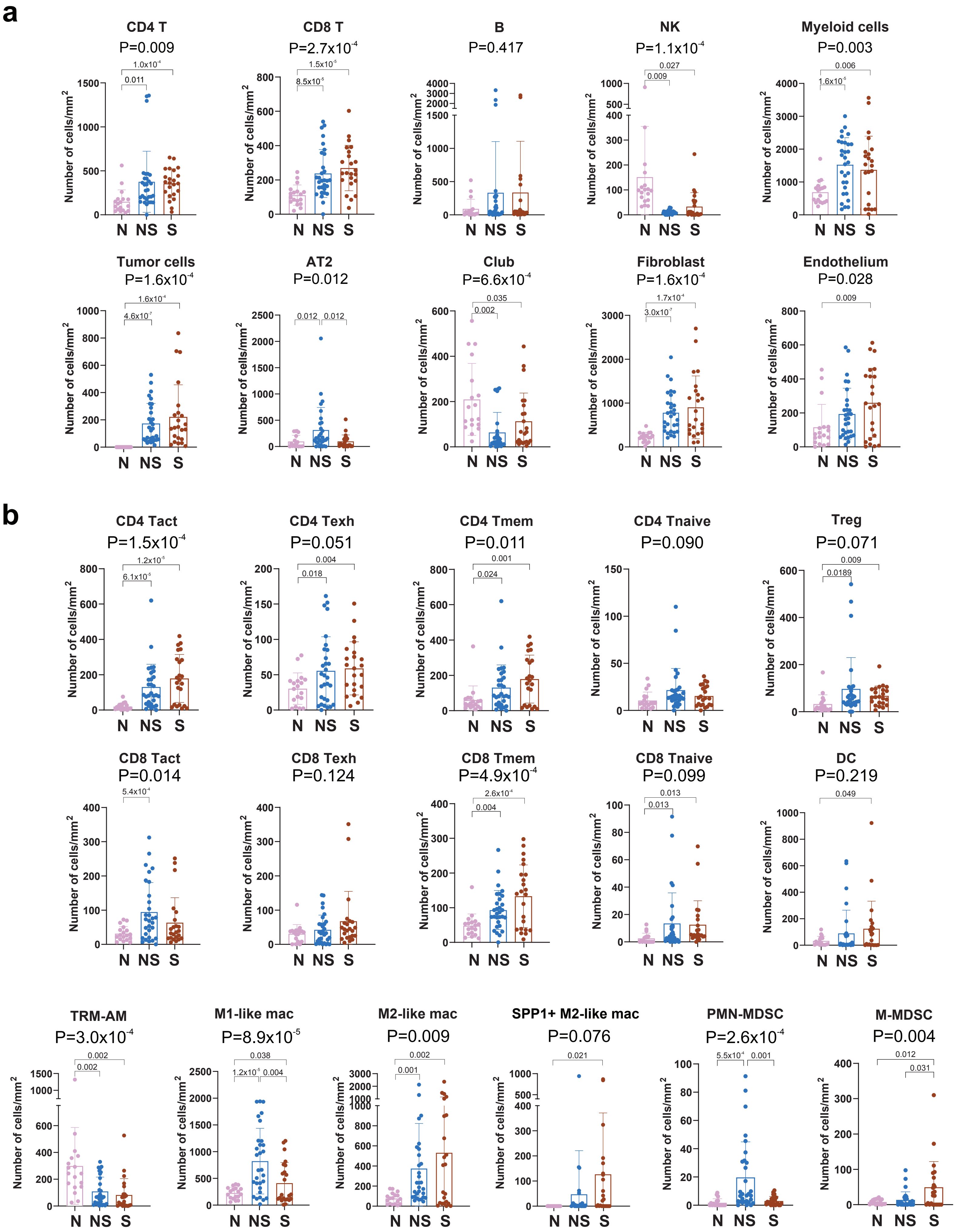

a
CD4 T P=0.009
CD8 T P=2.7x10-4
B P=0.417
NK P=1.1x10-4
Myeloid cells P=0.003
Tumor cells P=1.6x10-4
AT2 P=0.012
Club P=6.6x10-4
Fibroblast P=1.6x10-4
Endothelium P=0.028
b
CD4 Tact P=1.5x10-4
CD4 Texh P=0.051
CD4 Tmem P=0.011
CD4 Tnaive P=0.090
Treg P=0.071
CD8 Tact P=0.014
CD8 Texh P=0.124
CD8 Tmem P=4.9x10-4
CD8 Tnaive P=0.099
DC P=0.219
TRM-AM P=3.0x10-4
M1-like mac P=8.9x10-5
M2-like mac P=0.009
SPP1+ M2-like mac P=0.076
PMN-MDSC P=2.6x10-4
M-MDSC P=0.004
Number of cells/mm2
N NS S

**Supplementary Figure 7.3. Enumeration of major cellular populations in IMC.**

a. Enumeration of major cell populations (number of cells/mm¬2) in normal lung (N), non-solid (NS) and solid (S) regions. Only significant p-values (< 0.05) are shown.
b. Enumeration of immune cell functional phenotypes (number of cells/mm¬2) normal lung (N), non-solid (NS) and solid (S) regions. Only significant p-values (< 0.05) are shown.

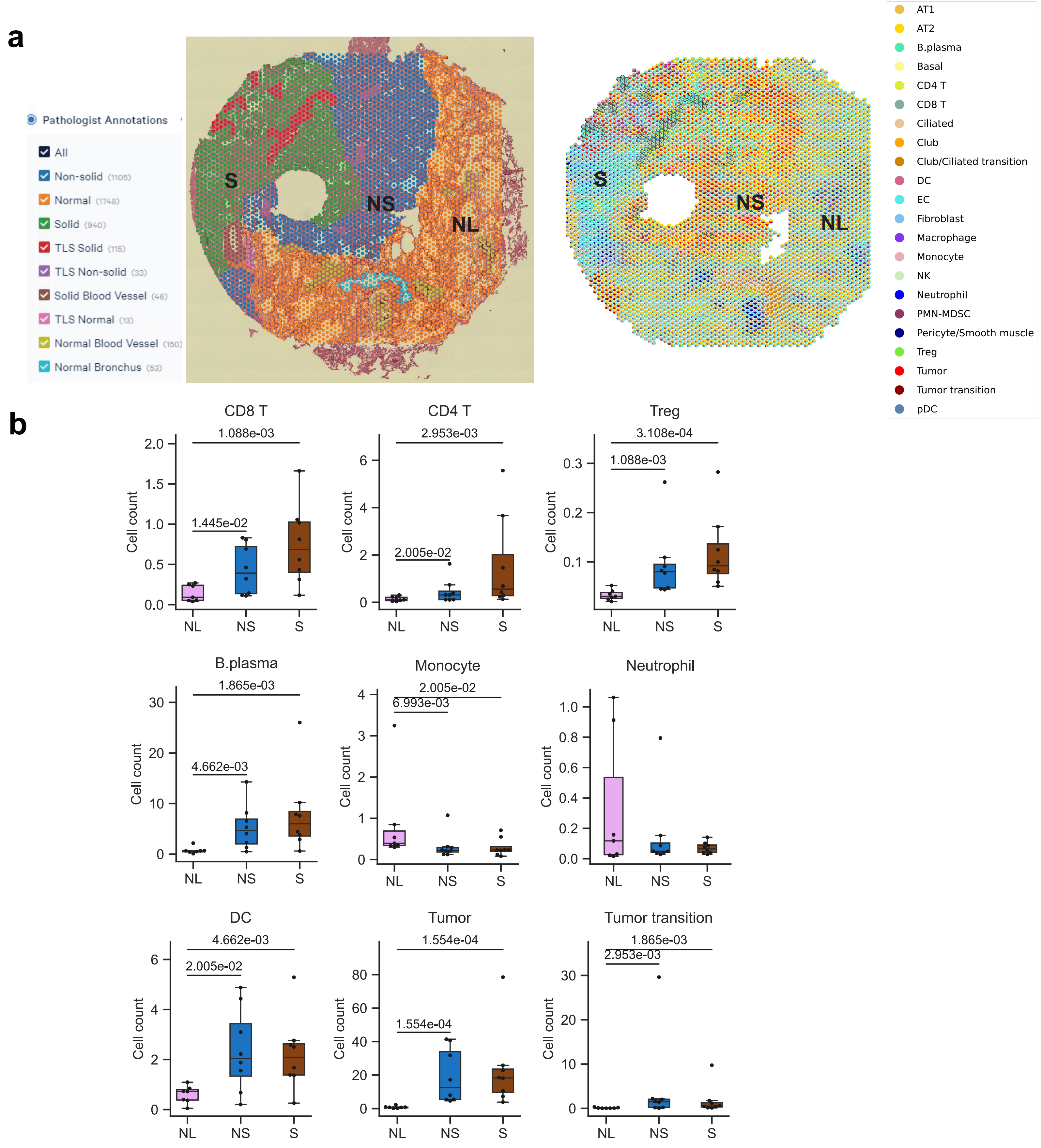


**Supplementary Figure 7.4: Spatial transcriptomics (10X Visium) analysis of part-solid nodules.**

a. Pathologist annotations (left) and cell type deconvolution (right) of spatial transcriptomics (10x Visium) spots of a representative patient (P18).
b. Abundance of major cell types obtained from spatial transcriptomic analysis of samples from normal lung (N), non-solid (NS) and solid (S) regions. Cell counts were assessed using deconvolution based on scRNA-seq data. P-values were calculated using the Mann-Whitney U test. Only significant p-values (< 0.05) are shown.

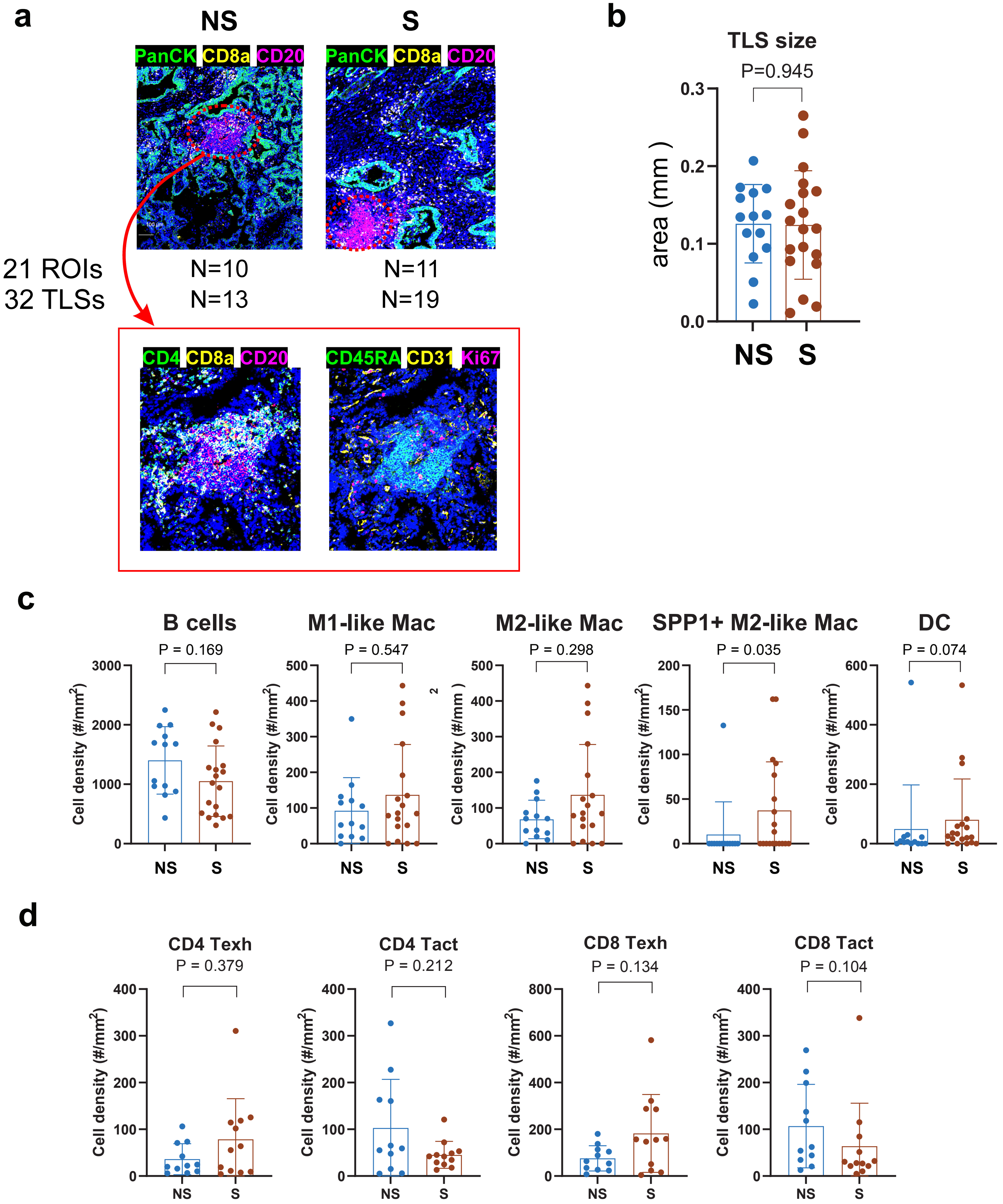


**Supplementary Figure 7.5: IMC analysis of tertiary lymph node structures (TLS).**

a. Representative multichannel IMC images of TLS (21 ROIs with 32 TLS) in non-solid (NS) and solid (S) components.
b. TLS size in mm2 in non-solid (NS) and solid (S) components.
c. Cell densities (per mm2) of specific immune populations in the TLS of non-solid (NS) and solid (S) components
d. Cell densities (per mm2) of specific immune populations outside the TLS in non-solid (NS) and solid (S) components.

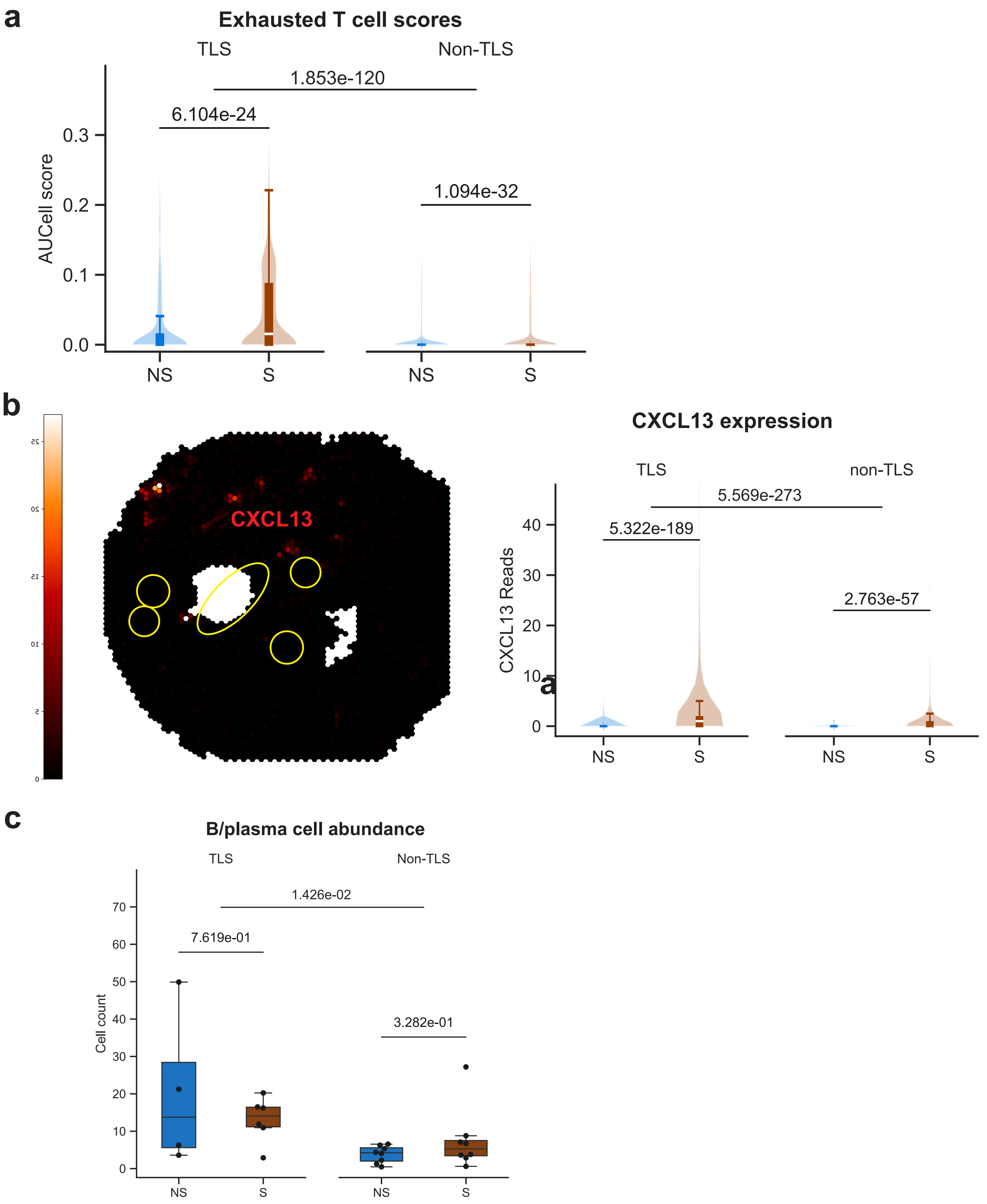


**Supplementary Figure 7.6. Spatial transcriptomics (10X Visium) analysis of TLSs**

a. exhausted T cell enrichment score in TLS and non-TLS spots in non-solid (NS) and solid (S) components. P-values were calculated using the Mann-Whitney U test. Only significant p-values (< 0.05) are shown.
b. ST analysis showing CXCL13 expression is confined to TLS (yellow circles, left panel) and CXCL13 expression as measured in TLS and non-TLS spots in non-solid (NS) and solid (S) components (right panel).
c. B cell abundance in TLS and non-TLS spots in non-solid (NS) and solid (S) components
